# An enhancer-AAV toolbox to target and manipulate distinct interneuron subtypes

**DOI:** 10.1101/2024.07.17.603924

**Authors:** Elisabetta Furlanis, Min Dai, Brenda Leyva Garcia, Thien Tran, Josselyn Vergara, Ana Pereira, Bram L. Gorissen, Sara Wills, Anna Vlachos, Ariel Hairston, Deepanjali Dwivedi, Sarah Du, Justin McMahon, Shuhan Huang, Annunziato Morabito, Arenski Vazquez, Soyoun Kim, Anthony T. Lee, Edward F. Chang, Taha Razzaq, Ahmed Qazi, Geoffrey Vargish, Xiaoqing Yuan, Adam Caccavano, Steven Hunt, Ramesh Chittajallu, Nadiya McLean, Lauren Hewitt, Emily Paranzino, Haley Rice, Alex C. Cummins, Anya Plotnikova, Arya Mohanty, Anne Claire Tangen, Jung Hoon Shin, Reza Azadi, Mark A.G. Eldridge, Veronica A. Alvarez, Bruno B. Averbeck, Mansour Alyahyay, Tania Reyes Vallejo, Mohammed Soheib, Lucas G. Vattino, Cathryn P. MacGregor, Carolina Piletti Chatain, Emmie Banks, Viktor Janos Olah, Shovan Naskar, Sophie Hill, Sophie Liebergall, Rohan Badiani, Lili Hyde, Ella Hanley, Qing Xu, Kathryn C. Allaway, Ethan M. Goldberg, Matthew J.M. Rowan, Tomasz J. Nowakowski, Soohyun Lee, Emilia Favuzzi, Pascal S. Kaeser, Lucas Sjulson, Renata Batista-Brito, Anne E. Takesian, Leena A. Ibrahim, Asim Iqbal, Kenneth A. Pelkey, Chris J. McBain, Jordane Dimidschstein, Gord Fishell, Yating Wang

## Abstract

In recent years, we and others have identified a number of enhancers that, when incorporated into rAAV vectors, can restrict the transgene expression to particular neuronal populations. Yet, viral tools to access and manipulate specific neuronal subtypes are still limited. Here, we performed systematic analysis of single cell genomic data to identify enhancer candidates for each of the telencephalic interneuron subtypes. We established a set of enhancer-AAV tools that are highly specific for distinct cortical interneuron populations and striatal cholinergic interneurons. These enhancers, when used in the context of different effectors, can target (fluorescent proteins), observe activity (GCaMP) and manipulate (opto-genetics) specific neuronal subtypes. We also validated our enhancer-AAV tools across species. Thus, we provide the field with a powerful set of tools to study neural circuits and functions and to develop precise and targeted therapy.

## INTRODUCTION

Over a century ago, pioneering work by Cajal demonstrated the amazing morphological diversity of interneurons. Recent work by numerous groups have been able to categorize this diversity at a molecular level using single cell genomics.^1–4^ Patch-seq and spatial transcriptomics allow the cellular complexity of these molecularly classified populations to be examined. However, these approaches are insufficient to fully understand their function across mammalian brains. To do so requires the ability to target and manipulate specific interneuron types to explore their complexity *in situ*.

Work over the past two decades has provided approaches for cell-type specific targeting in genetically amenable species such as mice, but the cost and time required to do so is often daunting. In addition, the inability to effectively target cell types in non-human primates (NHPs) and other less genetically tractable species have slowed down similar progress in other mammals. Recombinant adeno associated viruses (rAAVs) can drive long-term gene expression *in vivo* and have become a popular tool for gene delivery. The specificity of rAAV-mediated gene expression can be controlled by capsid choice and gene regulatory elements. Over the past eight years, beginning with the use of mDLX,^5^ a pan-interneuron enhancer, we and others have increasingly used cell-type specific enhancers in the context of AAVs as an effective way to target different cell classes, including interneuron subtypes, in a manner that is both economically and temporally expedient across mammalian species.^6–11^ While we previously reported the discovery of a parvalbumin (PV)-specific enhancer to broadly target this population of interneurons,^6^ progress towards finding enhancers that allow targeting of cardinal and recently defined subtypes of interneurons have until now proven elusive. Here we present our efforts to broaden the toolkit for targeting interneuron classes, particularly within the cerebral cortex.

Depending upon the criteria, interneuron subtypes at present are thought to range in number from approximately 30-120 subtypes.^1–4^ This diversity appears to center around four major large cardinal classes, PV, somatostatin (SST), vasoactive intestinal peptide (VIP) and lysosomal-associated membrane protein 5 (LAMP5). Each of these cardinal groups include multiple different subclasses. The ability to target each of these subtypes would provide an ideal toolset to explore interneuron function and connectivity. At present our goal is to identify enhancers that work in the context of AAVs to target each of the major cardinal classes, and the most prominent subclasses within each of these cardinal divisions. To this end we present here a set of seven enhancers. These include the two major PV subclasses, basket and chandelier cells, two major SST classes, non-Martinotti interneurons and broadly across the infragranular classes, as well as the cardinal VIP, LAMP5 and cholinergic groups. Notably, while each of these enhancers can deliver a fluorescent reporter specific to these seven interneuron classes, they also work well in conjunction with a number of different payloads, including optogenetic tools, activity reporters, and cell biological reporters (e.g. synapses). In addition, the identified enhancer sequences are conserved between mice and primates and largely work across species. Here we present each of them in the context of rodents and then NHPs, and in select cases demonstrate their functionality *in vivo*. Despite these obvious advantages, the specificity and sensitivity of these enhancers varies in accordance with their viral titer and method of introduction. To maximize their usefulness to the community, we have tested each of these enhancers across a range of titers and delivery methods and present each with an optimized standard operating procedure (SOP). In the context of enhancers and parallel approaches developed by others participating in this BRAIN Initiative Armamentarium, we believe this present toolkit represents a major step forward in the accessibility of cortical interneurons for discovery and experimentation.

## RESULTS

### Identification and testing of cell-type specific enhancers for cortical interneurons and striatal cholinergic neurons

To identify potential cell type specific enhancers, we took advantage of the single-cell assay for transposase accessible chromatin using sequencing (scATAC-seq)^12^ to capture the chromatin accessibility patterns among different cell types for both cortical interneuron and striatal neuron populations. To enrich for GABAergic neurons, Dlx5a-Cre::INTACT mice were used. GFP^+^ nuclei from cortex (anterior lateral motor cortex, ALM) and striatum were isolated by FACS and scATAC-seq data were collected (12,403 nuclei from cortex and 7,401 nuclei from striatum after quality control) (Figure 1A). Cells were first clustered based on the similarities of chromatin accessibility and then annotated based on the inferred activity of known marker genes for the cortex (Figure 1B) and the striatum (Figure 1C), respectively. Next, we identified top cell type-specific enhancers using an accurate and fast cosine similarity-based method, COSG,^13^ and selected the top enhancer candidates for *in vivo* testing.

**Figure 1.**
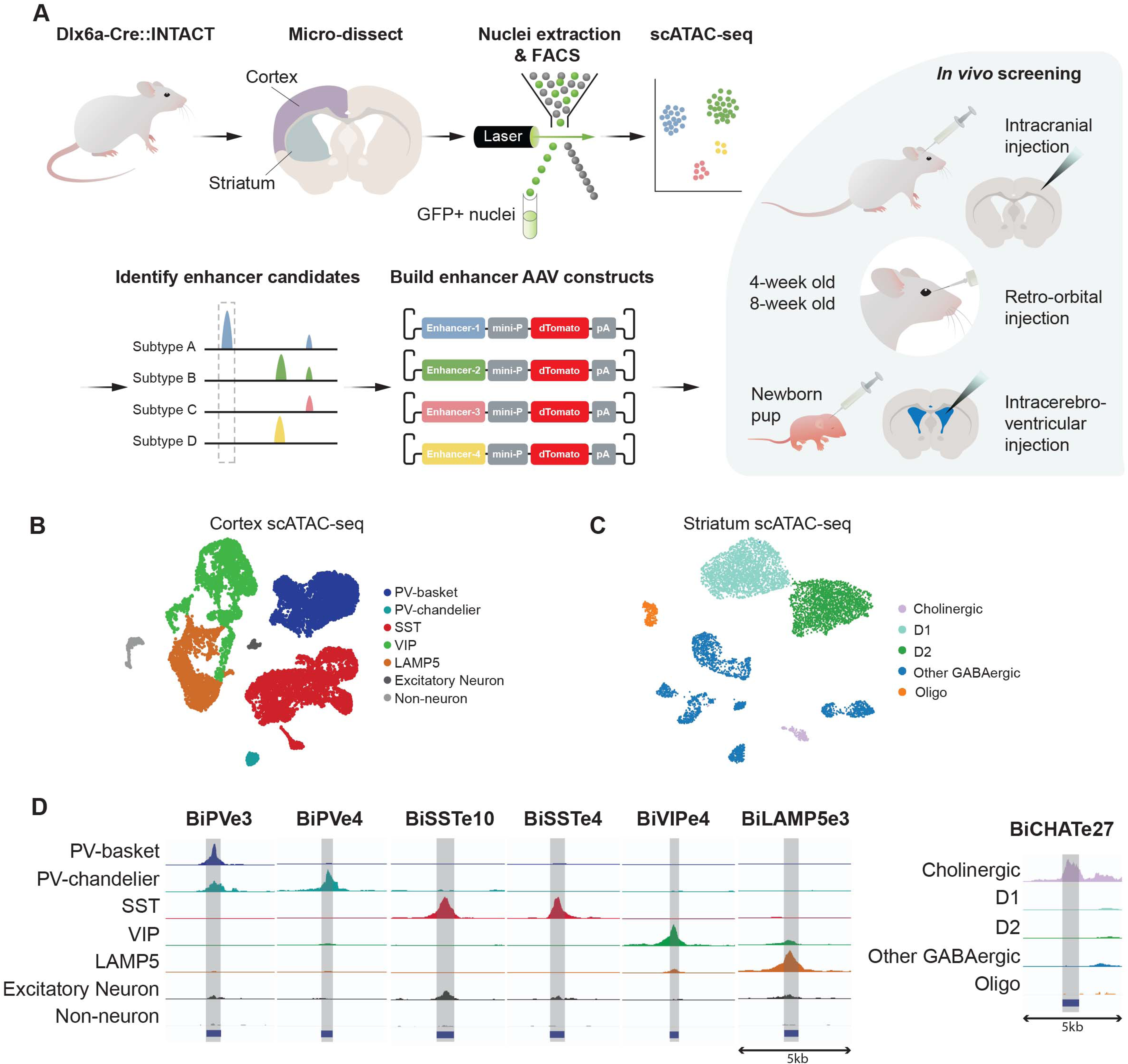
Strategies to identify and characterize enhancer-based AAVs tools with cell type specificity. (A) A schematic illustrating the identification and testing of cell type-specific enhancers in rAAV constructs. (B and C) UMAP projections of single cell ATAC-seq data from mouse cortex (B) and striatum (C). Cells are colored by annotated cell types. (D) Chromatin accessibility pattern of the 7 top enhancers, visualized as normalized genome browser tracks representing the aggregated signals of cells from different cell types..

The sequences for the selected enhancer candidates were cloned into a rAAV construct to drive the expression of the dTomato fluorescent reporter, along with a minimal promoter. These constructs were then packaged into rAAV vectors with the PHP.eB capsid as an effective method to cross the blood-brain barrier (BBB)^14^ and tested for their specificity for the intended target population *in vivo*. The AAVs with enhancer candidates were delivered either by retro-orbital (RO) injection into 4-week or 8-week old mice, intracerebroventricular (ICV) injection into P0/P1 pups, or by intracranial stereotactic injection in 5-8 weeks adult mice or pups (Figure 1A). Three weeks after injection, the brains were harvested and processed. Enhancer performance was evaluated based on two parameters: 1) specificity for the intended target population, which was determined by the percentage of co-localization of dTomato-positive cells labeled by the AAV-enhancer tool and cell-type specific markers of interest, labeled using specific antibodies (when available) or *in situ* probes (# of dTom+marker+ / # of dTom+); 2) sensitivity (# of dTom+marker+ / # of marker+) indicating the efficiency of a given enhancer in targeting all the cells of a given population, in a given area and under the conditions used.

Among all the enhancer candidates tested *in vivo*, 7 of them were highly specific for a specific interneuron or cholinergic neuron subtype and were further characterized. All 7 top enhancers show specific enrichment of ATAC-seq signals in their target cell types (Figure 1D). We named these enhancers “Bi” (Broad Institute) followed by the target population and enhancer number (for example, “BiPVe3” stands for “Broad Institute” PV enhancer e3).

### Enhancer-based viral targeting of PV neuron subtypes in mice

Having previously identified both a pan-interneuron enhancer^5^ and a pan-PV enhancer^6^, we aimed to identify enhancers specific to the two major PV subtypes, the basket cells and chandelier cells. We designed vectors including the top enhancer candidates for basket and chandelier cells. We then generated rAAVs expressing dTomato reporter mediated by these sequences and injected them individually in mice for *in vivo* testing. Two of these nine AAVs tested showed high specificity and efficiency in targeting the cell types of interest (BiPVe3 for basket, and BiPVe4 for chandelier neurons).

To more precisely evaluate the two top enhancers’ activity across the mouse central nervous system, we first analyzed the dTomato reporter expression across the whole brain. To do this, we retro-orbitally injected rAAVs in 4 weeks-old mice and we collected parasagittal sections from 5 standard medio-lateral coordinates and quantified the number of dTomato-positive cells (see methods for details) posteriorly from the cerebellum and anteriorly through the olfactory bulbs, in order to thoroughly profile the major mouse brain areas (Figures 2A and 2B). Importantly, for all the analysis performed in this study, dTomato signal was not amplified using antibodies (except for Figures 5E, 6, and S8C), in order to accurately readout the endogenous levels of reporter expression. This allowed us to better evaluate the enhancer activity in modulating effector genes expression across brain regions and cell types.

**Figure 2.**
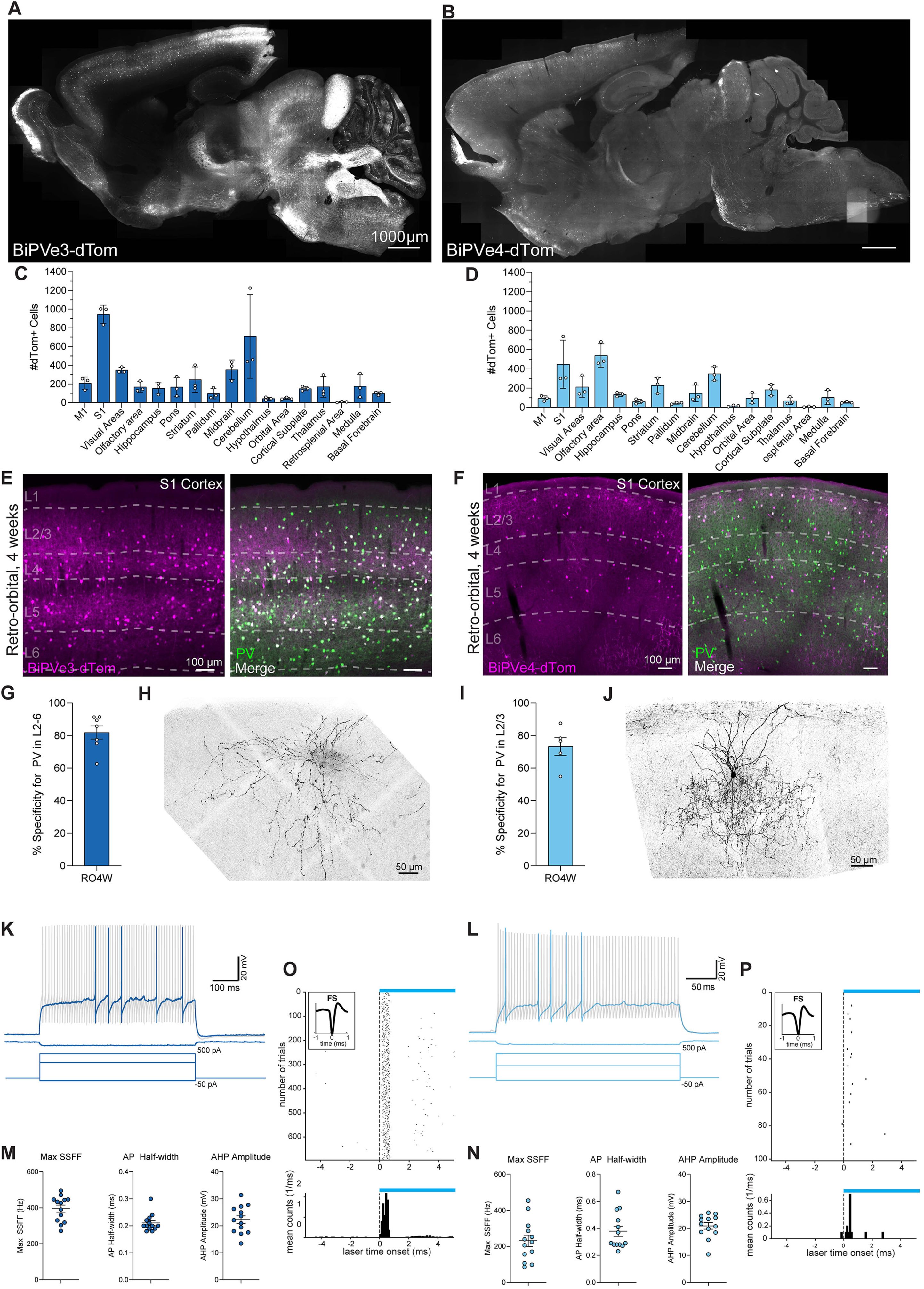
Enhancer-based viral targeting of PV neuron subtypes in mice. (A and B) Representative para-sagittal sections showing the distribution of dTomato-positive cells resulting from expression controlled by the BiPVe3 (A) or BiPVe4 (B) enhancers within the mouse brain, three weeks after retro-orbital AAV injection at 4 weeks of 2.0E+11 total vg/mouse. Scale bar is indicated in the figure. (C and D) Bar graphs showing the number of dTomato-positive cells under the control of the BiPVe3 (C) or BiPVe4 (D) enhancers in the brain areas indicated. Data from 5 sagittal sections from 5 different medio-lateral coordinates were collected, to cover the majority of brain regions; each individual datapoint represents a distinct biological replicate (N=3 mice). Error bars represent standard error of the mean (SEM). (E and F) Representative images showing the expression of dTomato reporter-expressing cells *(in magenta*) under the control of the BiPVe3 (E) or BiPVe4 (F) enhancers, and parvalbumin (*Pvalb*)-positive cells (*in green*) in the primary somatosensory cortex (S1) of the mouse brain, following retro-orbital AAV injection at 4 weeks. dTomato-positive cells co-expressing the PV marker (merge) are labeled *in white*. Cortical layers (L1-6) and scale bars are indicated in the figure. (G and I) Bar graphs showing the percentage specificity of BiPVe3 (G) or BiPVe4 (I) in targeting PV-positive neurons in the indicated layers. N=7 and 6 mice, respectively. Error bars represent SEM. RO4w= retro-orbital injection at 4 weeks of age (H and J) Representative confocal stack of a biocytin-filled cell expressing BiPVe3-dTomato (H) or BiPVe4-GFP (J) in L2/3 of primary somatosensory cortex, displaying characteristic PV basket (H) and chandelier (J) cell morphology. Scale bar is indicated in the figure. (K and L) Representative traces from current clamp recordings of cells expressing (K) BiPVe3-dTomato or (L) BiPVe4-GFP in L2/3 primary somatosensory cortex in response to 600 ms (K) or 300 ms (L) square wave current injections. In both cases, rAAV transduced cells show classical, non-accommodating, fast-spiking firing behavior. (M and N) BiPVe3-dTomato and BiPVe4-dTomato expressing cells in L2/3 display characteristic electrophysiologic properties of fast-spiking PV-positive interneurons, including a high maximum steady state firing frequency (SSFF), a narrow action potential (AP) halfwidth, and large afterhyperpolarization (AHP) (*n* = 13 cells from *N* = 4 mice for BiPVe3; *n* = 13 cells from *N* = 2 mice for BiPVe4: 4 cells from V1, 5 cells from A1 and 4 cells from S1). Error bars represent SEM. (O and P) Example of fast-spiking cell showing successful optogenetic *in vivo* activation across trials of cells transduced with with rAAV-BiPVe3-ChR2 (O) and rAAV-BiPVe4-ChR2 (P) viruses.

The BiPVe3 enhancer shows strong activity in putative basket cells across all the major cortical areas and layers (and as expected showed none in layer 1 (L1) and top layer 2/3, which are known not to contain this population), in hippocampus, in putative Purkinje cells in the cerebellum, as well as in other areas such as the midbrain or the olfactory bulb (Figures 2A and 2C). By contrast, the BiPVe4 enhancer targets cells predominantly located in more superficial cortical areas (in accordance with the characteristic anatomical distribution of chandelier cells), in the pyramidal layer of the hippocampus, as well as in putative fast-spiking neurons of the striatum (which likely share genetic homology with chandelier populations). In addition, we observed moderate labeling in areas such as the midbrain, hypothalamus, cerebellum and medulla (Figures 2B and 2D).

Analysis using fluorescent in situ hybridization (FISH) for a pan-GABAergic neuron marker (*Gad2*) showed that the vast majority of rAAV transduced dTomato-positive cells in the cortical regions belong to the broad class of GABAergic interneurons (average specificity of 84.22% ± 2.67% for BiPVe3 and 78.15% ± 7.33 BiPVe4, Figure S1A and S1C).

To evaluate the cell type-specificity of the two enhancers for PV neuron subtypes, the distribution of cells in the primary somatosensory cortex (S1, as a model cortical region) was quantified, particularly with regard to cortical laminar position (Figures 2E and 2F). AAV-BiPVe3-dTomato labels cells across all cortical layers (excepting L1), with lower density in the upper regions of L2/3 and bottom region of L6 (Figures 2E and S1B). AAV-BiPVe4-dTomato, on the other hand, labels, as expected from the chandelier population, cells mostly restricted in L2/3, with sparse marking across other cortical layers (Figures 2F and S1D). Both basket and chandelier cells are known to be enriched for the marker gene parvalbumin (*Pvalb*), with basket cells showing 100% colocalization rate with this marker, while only about 50% of the chandelier cells in S1 region express Pvalb.^2,3,15^ Indeed, dTomato-positive cells show in both cases high levels of colocalization with the PV marker, as shown by the high percentage specificity in the expected cortical layers (average specificity of 81.9% ± 10.7% in layers 2-6 for BiPVe3, 73.3% ± 12.2% in layer 2/3 for BiPVe4, Figures 2E, 2F, 2G, 2I, S1B and S1D).

To further validate the successful targeting of these two PV subtypes by these two enhancers, we investigated the differential expression of markers known to be enriched in basket and chandelier cells, such as *Syt2* and *Pthlh*^16^, respectively. We thus performed fluorescent in-situ hybridization (FISH) on brain sections from mice injected with either AAV-BiPVe3-dTomato or AAV-BiPVe4-dTomato vectors and quantified the percentage of colocalization of dTomato with these marker mRNAs in L2/3 and L5 of S1 (Figures S1E-H). BiPVe3-dTomato cells show high specificity (81.85% ± 4.86%) and good sensitivity (63.17% ± 11.67) for the marker *Syt2* in L5 (Figures S1E and S1F, *left panel*). However, only a smaller fraction of BiPVe3-dTomato cells colocalize with *Pthlh* in L2/3 (average specificity of 21.38% ± 4.71% and sensitivity of 33.19 ± 9.67, Figure S1E and S1F, *right panel),* suggesting a specific labeling of basket cells by the enhancer BiPVe3. On the other hand, BiPVe4-dTomato cells show good colocalization in L2/3 with the marker gene *Pthlh*, known to be enriched in a subset of chandelier cell*s* (average specificity of 45.03% ± 0.91% and sensitivity of 33.92 ± 5.62, Figure S1G and S1H, *right panel*). A smaller fraction of BiPVe4-dTomato cells showed expression of *Syt2* (average specificity 39.84% ± 4.37% and sensitivity 17.28% ± 7.48%, Figures S1G and S1H, *left panel*), suggesting a possible unspecific labeling of basket cells by the enhancer BiPVe4.

The Syt2 gene enriched in basket cells encodes for a pre-synaptic molecule found at synaptic terminals. We therefore performed immunohistochemistry for Syt2 in AAV-BiPVe3-dTomato-infected mouse S1 and observed a high number of dTomato-positive synaptic terminals colocalizing with Syt2 (Figure S1I). Similarly, AnkG marks the axon initial segments (AIS) of excitatory pyramidal neurons, which are the primary targets of chandelier cell axon terminals.^18^ Indeed, we observed that terminals of BiPVe4-dTomato cells form characteristic cartridges that colocalize with AnkG in L2/3 of the cortex (Figure S1J). Altogether, these data suggest that BiPVe4 enhancer targets PV-positive chandelier cells, while BiPVe3 efficiently marks PV-positive basket cells in the mouse cortex.

Previous studies using BBB-crossing AAV capsids mostly performed systemic, intravenous injections by delivering AAVs in the retro-orbital (RO) sinus of adult mice.^6–8^ We wanted to test if enhancers show similar specificity and selectivity when delivered with other methods and at different ages. We therefore tested the AAV-BiPVe3-dTomato and AAV-BiPVe4-dTomato viruses by RO injections at 4 and 8 weeks of age, with intracerebroventricular (ICV) injections in P0-1 pups and local intraparenchymal injections in pups (P5) and adult mice (5-8 weeks-old). Both BiPVe3 and BiPVe4 enhancers had the strongest activity and specificity when injected by RO at 4 weeks (Figures 2E, 2F, 2G, 2I, and S1C and S1D). Nonetheless, when injected by RO at 8 weeks, both BiPVe3 and BiPVe4 perform almost equally well (Figure S2A_ii_ and S2B_ii_), as shown by the percentage of specificity and sensitivity (Figure S2C_ii_ and S2D_ii_). When injected perinatally by ICV, BiPVe3-dTomato cells show good specificity for PV neurons across layers; however, L2/3 shows overall lower numbers of dTomato cells, indicating low sensitivity for this cortical area (Figure S2A_i_ and S2C_ii_). On the other hand, by ICV BiPVe4 has very poor labeling efficiency in the cortex (Figure S2B_i_ and S2D_i_), but can drive strong reporter expression in the hippocampus (data not shown). These results may reflect that by ICV injection, these viruses fail to efficiently reach superficial layers. When injected intracranially in adults, BiPVe3 shows high activity in PV neurons, although it shows non-specific labeling of pyramidal neurons located in L5 (Figure S2A_iii_ and S2C_iii_). On the other hand, BiPVe4 maintains good specificity and shows the expected laminar distribution when injected using this method, both in the cortex (Figure S2B_iii_ and S2D_iii_) and hippocampus^17^. Interestingly, local injections in pups demonstrate that BiPVe3 labels PV neurons effectively and specifically during early development: when injected intracranially in S1 at P5, BiPVe3 exhibits up to 90% specificity, particularly in lower layers (Figure S2A_iv_ and S2C_iv_). To our knowledge, BiPVe3 is the first tool that allows labeling of PV neurons as early as the first postnatal week. By contrast, BiPVe4 shows non-specific pyramidal cells labeling when injected locally in pups (S2B_iv_ and S2D_iv_).

The morpho-electrophysiologic properties of cells labeled by rAAV-BiPVe3-dTomato or rAAV-BiPVe4-dTomato using RO-injections were further characterized using whole cell patch clamp recordings in acute brain slices from L2/3 of primary somatosensory, visual or auditory cortices (S1, V1 and A1). Cells labeled with BiPVe3-dTomato in S1 displayed characteristic PV basket cell morphology, with multipolar dendritic arbors and highly branching axonal arbors (Figure 2H)^19^. These cells also displayed electrophysiological properties consistent with fast spiking PV basket cells, including the ability to sustain high frequency firing with minimal adaptation, narrow spikes with a large afterhyperpolarization, and a low input resistance (Figures 2K and 2M; Table S1). In contrast, cells labeled with BiPVe4-dTomato displayed characteristic L2/3 PV chandelier cell morphology, with a dendritic arbor biased towards L1 and a highly branching axonal arbor with cartridges of synaptic boutons vertically oriented along the axon initial segments of pyramidal cells (Figure 2J). Cells labeled with BiPVe4-dTomato across S1, V1 and A1 also displayed fast spiking behavior, with large action potential peak, narrow AP halfwidth, large after-hyperpolarization and large dV/dt maximum values for the first action potential, characteristic of fast-spiking interneurons (Figures 2L and 2N; Table S1). These findings are in agreement with previous reports characterizing chandelier cells.^20,21^

We then tested the possibility of virally opto-tagging chandelier cells and basket cells *in vivo*. We thus expressed the light-gated ion channel channelrhodopsin tagged to an mCherry fluorescent reporter (ChR2-mCherry) under the control of BiPVe3 and BiPVe4. We then intracranially inject these two rAAVs in L2/3 of the primary visual cortex (V1) in adult mice (where both enhancers show good PV specificity and sensitivity, with low off-target labeling) and performed *in vivo* recording in supragranular layers of head-fixed mice. We show that mCherry-positive cells have waveforms characteristic of fast spiking cells (FS), as expected for both basket and chandelier neurons (example cells in Figure 2O and 2P), indicating that these two PV subtypes can be opto-tagged *in vivo* using both BiPVe3 and BiPVe4.

Altogether, these data show that BiPVe3-dTomato and BiPVe4-dTomato cells show anatomical, morphological, molecular and electrophysiological characteristics typical of basket and chandelier neurons. We thus conclude that BiPVe3 and BiPVe4 enhancers can be used as tools to efficiently target these two PV interneuron subtypes in the mouse brain.

### Enhancer-based viral targeting of SST-positive interneuron subtypes in mice

Somatostatin (SST)-positive neurons represent an abundant, heterogeneous class of interneurons in the mouse brain. Spatial transcriptomic analysis, coupled with morphological, anatomical and electrophysiological characterizations, have revealed approximately 9 subtypes across the mouse cortex.^3,4^ Despite their unique properties and flavors, SST interneurons can be broadly divided into two main classes: non-Martinotti and Martinotti SST cells, the latter of which extends axons which ramify in layer 1, while the former projects axons locally.^3,4^ To gain access to the SST populations, we identified a group of candidate enhancers that were predicted to be active in SST-positive interneurons, and we cloned them into rAAV constructs to drive the expression of the dTomato fluorescent reporter, for *in vivo* testing. Of these candidates, BiSSTe10 and BiSSTe4 showed the highest specificity and sensitivity in targeting the SST populations and they each target distinct SST subtypes.

We first assessed the BiSSTe10 and BiSSTe4 activity in the mouse brain, by quantifying in para-sagittal sections the total number of dTomato-positive cells across the whole brain (Figures 3A and 3B), following RO injections in 4 weeks-old mice and ICV injections in pups, respectively. BiSSTe10-dTomato-positive cells heavily label all the major cortical areas, with neurites invading cortical layer 4, as expected for non-Martinotti SST neurons (Figure 3A). In addition, BiSSTe10-dTomato cells sparsely reside in the ventral region of the striatum, olfactory bulb, hypothalamus and the pallidum (Figures 3A and 3C). The soma of BiSSTe4-dTomato cells, on the contrary, mostly reside in deeper (L5) cortical layers (with a bias for more posterior cortical areas such as the somatosensory and visual cortices, possibly due to the injection method) with axons heavily innervating L1. Very sparse dTomato labeling was also observed in the midbrain, olfactory bulb and striatum (Figures 3B and 3D). Similarly to our PV-subtype enhancers, BiSSTe10 and BiSSTe4 show high levels of activity in GABAergic neurons, as shown by the high percentage of colocalization of dTomato-positive cells with the GABAergic marker *Gad2* (average specificity of 78.26% ± 11.03% for BiSSTe10 and 69.24% ± 2.08% for BiSSTe4, Figure S3A and S3C).

**Figure 3.**
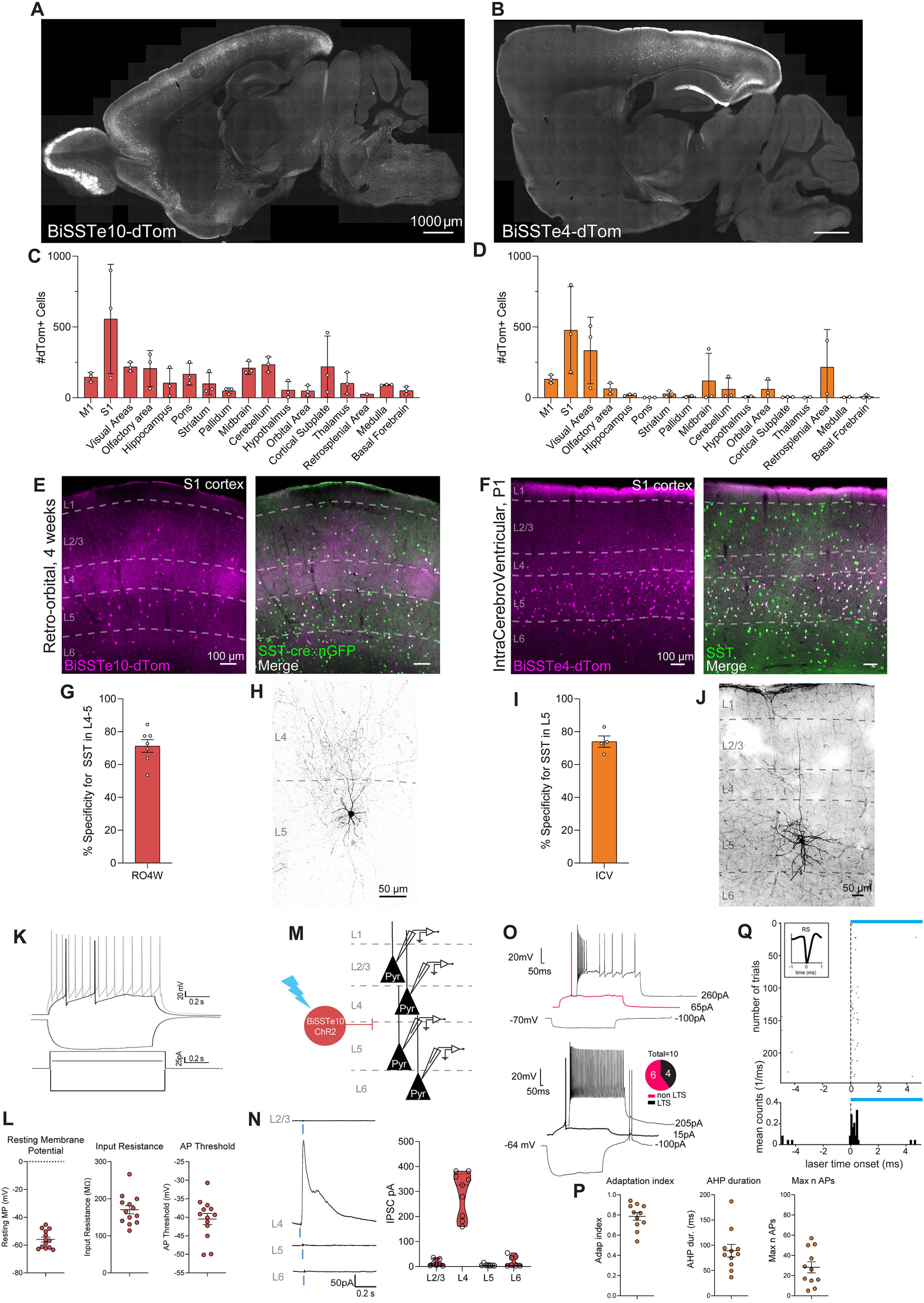
Enhancer-based viral targeting of SST neuron subtypes in mice. (A and B) Representative para-sagittal sections showing the distribution of dTomato-positive cells resulting from expression controlled by the BiSSTe10 (A) or BiSSTe4 (B) enhancers within the mouse brain, three weeks after retro-orbital AAV injection at 4 weeks of 2.0E+11 total vp/mouse (A) or ICV injections of 1.0E+10 total vp/pup at P1, respectively. Scale bar is indicated in the figure. (C and D) Bar graphs showing the number of dTomato-positive cells under the control of the BiSSTe10 (C) or BiSSTe4 (D) enhancers in the brain areas indicated. Data from 5 sagittal sections from 5 different medio-lateral coordinates were collected, to cover the majority of brain regions. Each individual datapoint represents a distinct biological replicate; N=3 mice (D). Error bars represent SEM. (E and F) Representative images showing the expression of dTomato reporter-expressing cells *(in magenta*) under the control of the BiSSTe10 (E) or BiSSTe4 (F) enhancers, and somatostatin (*SST*)-positive cells (*in green*) in the primary somatosensory cortex (S1) of the mouse brain, following retro-orbital AAV injection at 4 weeks or ICV injections at P1, respectively. dTomato-positive cells co-expressing the SST marker (merge) are labeled *in white*. Cortical layers (L1-6) and scale bars are indicated in the figure. (G and I) Bar graphs showing the percentage specificity of BiSSTe10 (G) or BiSSTe4 (I) in targeting SST-positive neurons across the layers indicated. N=7 and 4, respectively. Error bars represent SEM. RO4w= retro-orbital injection at 4 weeks of age, ICV=intracerebroventricular. (H and J) Representative confocal stack of a biocytin-filled cell expressing BiSSTe10-dTomato (H) following RO 4 weeks AAV injection or BiSSTe4-dTomato following ICV injection (J) in S1, displaying characteristic morphology. Scale bar is indicated in the figure. (K) Representative traces of voltage responses to 500 ms step current injection in current-clamp whole-cell configuration of BiSSTe10-dTomato positive cells in L4/5 primary somatosensory cortex, following RO 4 weeks AAV injections. (L) Dot plots showing three intrinsic physiological properties of BiSSTe10-dTomato positive cells. *n* = 13 cells from *N* = 2 mice. Error bars represent SEM. (M) Schematic representation of testing the synaptic connection from BiSSTe10-positive to pyramidal (Pyr) neurons located in different cortical layers. AAV-BiSSTe10-ChR-mCherry was injected by RO at 4 weeks and activity was recorded in Pyr neurons in S1. (N) *left panel*: Example traces of photo-stimulation-evoked postsynaptic currents recorded from pyramidal neurons in the indicated layers at 0 mV under voltage-clamp configuration. Experiments were performed in the presence of TTX and 4AP. Blue bar indicates ChR2 photo-stimulation (470 nm, 1 ms). *right panel:*Violin plot showing IPSCs responses of pyramidal neurons from the indicated layers, following light stimulation of BiSSTe10-dTomato cells. Data highlight a very selective inhibition from BiSSTe10-mCherry-positive cells to L4 pyramidal neurons. *n* = 30 cells (7 for L2/3, 9 for L4, 7 for L5 and 7 for L6), from *N* = 4 mice.(O) *Top*: Representative traces of a fast adapting (non LTS) (*top pane*l) and low threshold spiking (LTS) (*bottom pane*l) recorded from BiSSTe4-tdtomato positive neurons in L5 of S1. LTS=low threshold spikers. (P) Dot plots representing three recorded intrinsic properties of BiSSTe4 cells. Graphs are presented as mean and SEM. n=10 from N=3 mice. (Q) Example of regular-spiking cell showing successful optogenetic *in vivo* activation across trials of cells transduced with rAAV-BiSSTe4-ChR2.

To test the cell type-specificity of BiSSTe10 and BiSSTe4 in targeting SST neurons in the cortex, we investigated the colocalization of dTomato cells with the SST marker in S1. In this cortical region, BiSSTe10 mostly labels cells located in L4/5, and has sparse labeling in L2/3 and L6 (Figure 3E). These cells show a high degree of colocalization with the SST marker (average specificity of 71.3% ± 10.2%) in L4-5, but lower specificity across other cortical layers, as expected (Figures 3E, 3G, S3B, S3D). This analysis further revealed a heavy innervation of BiSSTe10-dTomato cells to L4 (Figures 3E), as expected for non-Martinotti SST neurons, in particular the SST-Hpse subtype, as previously described.^4^ Similarly, BiSSTe4 has the highest activity in deeper cortical layers, such as L5, where it shows good specificity for SST (average specificity of 73.9% ± 6.9%, Figures 3F, 3I, and S3D). Moreover, as distinct from BiSSTe10 and in line with morphological characters of Martinotti SST neurons, cells labeled by BiSSTe4-dTomato mostly project their axons to L1 (Figure 3F).

To determine the identity of the SST subtypes labeled by BiSSTe4 and BiSSTe10, respectively, we tested by FISH the differential enrichment of marker genes known to be expressed in distinct SST populations, such as *Hpse*, *Nmbr, Mme* and *Calb2*.^4^ We quantified the percentage of colocalization of either BiSSTe10- or BiSSTe4-dTomato with these markers in L5 or L6 of S1 (Figures S3E-H). BiSSTe10-dTomato cells show overall good specificity and high sensitivity for *Hpse* in L5 (average specificity: 50.2% ± 15.74% and sensitivity: 74.91% ± 15.33%, Figures S3E and S3F, *left panel*).) and moderate colocalization with the marker *Nmbr* in L6 (average specificity: 32.89% ± 3.48% and sensitivity: 30.34% ± 8.12%, Figures S3E and S3F, *right panel*). On the other hand, BiSSTe4 enhancer shows low activity in either *Mme*- and *Calb2*-positive neurons (average specificity for *Mme* in L5: 10.25% ± 2.02%, average specificity for *Calb2* in L5: 9.8% ± 0.72, Figures S3G and S3H), suggesting that the BiSSTe4 enhancer only partially targets the *Mme*- and *Calb2*-positive SST populations. Importantly, Martinotti and non-Martinotti neurons represent a highly heterogeneous population of SST cells that are distinct in their gene expression, morphology and electrophysiological characteristics. No single marker gene allows for these two broad SST classes to be distinguished. As such while the two enhancers we identify allows these populations to be roughly identified, users will need to carefully employ specific markers and morphologies to ascertain the specific SST populations targeted.

As with our PV subtype-specific enhancers, BiSSTe10 and BiSSTe4 also show highly variable labeling when injected at different ages and via different injection routes. BiSSTe10 shows optimal specificity for SST in cortical L4/5 when injected by RO at 4 weeks (Figures 3E, 3G and S3B), as well as 8 weeks (Figures S4A_ii_ and S4C_ii_). However, BiSSTe10 shows low activity in the cortex upon ICV injections, as indicated by the low sensitivity. Despite the low labeling efficiency, this injection route still results in good high specificity for SST-positive neurons in L4 and L5 (Figures S4A_i_ and S4C_i_). While, BiSSTe4 retains good specificity for SST by RO injection at either 4 or 8 weeks (Figures S4B_i_, S4B_ii_, S4D_i_ and S4D_ii_), it shows optimal activity when injected by ICV (Figures 3F, 3I and S3D). Finally, intracranial injections of both the BiSSTe10- and BiSSTe4-dTomato in S1 of adult mice show moderate specificity for SST neurons in supragranular layers, accompanied by non-specific labeling of putative PV-positive neurons in L2/3 and pyramidal neurons in L5 (Figures S4A_iii_ and S4C_iii_). Interestingly, injections of BiSSTe10-dTomato in V1 yielded higher specificity for SST neurons than in S1 (data not shown).

We next examined the morphology of BiSSTe10-dTomato and BiSSTe4-dTomato cells, as well as the electrophysiological properties of the BiSSTe10-dTomato population following RO injections at 4 weeks or ICV injection on pups, respectively. Whole cell patch clamp recordings were performed upon acute brain slices, in L5 of S1 and V1. Biocytin filling of BiSSTe10-dTomato cells revealed the stereotypical SST-Hpse morphology, with processes heavily innervating L4 (Figures 3H). BiSSTe4-dTomato cells, on the contrary, show extensive arborization in L5/6, with axonal projections to superficial cortical layers (Figure 3J).

As expected, BiSSTe10 positive neurons showed characteristic regular spiking features, with no tendency for hyperpolarization-induced rebound bursting (Figure 3K and 3L).^18^

BiSSTe4-tagged cells are primarily localized in the deep layers following ICV injections. Thus, we examined the electrophysiological properties of marked cells, focusing on L5. The recorded cells exhibited mixed characteristics typical of L5 Martinotti including a moderate or fast-adapting firing pattern (Figure 3O).^22,23^ A subset of these cells displayed rebound bursting after hyperpolarization (low threshold spikers; LTS), previously associated with T-shaped and Fanning out L1-projecting subtypes in S1 (n=4/10). In agreement with this result, morphological reconstructions of biocytin-filled cells confirmed that a proportion of the recorded population sent axons to the supragranular layers (Figure 3J). A substantial fraction of non-LTS cells exhibited a rapidly adapting firing pattern, characterized in some cases by an extremely low maximum spike count in response to increasing current injection (Figure 3O). Between these cells, a variable afterhyperpolarization (AHP) duration was observed following the action potential, indicating the potential presence of a non-specific neuronal population among the tagged neurons. (Figure 3P and Table S1). A prolonged AHP duration has been proposed as an electrophysiological signature of the Martinotti cell subpopulation in the deep layers of the cortex.^22^

Non-Martinotti, SST-Hpse neurons have been shown to selectively inhibit L4 pyramidal neurons in the cortex.^4^ To confirm that our BiSSTe10 enhancer indeed labels SST-Hpse neurons, we injected by RO at 4 weeks a rAAV construct expressing Channelrhodopsin, under the control of the BiSSTe10 enhancer (AAV-BiSSTe10-ChR2-mCherry). Whole cell recording of pyramidal neurons located in different cortical layers (L2/3, L4, L5 and L6), following light stimulation of BiSSTe10 cells, as expected revealed a highly selective IPSC response in L4 pyramidal neurons (Figure 3M, 3N and 3O), but not in other layers. Together, these results strongly demonstrate the specificity of BiSSTe10 in targeting non-Martinotti SST-Hpse neurons in the mouse cortex.

To further investigate the activity of BiSSTe4 targeted cells, an rAAV construct carrying channelrhodopsin under the control of this enhancer was created (AAV-BiSSTe4-ChR2-mCherry). Following intracranial injection of this vector into V1, we performed *in vivo* recordings in supragranular layers of head-fixed mice, where BiSSTe4 enhancer shows good specificity for SST neurons. As expected for Martinotti SST neurons, BiSSTe4-labeled cells show wave properties typical of regular spiking (RS) cells (Figure 3Q).

Altogether, these data show that BiSSTe10 and BiSSTe4 enhancers efficiently label both non-Martinotti and Martinotti SST-neurons, with each targeting a cross-section of these types within the cortex.

### Enhancer-based viral targeting of CGE-derived interneuron subtypes in mice

Unlike the SST- and PV-positive neurons, which entirely originate from the medial ganglionic eminence (MGE), the other two main GABAergic cardinal classes (VIP- and Lamp5-positive) arise from the caudal ganglionic eminence (CGE).^24–26^ Despite their shared developmental location, these additional interneuron types are morphologically and functionally quite distinct from one another: VIP neurons are known to inhibit other GABAergic neurons, while LAMP5 neurons provide long-lasting inhibition to pyramidal neurons.^27–29^ Given their important roles in cortical function, we aimed to develop tools to target and manipulate these two cardinal interneuron classes. We thus selected the top enhancer candidates that showed high predicted activity in these two populations, and generated rAAV expressing dTomato reporter under their control for *in vivo* testing in mice using a combination of viral delivery routes and ages. Of these top candidates, BiVIPe4 and BiLAMP5e3 showed the highest efficiency in selectively targeting VIP and LAMP5, respectively. Interestingly, the BiVIPe4 enhancer is derived from a 1kb-long sequence that was originally chosen. We then performed enhancer bashing and identified its core sequence, BiVIPe4. This enhancer is 399bp in length and is more efficient than the original sequence in labeling VIP cells.

To address the overall activity of these two enhancers, we readout their ability to drive the expression of the dTomato reporter across the mouse brain. AAV-BiVIPe4-dTomato was injected by ICV at perinatal ages, as rAAVs with VIP-specific enhancers packaged in PHP.eB capsid do not work well by RO injection (Jonathan Ting, Allen Institute, personal communication). Three weeks post injection, we quantified the total number of dTomato-positive cells in selected brain areas and observed that BiVIPe4 shows overall moderate degree of GABAergic neurons labeling with 36.69% ± 1.16% specificity for *Gad2*. (Figure S5A), however it is active in a sparse population of putative VIP neurons in the cortex, with higher accumulation in superficial cortical layers as expected for this cell class. Sparse cell labeling can be observed also in areas such as the hippocampus, midbrain and the striatum (Figures 4A and 4C).

**Figure 4.**
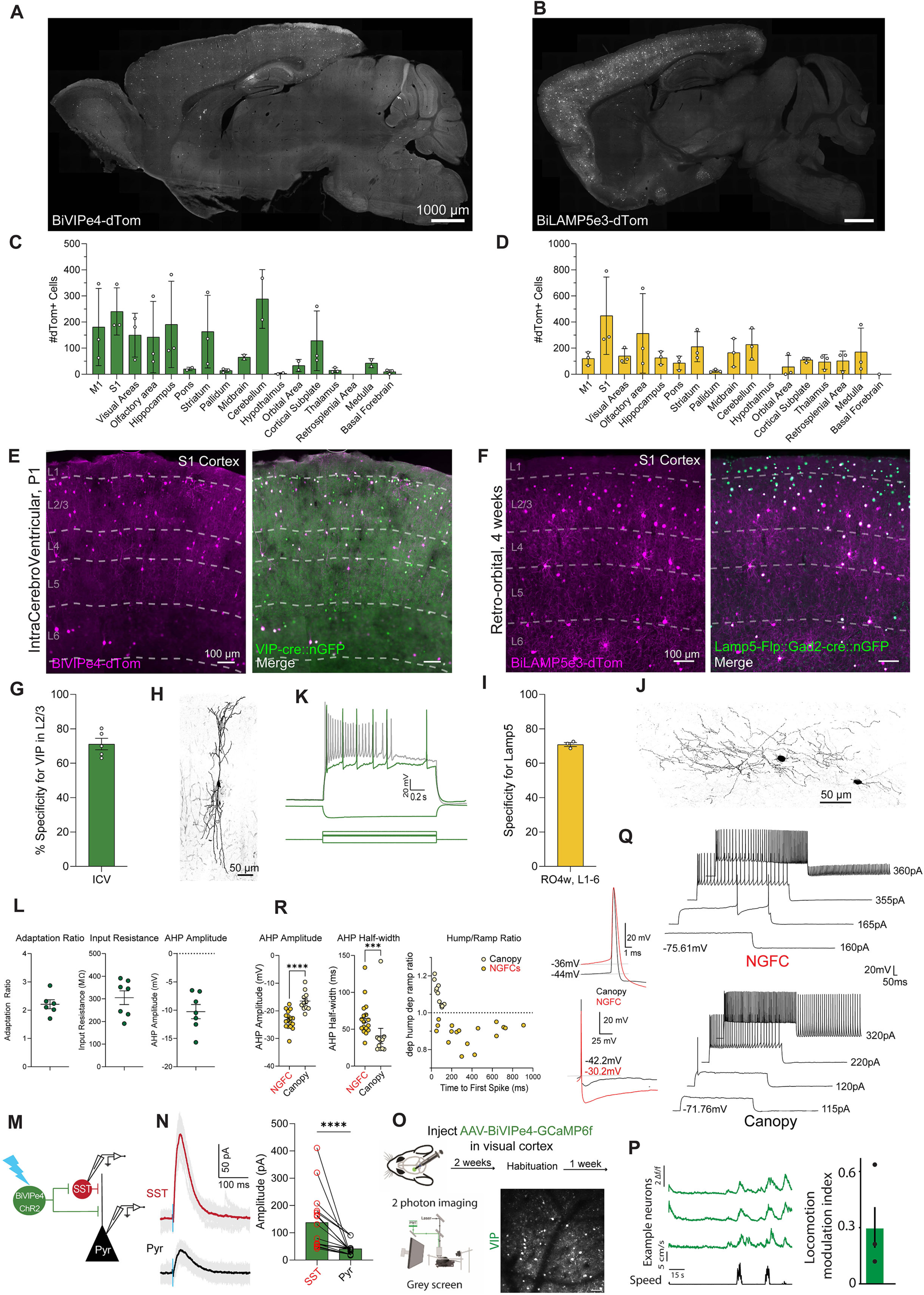
Enhancer-based viral targeting of CGE-derived cINs in mice. (A and B) Representative para-sagittal sections showing the distribution of dTomato-positive cells resulting from expression controlled by the BiVIPe4 (A) or BiLAMP5e3 (B) enhancers within the mouse brain, three weeks after ICV injections of 1.0E+10 total vg/pup at P1 (A) or retro-orbital AAV injection at 4 weeks of 2.0E+11 total vg/mouse (B) respectively. Scale bar is indicated in the figure. (C and D) Bar graphs showing the number of dTomato-positive cells under the control of the BiVIPe4 (C) or BiLAMP5e3 (D) enhancers in the brain areas indicated. Data from 5 sagittal sections from 5 different medio-lateral coordinates were collected, to cover the majority of brain regions. Each individual datapoint represents a distinct biological replicate; N32 mice. Error bars represent SEM. (E and F) Representative images showing the expression of dTomato-expressing cells *(in magenta*) under the control of the BiVIPe4 (E) or BiLAMP5e3 (F) enhancers, and vasoactive intestinal peptide (*Vip*)-positive cells (E) or the genetically-expressed GFP reporter under the control of Lamp5-Flp and Gad2-cre (F) (*in green*) in the primary somatosensory cortex (S1) of the mouse brain, following ICV injections at P1 (E) or retro-orbital AAV injection at 4 weeks (F), respectively. dTomato-positive cells co-expressing the marker (merge) are labeled *in white*. Cortical layers (L1-6) and scale bars are indicated in the figure. (G and I) Bar graphs showing the percentage specificity of BiVIPe4 (G) or BiLAMP5e3 (I) in targeting VIP- or LAMP5-positive neurons in L2/3 (G) or across all cortical layers (I). N=5 and N=3 mice, respectively. Error bars represent SEM. (H and J) Representative morphological reconstructions of neurobiotin-filled BiVIPe4-mCherry-positive neuron in L2/3 (H) and a BiLAMP5e3-dTomato (J) neuron in L1 of S1. Scale bar is indicated. (K) Representative traces of voltage responses to 800 ms step current injection in current-clamp whole-cell configuration of BiVIPe4-mCherry-positive cells. Scale bar is indicated. (L) Summary plots showing the intrinsic physiological properties of BiVIPe4-mCherry--positive cells. *n* = 7 cells from *N* = 2 mice for BiVIPe4. Error bars represent SEM. (M) Schematic representation of testing the synaptic connection from BiVIPe4-positive cells to SST-positive and pyramidal (Pyr) neurons. AAV-BiVIPe4-ChR-mCherry was injected at P0 into GIN-GFP mice and photo-evoked activity was recorded in SST and Pyr neurons in S1. (N, *left panel*) Example traces of photo-stimulation-evoked postsynaptic currents recorded from an SST cell and a pyramidal neuron at 0 mV under voltage-clamp configuration. Gray traces depict individual sweeps, and solid traces the average of these sweeps. Blue bar indicates ChR2 photo-stimulation (470 nm, 3 ms). (*right panel*): Histogram showing the amplitude of postsynaptic response of SST and simultaneously recorded nearby pyramidal neurons upon optogenetic stimulation of BiVIPe4-labeled cells. *n*=16 SST and pyramidal cells from N=4 mice. Error bars represent SEM. (O) Imaging locomotion responses of BiVIPe4 targeted neurons in the visual cortex. Mice were injected with AAV-BiVIPe4-GCaMP6f in the visual cortex. After operation recovery, mice were habituated to head-fixation and were able to run freely on a belt treadmill. Following habituation, spontaneous activity in VIP interneurons was acquired while the mice were presented with a gray screen. *Bottom left*: Set up for two-photon imaging of VIP interneurons in the visual cortex of awake-behaving mice. *Bottom right*: An example image plane in L2/3 showing GCaMP6f-expressing neurons. (P) Traces of three example neurons of three neurons (*orange*) in relation to running speed (*black*). *Right panel*: Mean of the locomotion modulation index (LMI; LMI = (RL–RM) / (RL+RM)). Individual points represent the mean LMI for each mouse (N= 3 mice). (Q) *Left panel:* Representative traces of action potentials recorded from L1 NGFC and Canopy interneurons. The top panel shows action potentials aligned to the maximum amplitude, while the bottom panel aligned to their action potential threshold. *Right panel*: Two examples of an NGFC and a Canopy cell responding to a 1-second increasing current injection. Differences in electrophysiological properties can be observed at subthreshold, threshold, and suprathreshold current steps. (R) Summary plots showing three intrinsic physiological properties of BiLAMP5e3-dTomato-positive cells. *n*=17 cells for NGFCs and *n*=11 cells for Canopy, from N=5 mice. Error bars represent SEM.***p<0.001 ****=p<0.0001 Mann Whitney test.

AAV-BiLAMP5e3-dTomato that is designed to target the LAMP5 expressing population was instead injected by RO at 4 weeks. BiLAMP5e3 shows high activity in the GABAergic population with 91.10% ± 2.02% average specificity for *Gad2* (Figure S5C) and presumptive LAMP5 neurons spread across the major cortical areas, but as expected, are more concentrated in superficial layers. Sparse labeling can be observed also in other areas like the hippocampus, striatum, medulla and the thalamus (Figures 4B and 4D).

We further validated the identity of BiVIPe4-dTomato and BiLAMP5e3-dTomato neurons by assessing their position, morphology and marker gene expression. We calculated the specificity of the BiVIPe4 enhancer by quantifying the number of dTomato cells colocalizing with the VIP marker in S1 (Figure 4E and 4G). BiVIPe4-dTomato cell soma appear small in size, a typical character of VIP neurons, and mostly located in superficial cortical layers (Figure 4E). As predicted, BiVIPe4-dTomato cells show a high degree of colocalization with the VIP marker across cortical layers, with highest levels in L2/3 (average specificity of 71.1% ± 7.4% in L2/3, Figures 4G and S5B). Contrary to BiVIPe4-dTomato, BiLAMP5e3-dTomato cells appear highly ramified and present large somas, typical of LAMP5 interneurons. Cell type-specificity of BiLAMP5e3 was assessed by quantifying the colocalization of dTomato with the green fluorescent (GFP) reporter whose selective expression was driven by the intersectional cross between LAMP5-Flp and Gad2-cre mice (Figure 4F). This intersectional strategy has been demonstrated to selectively label LAMP5-positive interneurons in the mouse cortex.^29^ BiLAMP5e3-dTomato cells show good specificity for LAMP5 interneurons in S1, across all cortical layers (average specificity of 70.7% ± 2.0% in L1-6, Figures 4F, 4I, and S5D).

As previously mentioned, rAAVs with VIP-specific enhancers packaged in PHP.eB capsid tend to work best when injected by ICV perinatally. When injected by RO either at 4 weeks or 8 weeks postnatally, BiVIPe4 shows very low labeling efficiencies (as evident by the percent sensitivity) and low specificity (Figures S6A_i_, S6A_iI_,S6C_i_ and S6C_iI_). While intracranial injections of AAV-BiVIPe4-dTomato in pups (Figure S5E and S5F) and in the adult provide better specificity for VIP neurons, this route of injection also generates a small degree of off-target labeling, particularly in putative LAMP5 neurons in L1 and PV neurons in L2/3 (Figures S6A_iii_ and S6C_iiI_). AAV-BiLAMP5e3-dTomato, on the other hand, shows optimal activity and specificity when injected by RO at 4 weeks (Figure 4F, 4I and S5D) and at 8 weeks (Figures S6B_ii_ and S6D_ii_). Conversely, when introduced by ICV, BiLAMP5e3 shows lower activity, as evident by the lower percent sensitivity (Figures S6B_i_ and S6D_i_). Finally, similarly to BiVIPe4, intracranial injections of AAV-BiLAMP5e3-dTomato result in good specificity for LAMP5 in superficial cortical layers, albeit with some off target labeling in PV cells (Figures S6B_iii_ and S6D_iii_).

To further validate the BiVIPe4 and BiLAMP5e3 enhancers, the morphological and electrophysiological properties of rAAV transduced cells were examined. To this end, following intracranial injection of the AAV-BiVIPe4-dTomato in P0-3 pups (Figure S5E and S5F), we performed *ex vivo* whole-cell recordings from dTomato-positive neurons in the superficial layers of S1. Four weeks post injection, most BiVIPe4-dTomato neurons exhibit a bipolar morphology, typical of cortical VIP neurons (Figure 4H).^30,31^ In addition, their firing pattern, intrinsic properties and high input resistance were consistent with them having a VIP identity (Figures 4K,L).^30,31^ Given the known strong functional connectivity from VIP neurons to SST cells,^27,32,33^ we further tested whether BiVIPe4-dTomato cells induce strong inhibitory postsynaptic current (IPSC) in SST cells.^27^ We intracranially injected AAV-BiVIPe4-ChR-mCherry at P0 to the GIN-GFP mouse line, in which a subset of SST cells express GFP (Figure 4M). To control for injection and expression variability of ChR expression, we recorded from GFP-positive cells together with nearby pyramidal neurons and compared the light-evoked response in GFP-positive cells to the response in the pyramidal neurons. Consistent with previous findings, BiVIPe4-ChR-mCherry cells elicited significantly larger inhibition in SST cells than in pyramidal cells (Figure 4N). Together, these results strongly support the specificity of the BiVIPe4 enhancer in targeting VIP inhibitory neurons.

To further investigate the activity of BiVIPe4-targeted cells, an AAV construct carrying the calcium sensor GCaMP6f under the control of this enhancer was created (AAV-BiVIPe4-GCaMP6f). Following injection of this virus into the visual cortex, animals were allowed to recover from the surgery. Post-recovery, mice were habituated to head-fixation and were able to run on a belt treadmill (Figure 4O). We first confirmed the GCaMP6f expression specificity for VIP interneurons by FISH and observed good specificity and sensitivity of BiVIPe4-GCaMP6f for VIP neurons in L2/3 (Figure S7C and S7D). Three weeks after surgery, spontaneous activity of BiVIPe4-targeted cells in L2/3 were recorded (Figure 4O). In line with previous findings,^34,35^ locomotion increased the mean amplitude of the response of putative VIP neurons (Figure 4P). To quantify the effect of locomotion, we computed the locomotion modulation index (LMI; 0.29 +/- 0.05; see Methods) for all recorded neurons. An LMI greater than zero indicates a positive correlation of the Δf/f0 with locomotion.

To assess the visual responses of the BiVIPe4-labeled cells, we recorded neuronal activities while the mice were presented with drifting sinusoidal gratings in eight directions and two different contrasts (100% (high) and 25% (low) (Figure S7A and S7B). Consistent with previous findings,^36^ we found similar proportions of neurons responding to both low and high contrast (low contrast: Activated: 50/130 (38.5%), suppressed: 21/130 (16.2%), not modulated: 59/130 (45.4%), high contrast: Activated: 49/128 (38.5%), suppressed: 22/128 (17.2%), not modulated: 57/128 (44.5%)). We computed the global orientation selectivity index (gOSI) of the BiVIPe4-labeled neurons activated by the visual stimuli, and found that on average, the gOSI was similar between the two contrast conditions used (High contrast gOSI = 0.09 + 0.01; low contrast gOSI = 0.10 + 0.01) (Figure S7A and S7B).

On the other hand, BiLAMP5e3-dTomato cells show a ramified morphology, with axons broadly innervating the cortical layers they reside in (Figure 4J).^29^ Whole cell recordings of BiLAMP5-dTomato cells following intracranial injections in S1 revealed that this enhancer shows activity in electrophysiologically distinct neuron types in L1, overlapping two well defined L1 subpopulations: early spikers Canopy (Canopy) and late spikers Neurogliaform cells (NGFCs) (Figure 4Q). We distinguished these two subtypes by using the most distinctive electrophysiological properties and their firing profile in current clamp configuration. NGFCs presented a near-threshold depolarizing ramping potential which converged in a delayed first spike at threshold (Figure 4R). Associated to this parameter, we found a pronounced and prolonged afterhyperpolarization (AHP) after the action potential, typical of this neuronal subtype.^37^ These parameters differed in Canopy interneurons characterized by a discrete depolarizing hump pre-threshold, and an early onset spike at threshold, followed by a faster and less pronounced AHP (Figure 4Q and 4R). These two subtypes exhibited a significant difference also in action potential width, threshold, and amplitude (Table S1). Electrophysiological features that have been shown in previous studies to effectively distinguish between these subtypes.^37,38^ In L2/3, intracranial injections unspecifically label some non-LAMP5-positive cells (Figure S6B _iii_). In line with this data, whole-cell slice recording reveals the presence of cells that show firing pattern typical of NGFC (late spikers LS), as well as fast (FS)- and regular-spiking (RS) neurons (putative PV and SST interneurons, respectively; n=12; 2 FS,5 RS;5 LS) (Figure S5G and S5H). Altogether, our multivariate analysis strongly demonstrates that BiVIPe4 and BiLAMP5e3 enhancers can be used as efficient tools to target VIP and LAMP5 interneurons in the mouse cortex.

### Enhancer-based viral targeting of Chat neurons in mice

Cholinergic (choline acetyl transferase, Chat) neurons are a separate class of neurons that use acetylcholine (ACh) as their primary neurotransmitter. In the mouse brain, they represent an important class of cell with a crucial role for a variety of functions including motor control, memory, and attention.^39^ Being able to target and manipulate them would have several advantages both in research and in clinics. To gain access to this population, we isolated GFP+ nuclei from the striatum of Dlx5a-Cre::INTACT mice and performed single-cell ATAC-seq experiments. Using this data, we identified several candidate enhancers for striatal cholinergic neurons. A few of these enhancers with the highest confidence score were selected and cloned for further *in vivo* testing in mice. One of these candidates (BiCHATe27) showed the best labeling efficiency and expression selectivity in Chat neurons and was further characterized.

To evaluate the activity of this enhancer in the CNS of mice, we injected AAV-BiCHATe27-dTomato vector retro-orbitally in 4 weeks-old mice and assessed the cell labeling across brain sections. The dTomato-positive neurons populate the striatal regions (most abundantly the caudoputamen, as expected for cholinergic neurons), with some labeling also in cortical areas (Figures 5A and 5B) in numbers consistent with the expected ACh-expressing population.^42^ In addition, BiCHATe27 shows mild activity in other nuclei of the midbrain, thalamus and medulla (Figures 5A and 5B).

**Figure 5.**
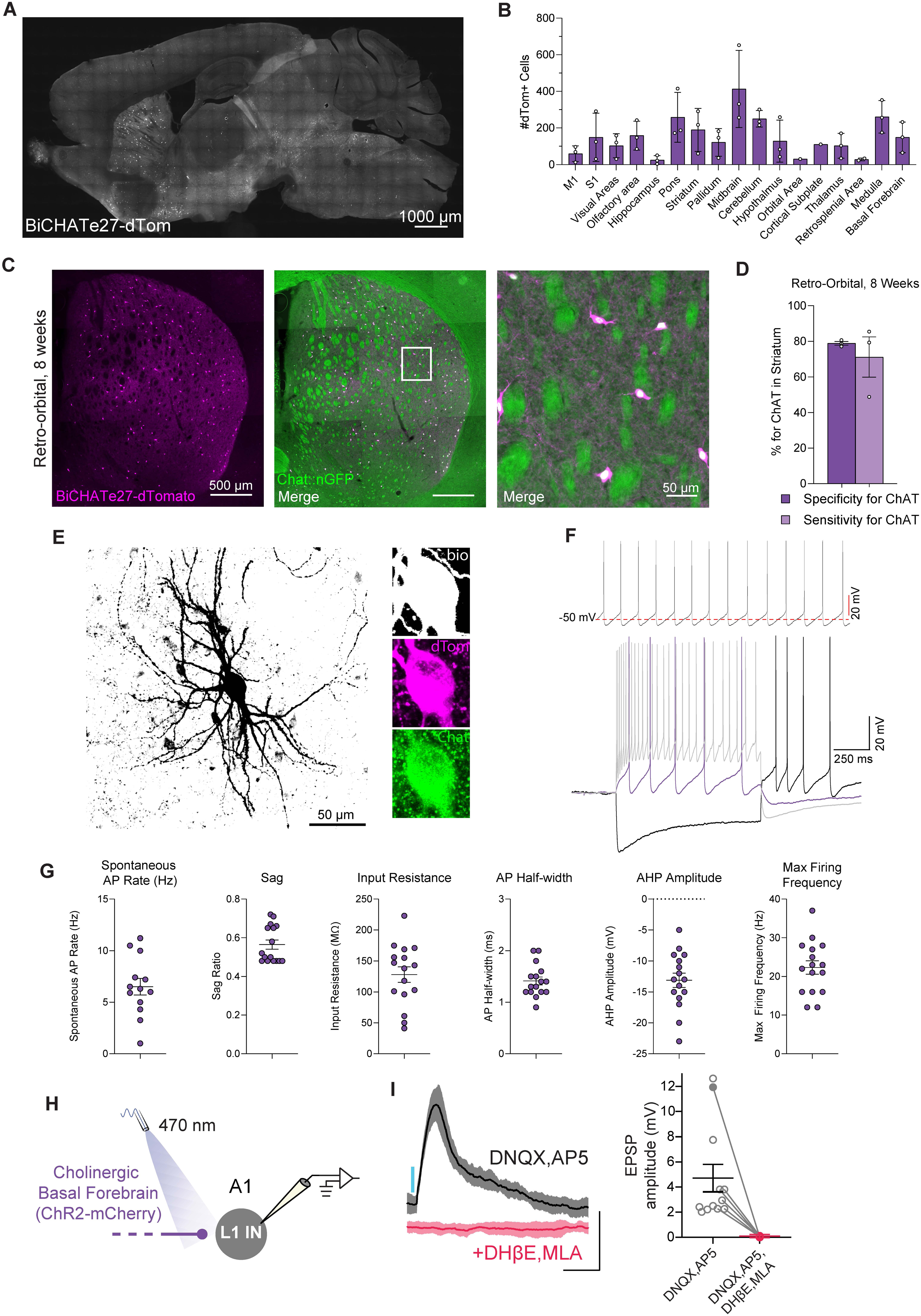
Enhancer-based viral targeting of CHAT-positive neurons in mice. (A) Representative para-sagittal sections showing the distribution of dTomato-positive cells resulting from expression controlled by the BiCHATe27 enhancer within the mouse brain, three weeks after RO injection of 2.0E+11 total vg/mouse at 4 weeks. Scale bar is indicated in the figure. (B) Histograms showing the number of dTomato-positive cells under the control of the BiCHATe27 enhancer in the brain areas indicated. Data from 5 sagittal sections from 5 different medio-lateral coordinates were collected, to cover the majority of brain regions. Each individual datapoint represents a distinct biological replicate; N=3 mice. Error bars represent SEM. (C) Representative image showing the expression of dTomato reporter *(in magenta*) under the control of the BiCHATe27 enhancer, and the Chat-cre dependent expression of GFP reporter (*in green*) in striatum of the mouse brain, following retro-orbital AAV injection at 8 weeks. The dTomato-positive cells co-expressing the marker (merge) are labeled *in white*. *Right panel*: zoomed-in image from the left panel. Scale bars are indicated in the figure. (D) Bar graphs showing the percentage specificity and sensitivity of the BiCHATe27 enhancer in targeting CHAT-positive neurons in the striatum. N=3 mice. Error bars represent SEM. (E) Biocytin filled reconstruction of BiCHATe27 infected cell recorded in mouse striatum. *Inset*: Reconstructed cell body in biocytin (*white*), BiCHATe27-dTomato (*magenta*), and CHAT immunostaining (*green*). Scale bar is indicated in the figure. (F) *Top panel:* Example trace of spontaneous action potential firing pattern during cell-attached patch clamp of BiCHATe27-dTomato infected cell. *Bottom panel:* Intrinsic firing properties recorded from BiCHATe27-dTomato infected cell, electrophysiological membrane and action potential firing responses at hyperpolarized (*black*), threshold (*red*), and maximum firing depolarization (*gray*) current steps. (G) Intrinsic membrane and firing properties of BiCHATe27 infected cells recorded in mouse striatum. *n* = 15-16 cells across N=3 mice. Error bars represent SEM. (H) Schematic of whole-cell current clamp recordings in layer 1 cortical interneurons (L1 INs) of primary auditory cortex (A1) in response to optogenetic stimulation of BiCHATe27-ChR2-mCherry axons. (I) *Left panel*: Representative (mean ± SD) excitatory postsynaptic potentials (EPSPs) from a L1 IN in response to a 5 ms blue light pulse delivered with a 470 nm LED (∼14 mW/mm2, 0.1 Hz). Responses were recorded in the presence of 20 μM DNQX and 50 μM AP5, AMPA-R and NMDA-R blockers, respectively, and eliminated in the presence of nAChR blockers 10 μM DHβE and 10 μM MLA (*red traces*). Scale bars: 5 mV, 200 ms. *Right panel*: Average EPSP amplitude in the presence of DNQX and AP5 (*n* = 12 cells) and in the presence of DNQX, AP5, DHβE and MLA (*n* = 6 cells), N=4 mice. Error bar represents SEM. Filled circle corresponds to the representative cell shown in the left panel.

To test the cell type-specificity of the BiCHATe27 enhancer, we further quantified the colocalization of BiCHATe27-dTomato cells in the caudoputamen with the cholinergic marker CHAT (Figure 5C). Following RO injection at both 4 weeks and 8 weeks of the AAV-BiCHATe27-dTomato in Chat-cre::INTACT mice, we observed strong dTomato labeling in the mouse striatum (Figures 5C and S8A_ii_). A high proportion of these BiCHATe27-dTomato cells co-express CHAT, as highlighted by the high percent average specificity for CHAT in both these conditions (average specificity of 83.6% ± 2.4% for RO4w and 78.8% ± 1.8% for RO8w). In addition, the high percent sensitivity shows the high efficiency of the BiCHATe27 enhancer in labeling all CHAT-positive neurons in the whole caudoputamen by RO injections at both ages (average sensitivity of 70.8% ± 21.9% for RO4w and 71.2% ± 19.6% for RO8w) (Figures 5D, S8A_ii_ and S8B_ii_). On the other hand, perinatal ICV injections of AAV-BiCHATe27-dTomato, despite maintaining a strong specificity for Chat neurons, is not the ideal method to thoroughly label all cholinergic neurons in this brain region, as indicated by the low percent sensitivity with this condition (Figures S8A_i_ and S8B_i_). Finally, intracranial injections in the caudoputamen show good colocalization of BiCHATe27-dTomato cells with the CHAT marker (Figures S8A_iii_ and S8B_iii_), demonstrating the reliability of the BiCHATe27 enhancer in labeling cholinergic neurons in the striatal regions with high specificity and sensitivity across multiple conditions.

To further characterize striatal cells labeled by BiCHATe27, we performed electrophysiological recordings in acute brain slices prepared from adult mice that received intracranial injection of AAV-BiCHATe27-tdTomato, followed by 2 weeks of incubation. Anatomical recovery of biocytin-filled BiCHATe27-dTomato expressing cells consistently revealed large aspiny neurons that co-labeled with CHAT (Figure 5E). Striatal BiCHATe27-dTomato expressing cells consistently exhibited spontaneous firing around 6Hz at rest with relatively slow (∼1.4 ms half-width) individual action potentials followed by large slow after hyperpolarizations (AHPs). Long negative current injections promoted an initial hyperpolarization followed by prominent sag in membrane potential, characteristic of the activation of an Ih conductance. BiCHATe27-tdTomato-expressing cells exhibited low maximal firing rates with strong AP accommodation in response to depolarizing current steps (Figures 5F and 5G). All these physiological properties reflect hallmark features of striatal cholinergic interneurons that readily distinguish them from surrounding medium spiny projection neurons and local circuit interneurons of the striatum.^40,41^

Cholinergic neurons in the brain are not restricted in striatal regions but can be found in other brain areas such as the basal forebrain, the brainstem or the thalamus. We therefore tested if our BiCHATe27 enhancer can be used to target cholinergic neurons residing in these regions. Neurons in the cholinergic basal forebrain form monosynaptic contacts with interneurons in the superficial layers of the auditory cortex^43^ that robustly respond to ACh via nicotinic acetylcholine receptors (nAChRs).^43–45^ We thus intracranially injected in the mouse basal forebrain AAV-BiCHATe27-ChR2-mCherry (Figure 5H) and first confirmed the specificity of our enhancer in labeling Chat neurons also in this region. We quantified the percent specificity of BiCHATe27 - mCherry cells for CHAT and showed that both cell soma located in the basal forebrain and mCherry-positive axons in the primary auditory cortex (A1) highly colocalize with this marker (Figures S8C and S8D). We then obtained brain slices containing A1 and recorded excitatory postsynaptic potentials (EPSPs) from A1 L1 interneurons in response to photo-stimulation of cholinergic basal forebrain BiCHATe27-ChR2-mCherry-positive axons (Figure 5H). These optogenetically-evoked EPSPs were recorded in the presence of the glutamatergic receptor antagonists DNQX and AP5 and were completely eliminated by the application of the nAChR blockers DhβE and MLA, consistent with direct, monosynaptic release of ACh (Figure 5I).

In conclusion, we showed that the BiCHATe27 enhancer can be used as a highly efficient tool to effectively and specifically target cholinergic neurons across multiple brain areas and with multiple methods of injections.

### Enhancer-based viral targeting of GABAergic and cholinergic neurons in non-human primates and human tissue

Although our cell-type specific enhancers were selected based on mouse genomic data, species conservation was considered when choosing the sequences. To evaluate if these enhancer-based viral tools can be used across species, we tested them in non-human primates (NHPs) and human brain slices. Specifically, AAVs carrying enhancer-dTomato or enhancer-ChR2-mCherry expression cassettes were delivered intracranially into the rhesus macaque cortex, hippocampus, or striatum (Figures 6A-G). 6-8 weeks following AAV delivery, NHP brains were extracted for *ex vivo* acute brain slice electrophysiological interrogation and tissue was subsequently drop-fixed and processed for combined anatomical recovery of recorded cells and immunohistochemistry (IHC) for appropriate markers. The dTomato-expressing cells infected with BiPVe3 or BiPVe4 AAVs typically co-expressed the PV marker in cortex and hippocampus respectively and displayed prototypical fast spiking profiles with short duration action potentials (APs) that minimally accommodated or broadened at sustained high maximal firing rates (Figures 6A and 6B; Table 1). Notably, NHP hippocampal BiPVe4-dTomato expressing cells were frequently observed to display axonal cartridge structure similar to rodent cortical chandelier cells (Figure 6Bi, Biii). Hippocampal BiSSTe10-dTomato expressing cells typically displayed horizontally oriented soma and dendrites within *stratum oriens* and showed high colocalization with the SST marker (Figure 6C). BiSSTe10 labeled hippocampal cells had wider, accommodating APs, and lower maximal sustained firing frequencies than BiPVe3 and BiPVe4 labeled subsets (Figure 6C, Table 1).

**Figure 6.**
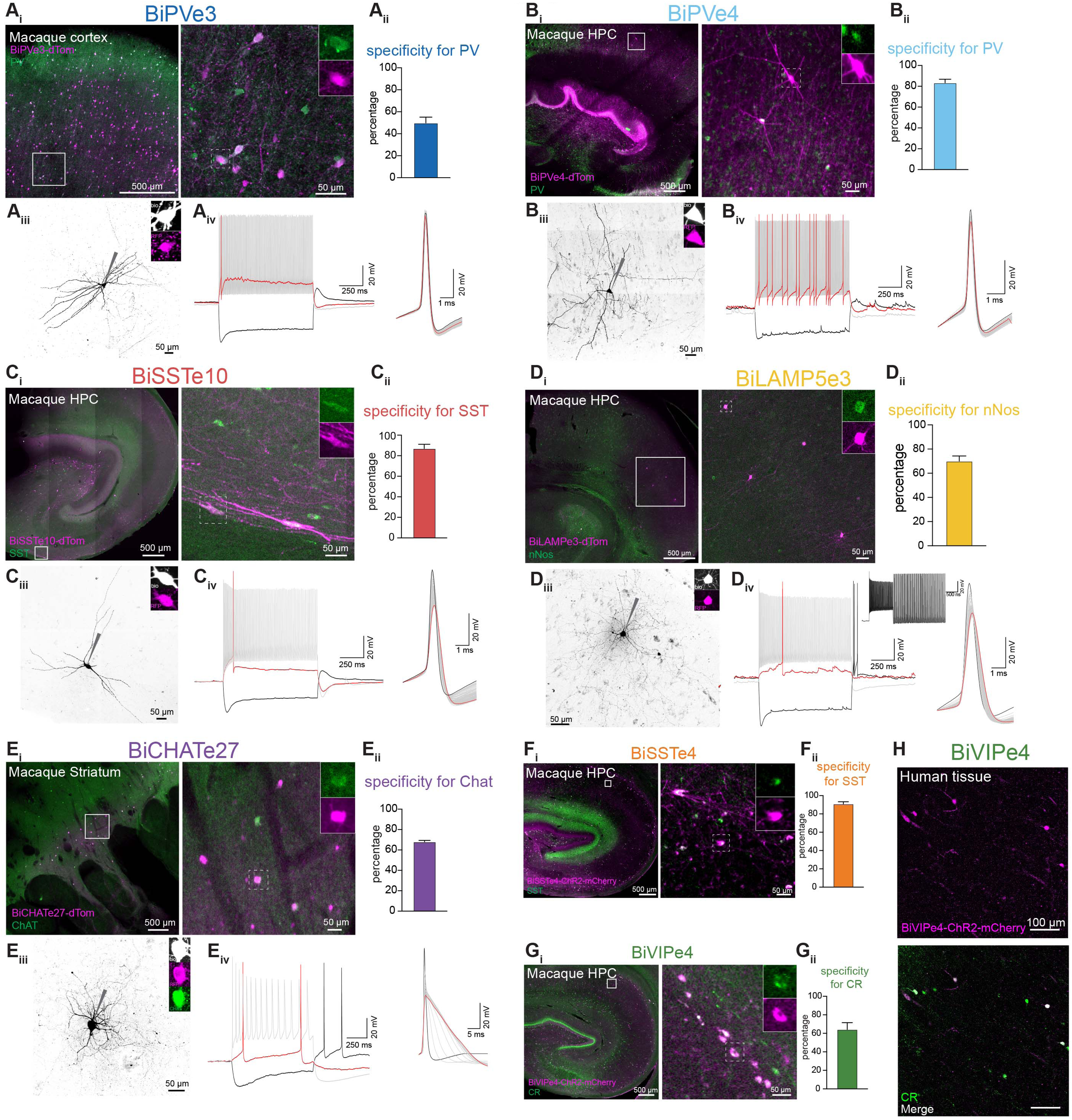
Validation of enhancer-based viral tools in non-human primates and human tissue. Validation of (A) BiPVe3, (B) BiPVe4, (C) BiSSTe10, (D) BiLAMP5e3, (E) BiCHATe27, (F) BiSSTe4, and (G) BiVIPe4 in rhesus macaque. (A-Gi) Overview images of reporter expression (*magenta*) colocalized with immunohistochemical (IHC) staining of target molecular markers (*green*). The scale bar is indicated in the figure. nNOS: neuronal nitric oxide synthase, CR: calretinin, HPC: hippocampus. (A-Gii) Percentage specificity of rAAV transduced cells for the target cell type-specific markers. BiPVe3: 12 sections from 1 animal, BiPVe4: 25 sections from 1 animal, BiSSTe10: 8 sections from 2 animals, BiLAMP5e3: 13 sections from 2 animals, BiCHATe27: 20 sections from 2 animals, BiSSTe4: 5 sections from 1 animal, BiVIPe4: 5 sections from 2 animals. Error bars represent SEM. (A-Eiii) Biocytin filled cell-reconstructions of rAAV transduced cells. (*Inset*) Reconstructed cell body in biocytin (*white*) and reporter expression (*magenta*). (A-Eiv) Intrinsic firing properties recorded from rAAV transduced cells. (*Left*) Electrophysiological membrane and action potential firing responses at hyperpolarized (*black*), threshold (*red*), and maximum firing depolarization (*gray*) current steps. (*Right*) Spike shape and accommodation of rAAV transduced cells depicting initial (*black*), intermediate (*gray*), and final (*red*) spikes during maximum firing trains. Inset in Div shows persistent firing of BiLAMP5 cells after the end of depolarizing pulse. (H) Validation of BiVIPe4 enhancer activity in human tissue. Representative images of human tissue sections transduced with AAV-BiVIPe4-ChR2-mCherry, co-stained with Calretinin (CR). The scale bar is indicated in the figure.

**Table 1.** Electrophysiological Intrinsic Membrane and Firing Properties of Targeted Macaque Cells. *n,* number of cells*. RMP*, Resting membrane potential (mV). *sAP freq*, spontaneous Action Potential frequency (Hz). *tau*, membrane time constant. *Rin*, input resistance (MΩ). Sag ratio, hyperpolarization sag. *AP Threshold*, action potential threshold (mV). *AP Half-width*, action potential half-width (ms). *Max firing*, maximum firing frequency (Hz). *AHP amp*, after-hyperpolarization amplitude (mV). *ISI accommodation ratio*, inter-spike interval accommodation ratio.

|  | **BiPVe3** | **BiPVEe4** | **BiSSTe10** | **BiCHATe27** | **BiLAMP5e3** |
| --- | --- | --- | --- | --- | --- |
| **Target cell** | PV Basket | PV Chandelier | non-Martinotti SST | Chat | LAMP5 IN |
| **# cells** | 8-17 | 17-19 | 23-32 | 14-17 | 11-14 |
| **RMP (mV)** | -63.25 ± 2.55 | -59.09 ± 0.93 | NA | -48.07 ± 1.43 | -66.4 ± 4.45 |
| **sAPs (Hz)** | NA | NA | NA | 3.47 ± 0.34 | 4.06 ± 1.10 |
| **tau** | 11.51 ± 2.59 | 15.59 ± 2.01 | 20.41 ± 1.54 | 57.67 ± 10.50 | 16.86 ± 1.31 |
| **Rin (mOhms)** | 114.76 ± 12.06 | 98.03 ± 15.67 | 143.22 ± 12.56 | 120.20 ± 13.16 | 145.64 ± 19.25 |
| **Sag Ratio** | 0.57 ± 0.04 | 0.79 ± 0.03 | 0.59 ± 0.03 | 0.50 ± 0.03 | 0.72 ± 0.03 |
| **AP Threshold (mV)** | -44.08 ± 1.01 | -44.47 ± 1.08 | -49.40 ± 1.33 | -37.03 ± 0.97 | -41.45 ± 1.05 |
| **AP Half-Width (ms)** | 0.41 ± 0.02 | 0.43 ± 0.05 | 0.73 ± 0.06 | 2.69 ± 0.25 | 0.72 ± 0.03 |
| **Max Firing (Hz)** | 177.33 ± 14.55 | 235.94 ± 21.57 | 124.23 ± 10.81 | 16.20 ± 2.86 | 91.71 ± 3.76 |
| **AHP Amplitude (mV)** | -19.51 ± 1.09 | -16.91 ± 1.07 | NA | -16.27 ± 0.98 | -8.38 ± 3.37 |
| **ISI accomodation Ratio** | 1.37 ± 0.05 | 1.37 ± 0.08 | 1.75 ± 0.11 | 5.52 ± 0.91 | 2.42 ± 0.157 |

BiLAMP5e3 in the hippocampus showed high activity in a sparse population of cells with Neurogliaform morphology, co-expressing the marker nNos, and also exhibiting regular firing properties with broader accommodating action potentials (Figure 6D, Table 1). Consistent with prior observations in macaque neocortex,^46^ the putative Neurogliaform cells labeled by BiLAMP5e3 showed limited late-spiking behavior relative to their rodent counterparts (delay to first spike at threshold 145 +/- 43 ms, n=14 cells, Figure 4Q and Table S1 for rodent counterpart). However, NHP BiLAMP5e3 labeled cells often entered a persistent firing mode characteristic of both rodent and human Neurogliaform cells (10/14 cells; Figure 6D_iv_ and 4R for rodent counterpart).^47^

NHP striatal labeling with BiCHATe27 revealed a sparse population of large cells that frequently co-labeled with CHAT (Figure 6E). As in rodent, and as typical of striatal cholinergic interneurons (also called TANs for tonically active neurons), these cells exhibited tonic AP firing at rest with relatively wide half-widths and significant accommodation and broadening upon depolarization induced sustained firing (Figure 6 D_iv_, Table 1). In the hippocampus, BiSSTe4- ChR2-mCherry labeled cells typically co-expressed SST and provided strong light-driven GABAergic inhibitory input to local pyramidal cells (Figure 6F and S9A). In contrast, BiVIPe4 - ChR2-mCherry labeled hippocampal cells that showed good colocalization with the marker calretinin and provided limited GABAergic input to local pyramidal cells, as expected for disinhibitory interneuron-selective VIP interneurons (Figure 6G and S9B). In sum, our combined IHC and electrophysiological profiling of cells labeled by our AAV-enhancer tools in NHP strongly support maintained selectivity for the intended neuronal subpopulations of the enhancer tested, providing a watershed opportunity for functional cell-type specific microcircuit interrogation across evolution.

To further expand the usage of our enhancer-based viral tools, we tested the VIP enhancer, BiVIPe4, in human brain tissue. Surgically resected human tissue collected from the temporal lobe of two patients was sectioned and incubated with rAAVs carrying the BiVIPe4-ChR2-mCherry expression cassette. Six days after transduction, we evaluated the specificity of BiVIPe4 in human tissues using calretinin (CR) as a marker for human VIP interneurons. We observed that cells expressing mCherry co-express CR (Figure 6H), suggesting BiVIPe4 enhancer remains selective for VIP cells in human tissue.

## DISCUSSION

The cerebral cortex contains a wide range of inhibitory interneuron subtypes which are central to the gating of information in this structure. While previous efforts have been aimed at understanding this diversity at the level of subtypes, the ability to target and manipulate distinct populations provides the clearest path to understanding their contribution to computation. In the absence of specific drivers for individual subtypes, the inability to target them limits our understanding of the functional contributions of each different interneuron class. Historically, targeting of such populations has been achieved by Cre- and Flp-based transgenic animals. However, these genetic lines require lengthy breeding processes and are not available in many species. Here by identifying enhancers that allow for the selective targeting of all major interneuron types, we provide a versatile toolset for researchers to explore the function of different interneuron subtypes. We use these enhancers in the context of recombinant AAVs, which are easy to package, have low immunogenicity, and can easily be used to deliver transgene in many species, including rodents and primates. Thus, this approach is both cost effective and efficient. In recent years, we and others have established a number of enhancers, that when incorporated into rAAV vectors, can restrict the transgene expression to certain neuronal cell types.^5–11^ We have previously published the Dlx enhancer, a pan-interneuron enhancer, and the E2 enhancer, a pan PV-enhancer.^5,6^ In this study, we expanded our efforts, aiming to identify enhancers for all interneuron cardinal types. We presented a toolkit to target all major interneuron cardinal types in the cerebral cortex. These enhancer-rAAV tools allow the targeting of two major PV subtypes, two major SST subtypes, as well as VIP, LAMP5 and cholinergic neurons. Importantly, while we made considerable efforts to identify the target population for each enhancer, it is only through the collective use of these tools by the field as a whole that the precise specificity of each of these enhancers will be confirmed. For instance, while we refer to the BiSSTe4 enhancer as specific to Martinotti cells, future work may indicate its specificity is in fact better described as an infragranular SST enhancer. It is also important to note that there has been a significant amount of effort by the community to develop enhancer-based cell targeting tools in recent years.^5–11^ In this study, we characterize the enhancer-rAAVs by assessing co-expression of cell-type specific marker genes, as well as the morphological and physiological properties of labeled cells. Nonetheless, each group uses somewhat different methods to characterize the enhancer tools they develop. While others may use different metrics and methods, such as single-nucleus RNA-seq, to evaluate their tools, each method has its strengths and limitations. Thus, a parallel evaluation of tools developed by different groups would be beneficial to the community.

To demonstrate the versatility of the cell-type specific enhancers we identified, we paired these enhancer elements with different payloads. We demonstrate that these tools provide the means to label specific interneuron types (fluorescent reporter), observe their neuronal activity (GCaMP), as well as to manipulate specific GABAergic and cholinergic neuron populations (optogenetic effector). For example, *in vivo* optogenetic activation of cells labeled by our AAV-enhancer tools can be used to study the recruitment of specific GABAergic interneurons for cortical function. The inhibitory effect of the activation of each interneuron subtype differs across them (Figure S10A), and varies across cortical layers. Analysis of the population activity shows that activation of LAMP5 cells results in a strong inhibition of cells in both shallow and deep layers, while activation of basket cells has a stronger effect on cells in deep cortical layers (Figure S10B). This is consistent with previous results on the anatomical projections of each type of interneuron subtype.

Beyond these examples, the practical use of these enhancers is potentially much broader. They can be paired with any payload to express various effectors in specific cell populations. For example, these enhancers can be used to express the helper proteins of mono-synaptic rabies for circuit tracing, or in conjunction with Cre- or Flip-based genetic lines for intersectional labeling. Moreover, these enhancers can also be used to deliver therapeutic proteins for gene therapy.

A further advantage of enhancers is that when chosen judiciously with regards to cross-species conservation, a single enhancer can be used in a variety of species. Although we used mouse interneuron single cell chromatin (i.e. ATAC-seq) data to identify cell-type specific enhancers, by selecting those whose sequence and accessibility is conserved, we find that they are often reliable for targeting similar cell types in other species, including human and NHPs. Indeed, by testing their activity in non-human primates and human brain slices, these enhancer elements often showed conserved targeting patterns in primate tissues. As such, our toolkit can be used to access specific cell types across species *in vivo* and *ex vivo*. This expands the applicability of our interneuron enhancer-rAAV tools and provides the potential to be used in therapeutic context for modulating inhibition and signaling.

In addition to cross-species uses, the same enhancer can often be used to target homologous cell types in different CNS structures. With regards to interneurons, the cerebral cortex and hippocampus represent two such regions. Indeed, the interneurons in these areas share the same developmental origins and single cell transcriptomic study suggests that most interneuron cell types have shared gene expression patterns between these two regions.^1^ Therefore, it is likely that these enhancer elements are regulated by shared gene regulatory networks in cortex and hippocampus. As a consequence, these vectors can be used to label corresponding cell types across both hippocampus and cortex, both in mice and NHPs. Indeed, in testing specific enhancers in regions other than the cortex, we find they label homologous or analogous cell types. A recent publication, for instance, uses the BiPVe4 enhancer and carefully performed IHC and electrophysiological validations to analyze chandelier cell types in the hippocampus in detail.^17^ In the future, such efforts by investigators in the field will be an essential component for validating the use of both our enhancers and others looking for similar regulatory elements. Similarly, we also observe that homologous cell types in different areas such as the striatum and the basal forebrain can be efficiently targeted by the BiCHATe27 enhancer (Figures 5C and S8).

One technical point of note, when characterizing the specificity of the enhancer-rAAVs, we analyzed the native fluorescent (dTomato) signal. Many studies tend to visualize reporter expression by immunostaining. We compared the signal from native dTomato fluorescent and that amplified by anti-dTomato antibody. We found that immunostaining improved the signal intensity and showed better visualization of neuronal processes (data not shown). However, upon quantification, the total number of dTomato-positive cells in the brain region of interest do not show significant differences regardless of whether native or antibody-enhanced staining is used (Figure S10C).

Most of the enhancer discovery studies tend to focus on one delivery method. In our study, we realized that the specificity of enhancer-rAAV can be affected by many factors, including viral load, injection age, and injection route. Here, we presented a systematic characterization of our enhancer-rAAV tools. We evaluated their performance under each of the four conditions: RO injection in 4-week old mice, RO injection in 8-week old mice, ICV injection in P0/P1 mice and stereotactic injection in adult mice or pups. We showed that delivery conditions can dramatically alter the specificity and sensitivity of enhancer-rAAV vectors. For example, of the two SST enhancers, the non-Martinotti SST targeting enhancer (BiSSTe10) performed best when delivered systemically in 4-week old mice (Figure 3E). On the other hand, the Martinotti SST targeting enhancer (BiSSTe4) worked most optimally when delivered by ICV in P0/P1 mice (Figure 3F). However, at the conditions tested, both enhancers performed poorly when injected by intracranial stereotaxic injections in the cortex (Figure 4). In this regard, we found that intracranial stereotactic injection is most sensitive to viral load, since by this injection method, the multiplicity of infection (MOI) will be very high, especially near the site of injection. We have found that the specificity of enhancers for the expected target cell types tend to decrease as we increase the viral load (data not shown). We thus recommend users to carefully scale viral vectors and test multiple injection conditions, to evaluate the labeling specificity and efficacy before performing experiments.

A further area where enhancers can improve access to the study of specific interneurons is with regard to development. In this regard, Parvalbumin (PV) interneurons are critical for the maturation of sensory circuits^48,49^ and are known to be dysfunctional in various neurodevelopmental disorders^50^. However, studying these neurons during early development has been challenging due to the late expression of endogenous parvalbumin. As a result, the field has resorted to complex intersectional genetic strategies^51^ or alternative transgenic lines with partial specificity or sensitivity^52^. To our knowledge, the BiPVe3 enhancer represents the first method that allows to target PV basket cells with high specificity (as high as 90.26% ± 5.65% in L4) and sensitivity (81.55% ± 15.61 across L2-6) even at early postnatal ages, when other tools fail to do so (Figure S2A_iv_ and S2C_iv_).

To facilitate best practices, we provide a standard operating procedure (Table 2) to allow users to choose the best delivery route based on their needs. Even so, we recommend users to test a series of different viral loads to optimize their approach depending on the enhancer, the titer and the payload being delivered. With regards to the latter, the relationship between expression level and selectivity varies widely for different payloads. While the delivery of functional components such as fluorescent proteins, opsins and activity reporters can tolerate a slight degree of off-target expression, catalytic proteins such as recombinases often fail to have the desired specificity with even relatively minor levels of off-target expression. For example, we have not been able to drive specific expression of wild-type Cre with the cell-type specific enhancers we have previously identified, possibly due to the fact that Cre-dependent recombination is extremely sensitive to off-target expression.^5^ However, there have been reposts in which cell-type specific enhancers are used to drive the expression of Cre variants with lower affinity to their binding sites.^11^ Notably, the analyses presented in this study were conducted only in the brain regions of interest (cortex or striatum). Many cell types can be found across different brain regions. For instance, cholinergic neurons exist in both striatum and basal forebrain. However, when we delivered BiCHATe27 vectors by RO injection, we observed robust labeling with high specificity for cholinergic neurons in the striatum but more moderate labeling in the basal forebrain (Figure 5A and 5B). Yet, direct injection of the same vector into the basal forebrain resulted in highly specific labeling of cholinergic neurons (Figure S8C and S9D). Thus, our enhancer tools have the potential to function across brain regions. However, the optimal injection routes may vary and need to be determined on a case-by-case basis.

**Table 2.**
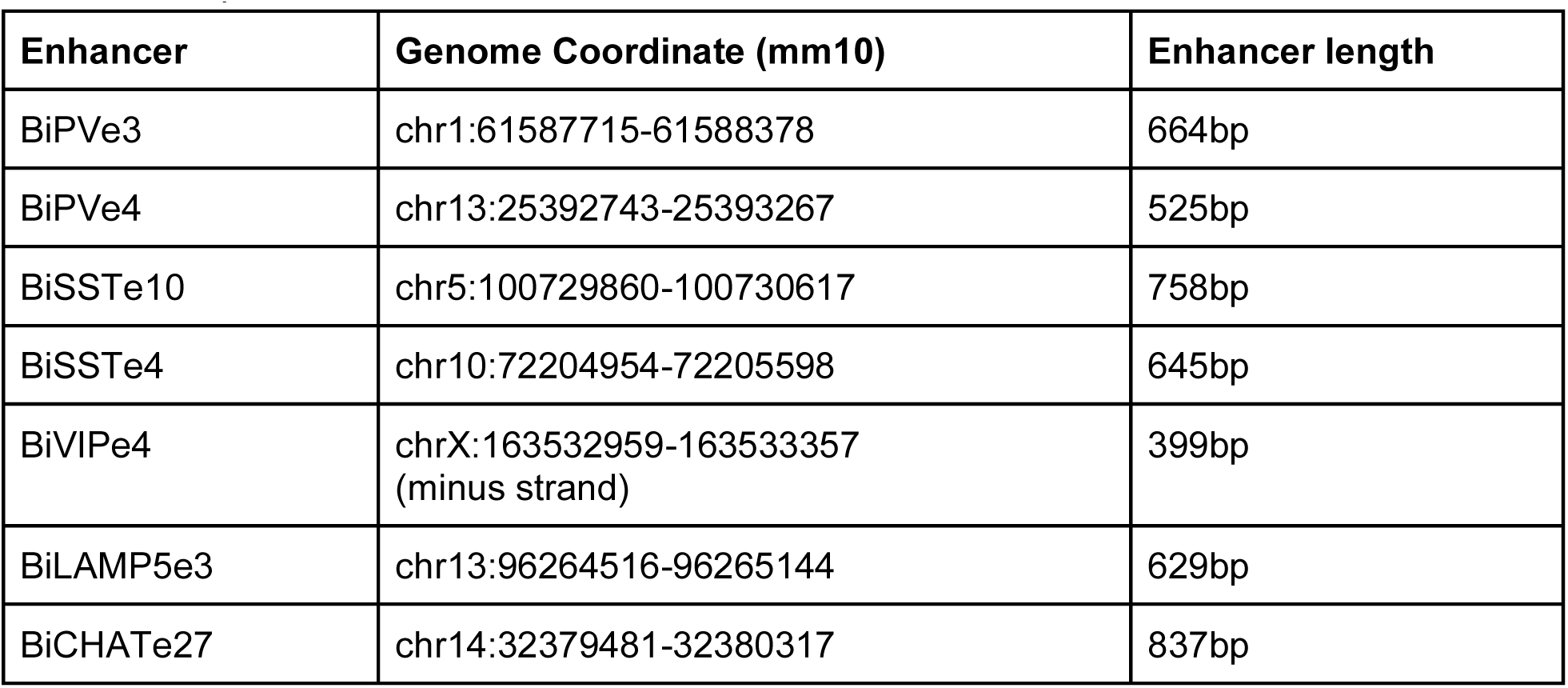
Summary of enhancer activity across injection conditions and ages. Table summarizing how, at the conditions indicated, each enhancer performs across different ages and routes of injection in mice. In green it’s indicated the condition that results in highest specificity and sensitivity of expression in the cell types of interest.

In this study, all the enhancer-rAAV constructs were tested with PHP.eB capsid. Each capsid has its own tropism bias. We have found that PHP.eB AAVs transduce PV cells more efficiently than other interneuron cell types. When we delivered PHP.eB AAVs carrying a GFP driven by a ubiquitous promoter (RO injection, 2E+11 viral genome/mouse), among all the interneuron cell types, we observed the strongest signal in PV cells and very minimal signal in VIP cells (data not shown). This is consistent with studies using single cell transcriptomic and spatial transcriptomic methods to characterize AAV tropisms.^53,54^ This also partially explains why the VIP enhancer (BiVIPe4) does not work well by RO injection, but it efficiently marks them following ICV injections. Thus, AAV tropism is another factor one needs to consider when using enhancer-based tools, as capsid choice can potentially alter the specificity of these enhancer constructs and the efficiency of neuronal labeling.

Of interest, the VIP enhancer we described here (BiVIPe4) is derived from a 1kb-sequence that was originally chosen. The packaging limit of AAVs is only about 4.7kb.^55^ Thus, identifying smaller enhancers will allow more space to accommodate larger payloads. In fact, the actual functional element of an enhancer is often short and contains binding sites for transcription factors. There have been reports using the enhancer bashing method to identify the core enhancer elements to be used with rAAV.^8,9^ We thus performed bashing experiments of the original VIP enhancer and identified a 399bp core sequence, BiVIPe4. This enhancer is located in the first intron of the gene *Grpr*, which is specifically expressed in VIP interneurons in the cortex. The gene, *Grpr*, is located on the minus strand and we thus designed BiVIPe4 to be on the minus strand. Interestingly, when we inverted the orientation of BiVIPe4 sequence on the rAAV vector, the vector lost specificity for VIP cells completely (data not shown), suggesting the orientation of an enhancer can be critical for its function. Thus, when designing enhancers, both size and orientation should be taken into consideration.

Taken together we present a collection of enhancers to target all major subclasses of cortical interneurons, as well as cholinergic populations, including large striatal interneurons. While the selectivity and specificity of each enhancer is unique and the optimal injection methods for using these enhancers differ in mode and timing of injection, each of these enhancers provides access to various populations with considerable selectivity. These enhancers add to the growing repertoire that we and other groups in this BRAIN Initiative consortium have identified for use in AAVs. Together this provides a broadened toolset for the targeting of specific cell types across mammalian species from rodents to primates.

## RESOURCE AVAILABILITY

### Lead contact

Further information and requests for resources and reagents should be directed to and will be fulfilled upon reasonable request by the lead contact.

### Materials availability

The plasmids newly created in this study are available from Addgene.

### Data and code availability

The genomic sequencing data related to this study are available on GEO (cortex: GSE165031 and GSE277242; striatum: GSE272181). This paper did not report original code. Any additional information required to reanalyze the data reported in this paper is available from the lead contact upon reasonable request.

## ACKNOWLEDGEMENTS

This effort was made possible by the contributions of the many groups (the majority of which are listed as authors). It was also greatly helped by the collaborative contributions of many of the investigators in the Brain Initiative consortium https://braininitiative.nih.gov/research/tools-and-technologies-brain-cells-and-circuits. Work in this paper was supported by Brain Initiative grants UF1MH130701 and UH3MH120096 to G.F. and Y.W., U24MH133236 to G.F., SNSF PostDoc.Mobility fellowship and EMBO Postdoctoral fellowship (ALTF 329-2022) to E.F.. This work was also supported by an NICHD Intramural Research Program (IRP) grant to CJM, and NIMH IRP grants to BBA and VAA. NHP work was carried out in collaboration with the NIH Comparative Brain Physiology Consortium (CBPC) at the NIH IRP. All enhancer-associate constructs described in this paper are available for scientific research from UCI Center for Neural Circuit Mapping through its U24MH13326 project https://cncm.som.uci.edu/brain-projects/ and Addgene https://www.addgene.org/collections/brain-initiative/. We would also like to thank Rima Pant for the help with the generation of one batch of BiPVe3 and BiPVe4 viruses.

---

**Table S1.** related to Figures 2-5. Table including all parameters obtained from whole-cell recording of cells labeled by AAV-enhancer tools, for all cells recorded. Each tab includes data from the enhancers indicated. Highlighted in yellow are the parameters plotted in the main figure.

**Figure S1.**
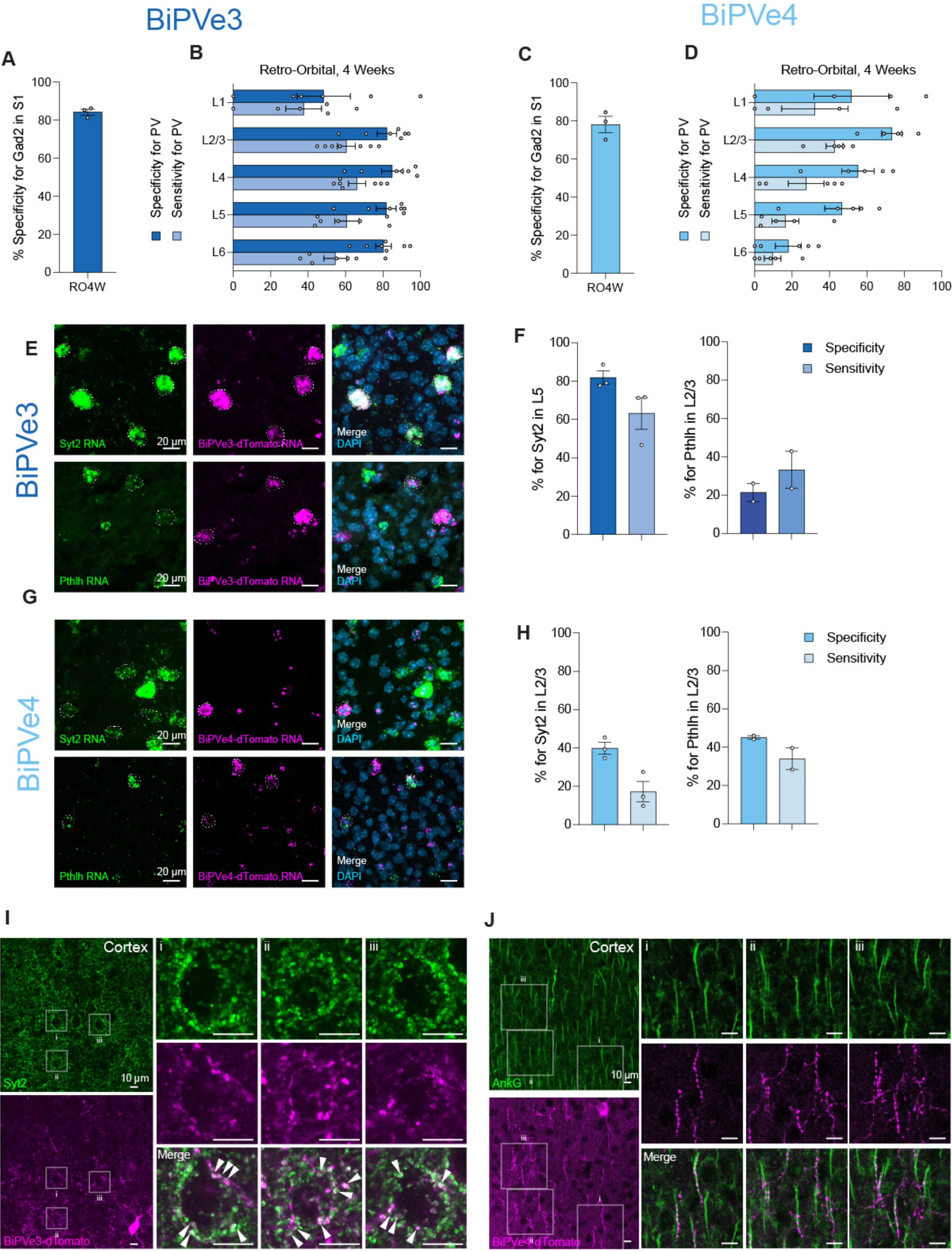
Validation of BiPVe3 and BiPVe4 cell type-specificity in the cortex, related to Figure 2. (A and C) Bar graphs showing the percentage specificity of BiPVe3 (A) or BiPVe4 (C) in targeting *Gad2*-positive neurons in S1. N=3 mice. Error bars represent SEM. (B and D) Bar graphs showing the percentage specificity of BiPVe3 (C) or BiPVe4 (D) in targeting PV-positive neurons, separately for each of the 6 cortical layers. N=7 and 6 mice, respectively. Error bars represent SEM. For BiPVe3 (B), the average # of cells that colocalize within each layer are L1: 0, L2/3: 104, L4: 202, L5: 245, L6: 201. For BiPVe4 (D), the average # of cells that colocalize within each layer are L1: 5, L2/3: 60, L4: 34, L5: 31, L6: 13. (E and G) Representative images of FISH for the marker genes *Syt2 (right panel*) and *Pthlh* (*left panel*) in mice injected with AAV-BiPVe3-dTomato (E) and AAV-BiPVe4-dTomato (G), in the cortical layers indicated. Regions of interest (ROI) are drawn to mark cells positive for the marker gene (*left panels*), for dTomato (*middle panels*), and cells co-expressing both RNAs (*right panels*). Scale bars are labeled in the figure. (F and H) Bar graphs showing the percentage specificity and sensitivity of the BiPVe3 (F) and BiPVe4 (H) enhancers in targeting cells expressing the markers indicated. Error bars represent SEM. (I and J) Images and zoom-ins (*i,ii,iii*) highlighting the colocalization of either BiPVe3-dTomato *(in magenta*) terminals with the basket-specific synaptic marker Syt2 (*in green*) (I) and BiPVe4-dTomato cartridges *(in magenta*) with the axon initial segment marker AnkG *(in green*) (J). Scale bars are indicated in the figure. The dTomato-positive terminals co-expressing the marker (merge) are labeled *in white.* Arrowheads highlight example terminals that show colocalization with the marker protein Syt2.

**Figure S2.**
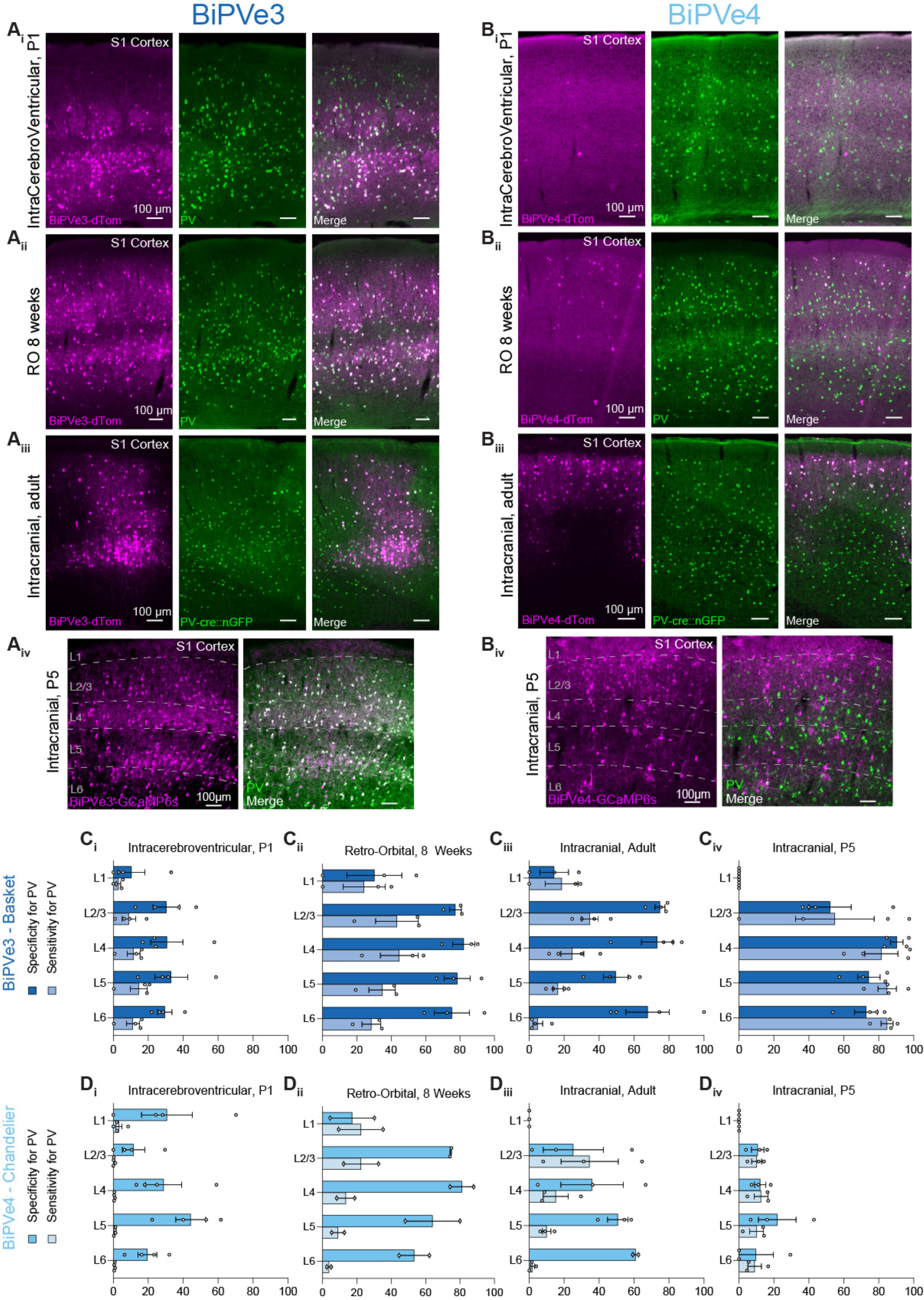
BiPVe3 and BiPVe4 enhancer specificity and sensitivity across injection conditions, related to Figure 2. (A and B) Representative columns showing the expression of dTomato reporter *(in magenta*) under the control of the BiPVe3 (A) or BiPVe4 (B) enhancers, and parvalbumin (*Pvalb*)-positive cells (*in green*) in the primary somatosensory cortex (S1) of the mouse brain, following ICV injection at P1 (*i)*, retro-orbital AAV injection at 8 weeks (*ii*) and intracranial, stereotactic injections in adult animals (*iii*) and P5 pups *(iv)*. dTomato-positive cells co-expressing the PV marker (merge) are labeled *in white*. Scale bars are indicated in the figure. (C and D) Bar graphs showing the percentage specificity and sensitivity of the BiPVe3 (C) or BiPVe4 (D) enhancers in targeting PV-positive neurons across the 6 cortical layers of S1, across the conditions indicated. N=2-4 mice. Error bars represent SEM. For BiPVe3 (C), the average # of cells that colocalize within each layer for each condition are as follows. For ICV, L1: 1, L2/3: 21, L4: 28, L5: 66, L6; 78. For RO8w, L1: 1, L2/3: 132, L4: 203, L5: 306, L6: 158. For Intracranial Adult, L1: 1, L2/3: 92, L4: 85, L6: 28. For BiPVe4 (D), the average # of cells that colocalize within each layer for each condition are as follows. For ICV, L1: 1, L2/3: 2, L4: 3, L5: 6, L6: 4. For RO8w, L1: 1, L2/3: 76, L4: 76, L5: 86, L6: 19. For Intracranial Adult, L1: 0, L2/3: 10, L4: 24, L5: 38, L6: 41.

**Figure S3.**
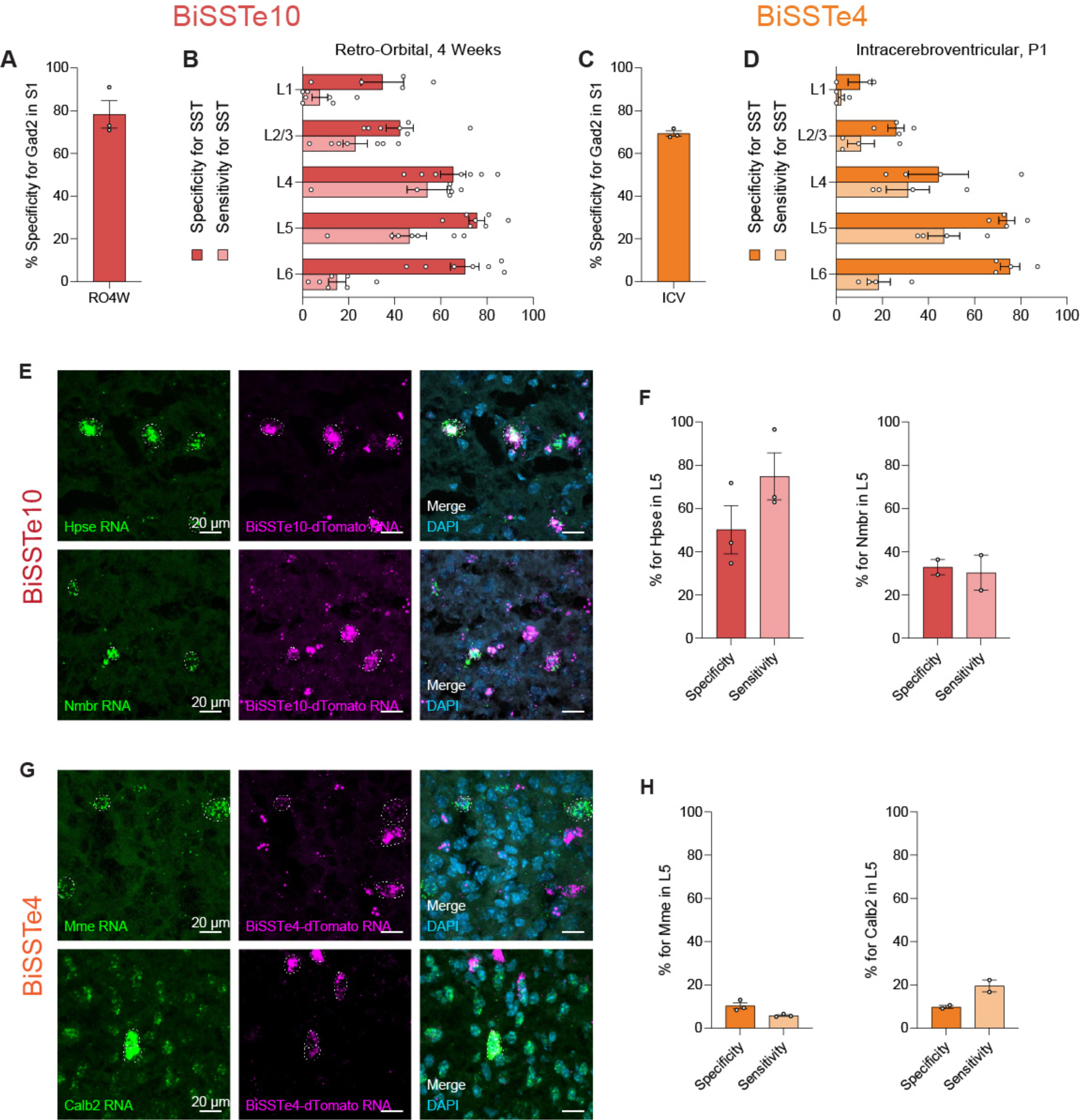
Validation of BiSSTe10 and BiSSTe4 cell type-specificity in the cortex, related to Figure 3. (A and C) Bar graphs showing the percentage specificity of BiSSTe10 (A) or BiSSTe4 (C) in targeting *Gad2*-positive neurons. N=3 mice. Error bars represent SEM. (B and D) Bar graphs showing the percentage specificity of BiSSTe10 (C) or BiSSTe4 (D) in targeting SST-positive neurons in S1, separately for each of the 6 cortical layers, across the conditions indicated. N=7 and 4 mice, respectively. Error bars represent SEM. For BiSSTe10 (B), the average # of cells that colocalize within each layer are L1: 1, L2/3: 47, L4: 80, L5: 162, L6: 25. For BiSSTe4 (D), the average # of cells that colocalize within each layer are L1: 0, L2/3: 9, L4: 20, L5: 99, L6: 79. (E and G) Representative images of FISH for the two SST subtype-specific markers indicated in mice injected with AAV-BiSSTe10-dTomato (E) and AAV-BiSSTe4-dTomato (G), in the cortical layers indicated. Scale bars are labeled in the figure. Regions of interest (ROI) are drawn to mark cells positive for marker genes (*left panels*), for dTomato (middle panels), and cells co-expressing both RNAs (*right panels*). Scale bars are labeled in the figure. (F and H) Bar graphs showing the percentage specificity and sensitivity of the BiSSTe10 (F) and BiSSTe4 (H) enhancers in targeting cells expressing the markers indicated. Error bars represent SEM.

**Figure S4.**
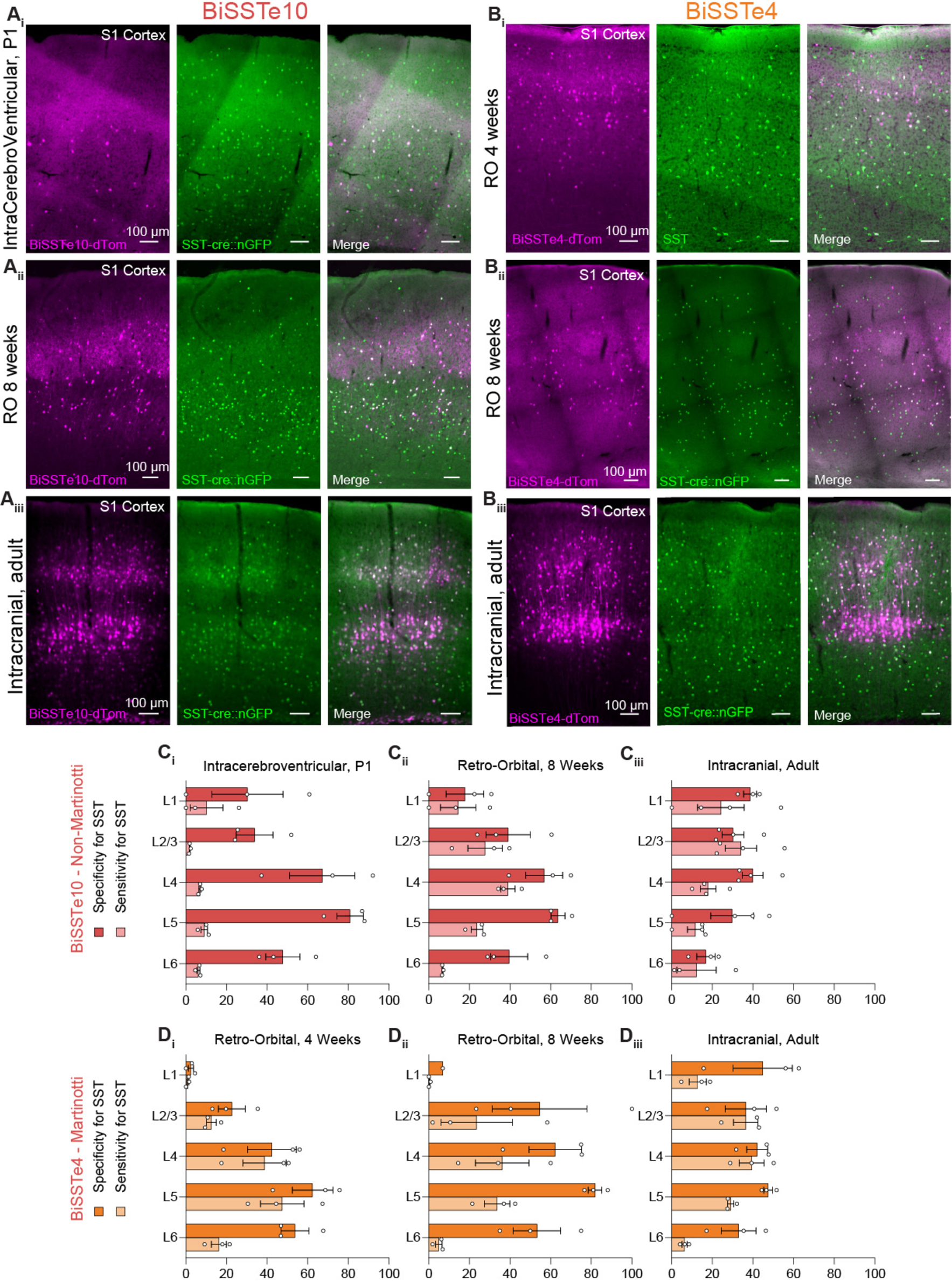
BiSSTe10 and BiSSTe4 enhancer specificity and sensitivity across injection conditions, related to Figure 3. (A and B) Representative columns showing the expression of dTomato reporter *(in magenta*) under the control of the BiSSTe10 (A) or BiSSTe4 (B) enhancers, and somatostatin (*Sst*)-positive cells (*in green*) in the primary somatosensory cortex (S1) of the mouse brain, following ICV injection at P1 (A*i*), retro-orbital AAV injection at 4 weeks (B*i*) RO at 8 weeks (*ii*) and intracranial, stereotactic injections (*iii*). dTomato-positive cells co-expressing the SST marker (merge) are labeled *in white*. Scale bars are indicated in the figure. (C and D) Bar graphs showing the percentage specificity and sensitivity of the BiSSTe10 (C) or BiSSTe4 (D) enhancers in targeting SST-positive neurons across the 6 cortical layers of S1, across the conditions indicated. N=3-4 mice. Error bars represent SEM. For BiSSTe10 (C), the average # of cells that colocalize within each layer for each condition are as follows. For ICV, L1: 1, L2/3: 5, L4: 15, L5: 63, L6: 46. For RO8w: L1: 1, L2/3: 85, L4: 149, L5: 156, L6: 28. For Local Injection Adult, L1: 2, L2/3: 71, L4: 43, L5: 85, L6: 116. For BiSSTe4 (D), the average # of cells that colocalize within each layer for each condition are as follows. For RO4w, L1: 1, L2/3: 28, L4: 72, L5: 205, L6: 189. For RO8w, L1: 1, L2/3: 37, L4: 95, L5: 169, L6: 17. For Local Injection Adult, L1: 1, L2/3: 85, L4: 71, L5: 112, L6: 40.

**Figure S5.**
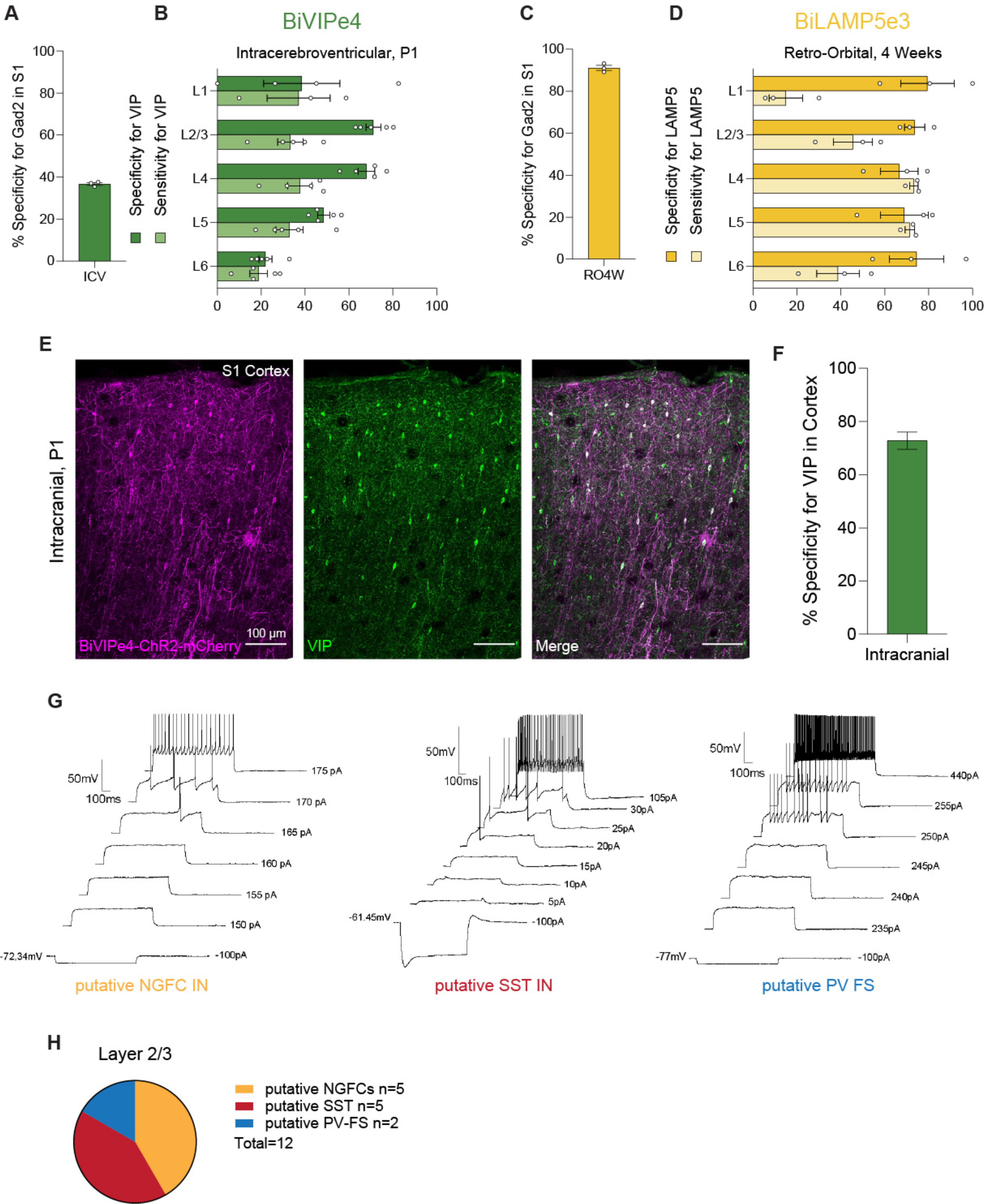
Validation of BiVIPe4 and BiLAMP5e3 cell type-specificity in the cortex, related to Figure 4. (A and C) Bar graphs showing the percentage specificity of BiVIPe4 (A) or BiLAMP5e3 (C) in targeting *Gad2*-positive neurons. N=3 mice. Error bars represent SEM. (B and D) Bar graphs showing the percentage specificity of BiVIPe4 (B) or BiLAMP5e3 (D) in targeting VIP- or LAMP5-positive neurons, separately for each of the 6 cortical layers in S1, across the conditions indicated. N= 4 and 3 mice, respectively. Error bars represent SEM. For BiVIPe4 (B), the average # of cells that colocalize within each layer are L1: 3, L2/3: 33, L4: 22, L5: 21, L6: 13. For BiLAMP5e3 (D), the average # of cells that colocalize within each layer are L1: 2, L2/3: 40, L4: 12, L5: 17, L6: 20. (E) Representative image showing the expression of ChR2-mCherry *(in magenta*) under the control of the BiVIPe4 enhancer, and VIP-positive cells (*in green*) in the primary somatosensory cortex (S1), following intracranial injection at P1. ChR2-mCherry-expressing cells co-expressing the VIP marker (merge) are labeled *in white*. Scale bar is indicated in the figure. (F). Bar graph showing the percentage specificity of BiVIPe4-ChR2-mCherry in targeting VIP-positive neurons. N=3 mice. Error bars represent SEM. (G) Examples of putative neurogliaform cells (NGFC), SST and PV-fast spiking (FS) cells responding to a 1-second increasing current injection. (H) Pie chart illustrating the proportion of putative NGFC/SST/PV-FS cells based on their firing properties, out of the 12 cells from 3 mice recorded in L2/3 of S1.

**Figure S6.**
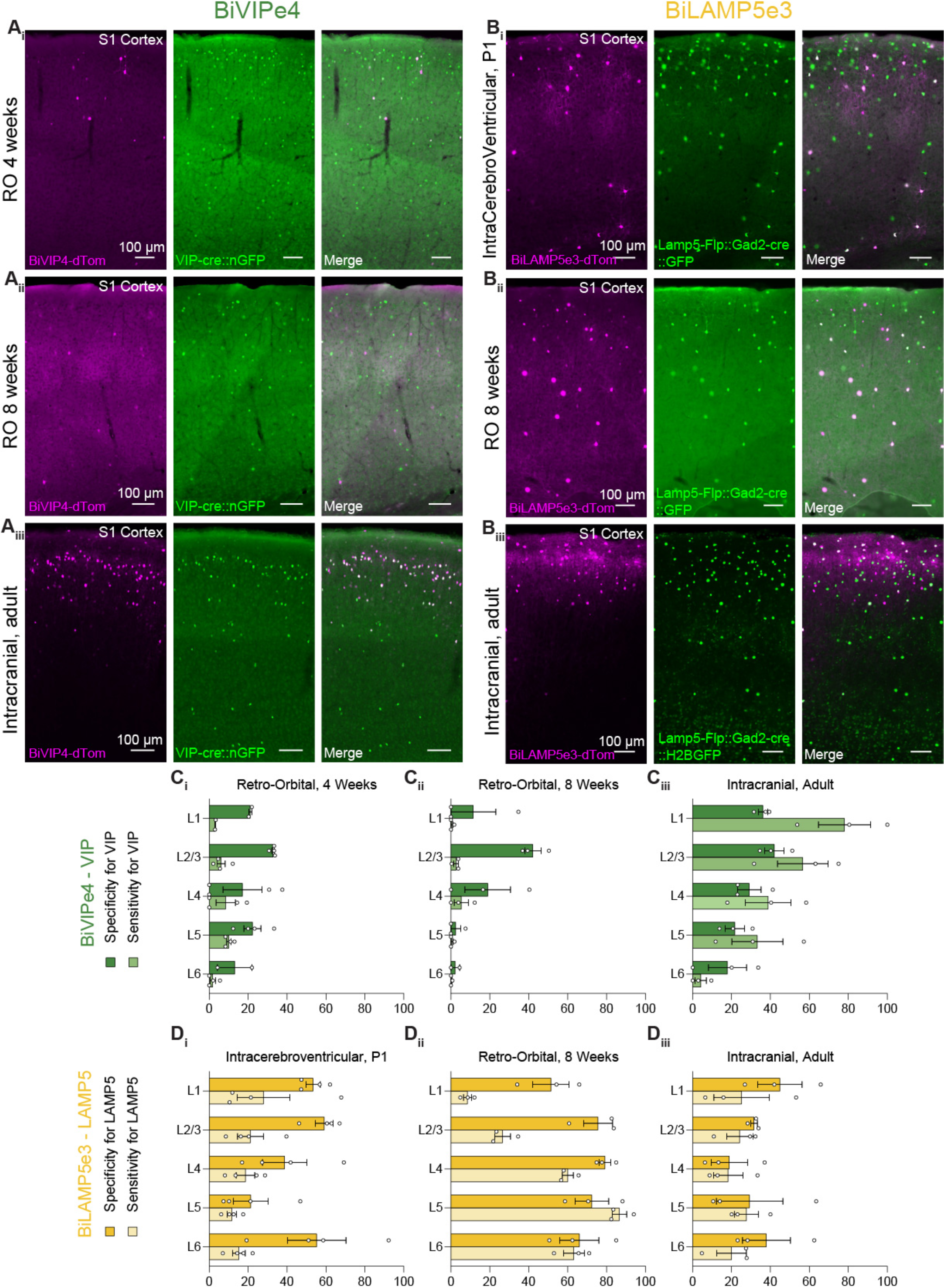
BiVIPe4 and BiLAMP5e3 enhancer specificity and sensitivity across injection conditions, related to Figure 4. (A and B) Representative columns showing the expression of dTomato reporter *(in magenta*) under the control of the BiVIPe4 (A) or BiLAMP5e3 (B) enhancers, and vasoactive intestinal peptide (*Vip*)-positive cells (A) or the genetically-expressed GFP reporter under the control of Lamp5-Flp and Gad2-cre (B) (*in green)* in the primary somatosensory cortex (S1) of the mouse brain, following ICV injection at P1 (B*i*), retro-orbital AAV injection at 4 weeks (A*i*) and 8 weeks (*ii*) and intracranial, stereotactic injections (*iii*). dTomato-positive cells co-expressing the markers indicated (merge) are labeled *in white*. Scale bars are indicated in the figure. (C and D) Bar graphs showing the percentage specificity and sensitivity of the BiVIPe4 (C) or BiLAMP5e3 (D) enhancers in targeting VIP- or LAMP5-positive neurons across the 6 cortical layers of S1, across the conditions indicated. N=4 and 3 mice, respectively. Error bars represent SEM. For BiVIPe4 (C), the average # of cells that colocalize within each layer for each condition are as follows. For RO4w, L1: 1, L2/3: 26, L4: 24, L5: 22, L6: 5. For RO8w, L1: 0, L2/3: 6, L4: 7, L5: 2, L6: 0. For Local Injection Adult, L1: 2, L2/3: 132, L4: 39, L5: 24, L6: 11. For BiLAMP5e3 (D), the average # of cells that colocalize within each layer for each condition are as follows. For ICV, L1: 2, L2/3: 34, L4: 11, L5: 7, L6: 13. For RO8w, L1: 2, L2/3: 113, L4: 105, L5: 78, L6: 72. For Local Injection Adult, L1: 1, L2/3: 23, L4: 6, L5: 21, L6: 29.

**Figure S7.**
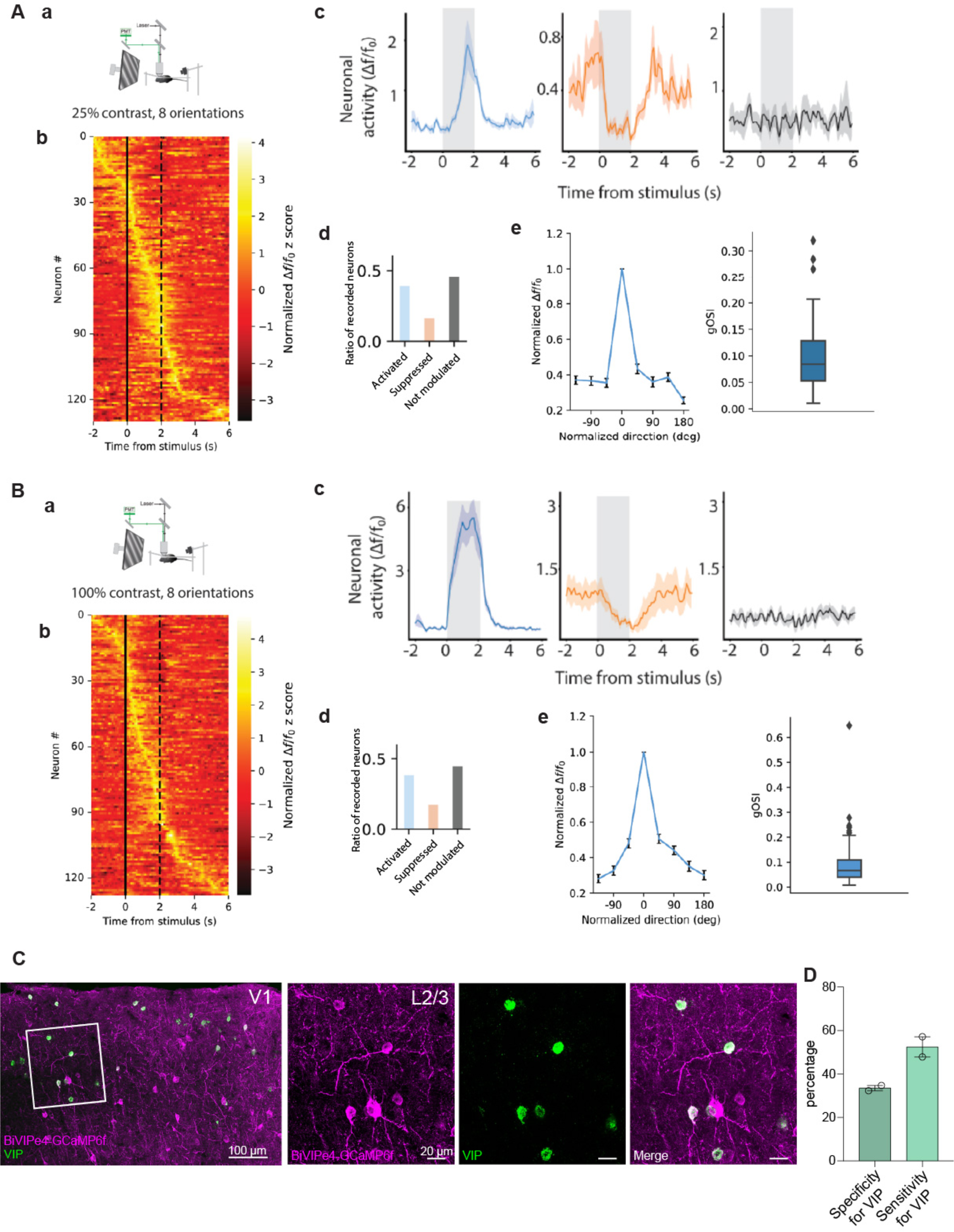
Imaging visual responses of VIP interneurons targeted by BiVIPe4 in the visual cortex, related to Figure 4. (A) Response of BiVIPe4-GCaMP6f cells to low contrast in the visual cortex. (a) Mice were presented with drifting sinusoidal gratings (25%; 8 directions). (b) Normalized activity (z-score) of all imaged VIP interneurons in response to the visual stimuli (*n* = 130 neurons from N=3 mice). Solid line: start of visual stimuli, dashed line: end of visual stimuli. (c) Examples of BiVIPe4-GCaMP6f cells activated (*left*), suppressed (*middle*), and not modulated (*right*) by the visual stimuli in their preferred direction. (d) Ratios of the different responses of VIP interneurons relative to total population (Activated: 50/130 (38.5%), suppressed: 21/130 (16.2%), not modulated: 59/130 (45.4%); paired *t*-test). (e) *Left panel*: average responses of BiVIPe4-GCaMP6f cells activated by low contrast stimuli in all eight directions (0 is the preferred direction of each neuron). *Right panel*: Global orientation selectivity index (gOSI) of BiVIPe4-GCaMP6f cells activated by the visual stimuli. (B) Response of BiVIPe4-GCaMP6f cells to high contrast in the visual cortex. (a) Mice were presented with drifting sinusoidal gratings (100%; 8 directions). (b) Normalized activity (z-score) of all imaged BiVIPe4-GCaMP6f cells in response to the visual stimuli (*n* = 128 neurons from N=3 mice). Solid line: start of visual stimuli, dashed line: end of visual stimuli. (c) Examples of BiVIPe4-GCaMP6f cells activated (*left*), suppressed (*middle*), and not modulated (*right*) by the visual stimuli in their preferred direction. (d) Ratios of the different responses of VIP interneurons relative to total population (Activated: 49/128 (38.5%), suppressed: 22/128 (17.2%), not modulated: 57/128 (44.5%); paired *t*-test). (e) *Left panel*: average responses of BiVIPe4-GCaMP6f cells activated by high contrast stimuli in all eight directions (0 is the preferred direction of each neuron). *Right panel*: Global orientation selectivity index (gOSI) of BiVIPe4-GCaMP6f cells activated by the visual stimuli. (C) *Left panel*: Representative image of BiVIPe4-GCaMP6f expression in V1 L2/3 (*in magenta*) with fluorescent *in situ* hybridization (FISH) for the *VIP* marker gene (*in green*). The GCaMP6f -positive cells co-expressing the marker gene (merge) are labeled *in white. Right panels*: zoomed-in image of the ROI from the left panel. Scale bars are shown in the figure. (D) Bar graph showing the specificity and sensitivity of BiVIPe4-GCaMP6f for the *VIP* marker. N=2 mice. Error bar represents SEM.

**Figure S8.**
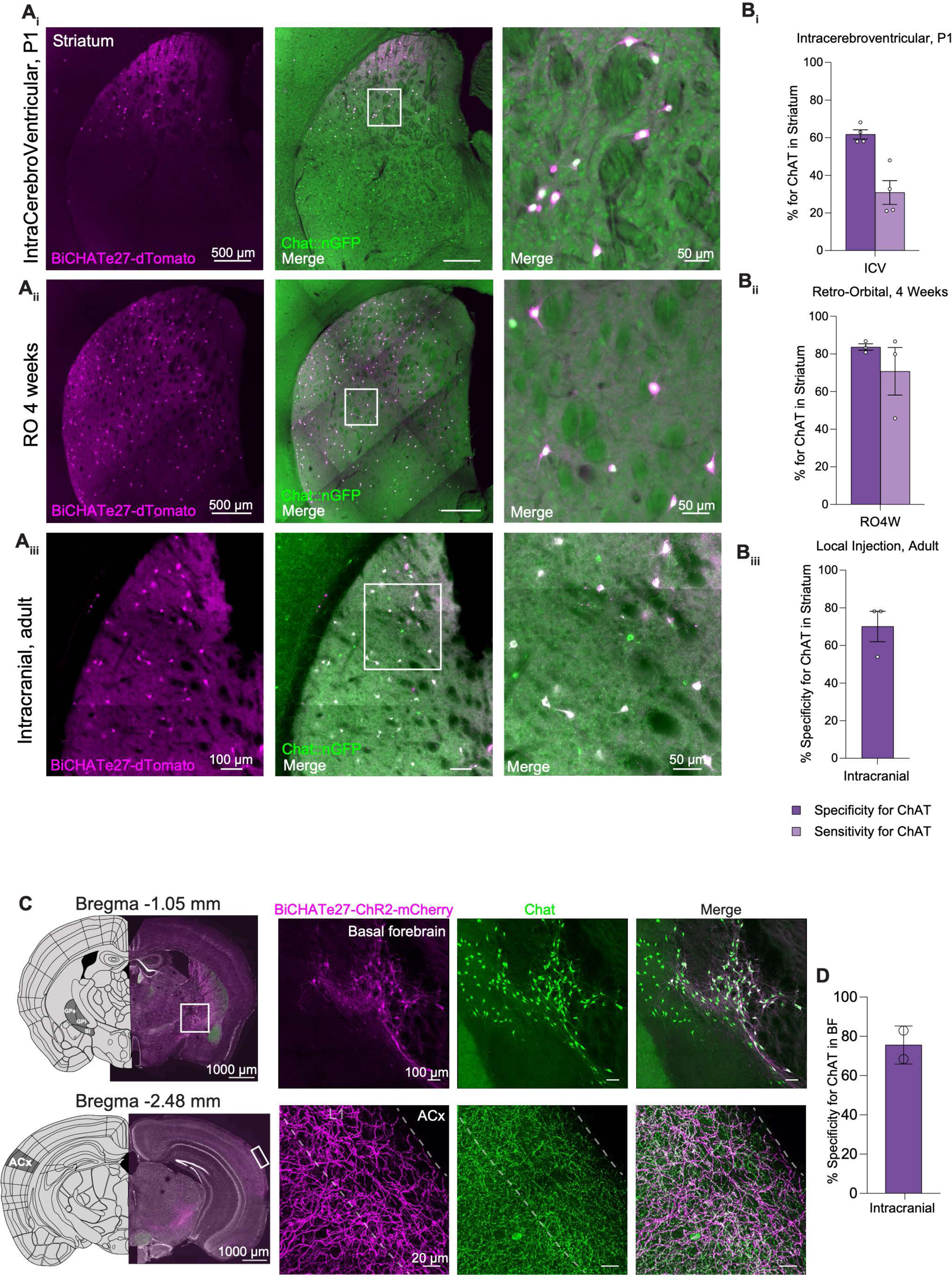
BiCHATe27 in Mouse, related to Figure 5. (A) Representative image showing the expression of dTomato reporter *(in magenta*) under the control of the BiCHATe27 enhancer, and the genetically-expressed GFP reporter under the control of Chat-cre (F) (*in green*) in striatum of the mouse brain, following ICV injection at P1 (*i*) retro-orbital AAV injection at 4 weeks (*ii*), and intracranial, stereotactic injections (*iii)*. dTomato-positive cells co-expressing CHAT (merge) are labeled *in white*. *Right panel*: zoomed-in image from the left panel. Scale bars are indicated in the figure. (B) Bar graphs showing the percentage specificity and percentage sensitivity of the BiCHATe27 enhancer in targeting CHAT-positive neurons in the striatum across the injection conditions indicated. N=3-4 mice. Error bars represent SEM. (C) *Left pane*l: Intracranial stereotactic injection location (basal forebrain - *top panel*) and recording location (L1 of primary auditory cortex (A1) - *bottom panel*) for the experiment in Figure 5H and 5I. *Right panel*: Representative image showing the expression of mCherry-expressing cells *in magenta (top panel*) and axons (*bottom panel*) under the control of the BiCHATe27 enhancer, and the immunolabeling of the marker CHAT (*in green*), following intracranial, stereotactic injections in the basal forebrain. mCherry-positive cells and axons co-expressing CHAT (merge) are labeled *in white*. (D) Bar graphs showing the percentage specificity of the AAV-BiCHATe27-ChR-mCherry in targeting CHAT-positive neurons in the cholinergic basal forebrain. N=2 mice. Error bar represents SEM.

**Figure S9.**
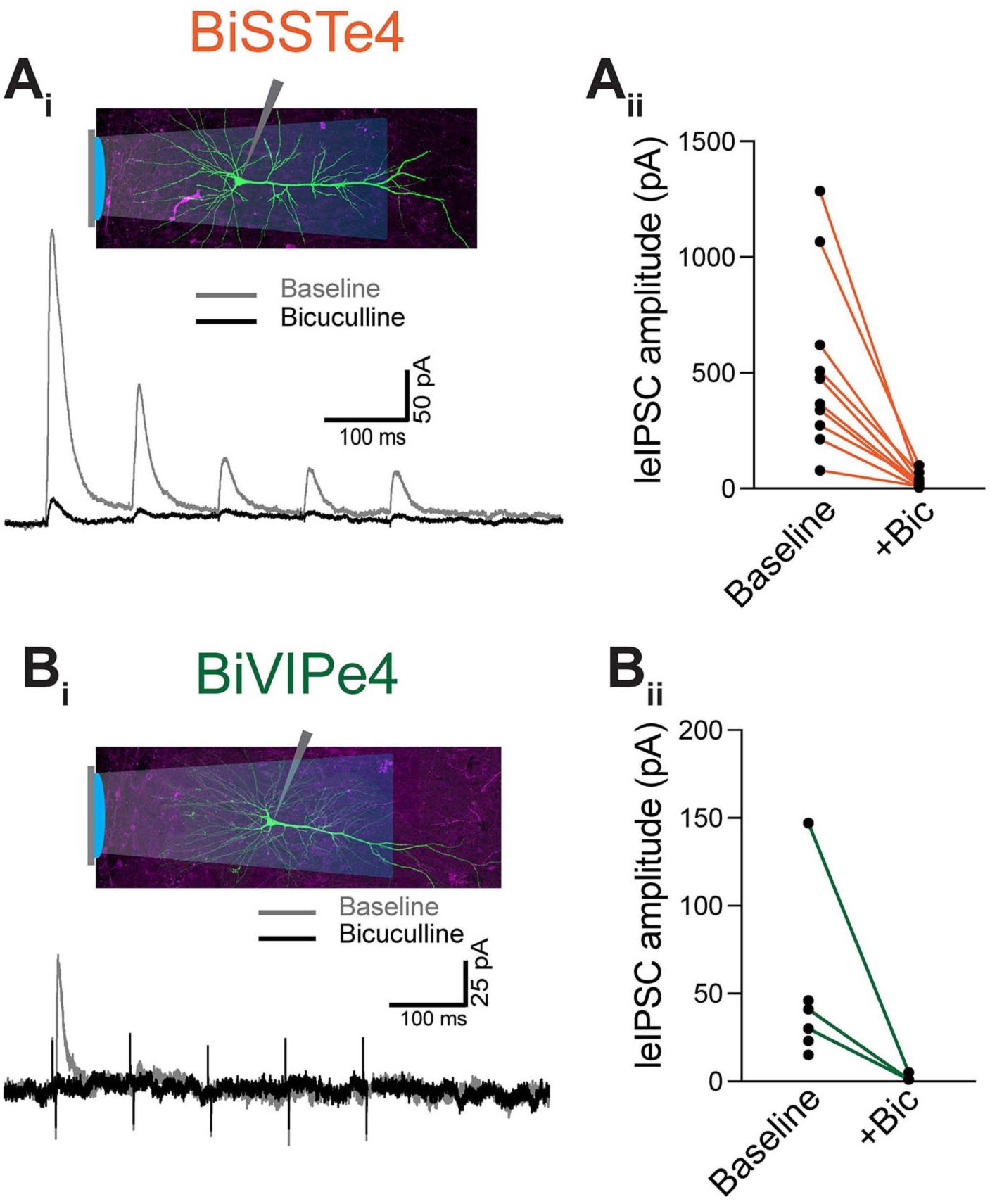
Optogenetic characterization of BiSSTe4 and BiVIPe4 in rhesus macaque, related to Figure 6. Functional validation of (A) BiSSTe4, and (B) BiVIPe4 in rhesus macaque. (A-Bi) Example traces of light-evoked IPSCs recorded from pyramidal cells following ChR2 activation of rAAV transduced cells at baseline and in the presence of GABAA antagonist bicuculline. (*Inset*) biocytin filled cell-reconstructions of recorded pyramidal cells. (A-Bii) Quantification of light-evoked IPSC amplitude at baseline and in the presence of bicuculline.

**Figure S10.**
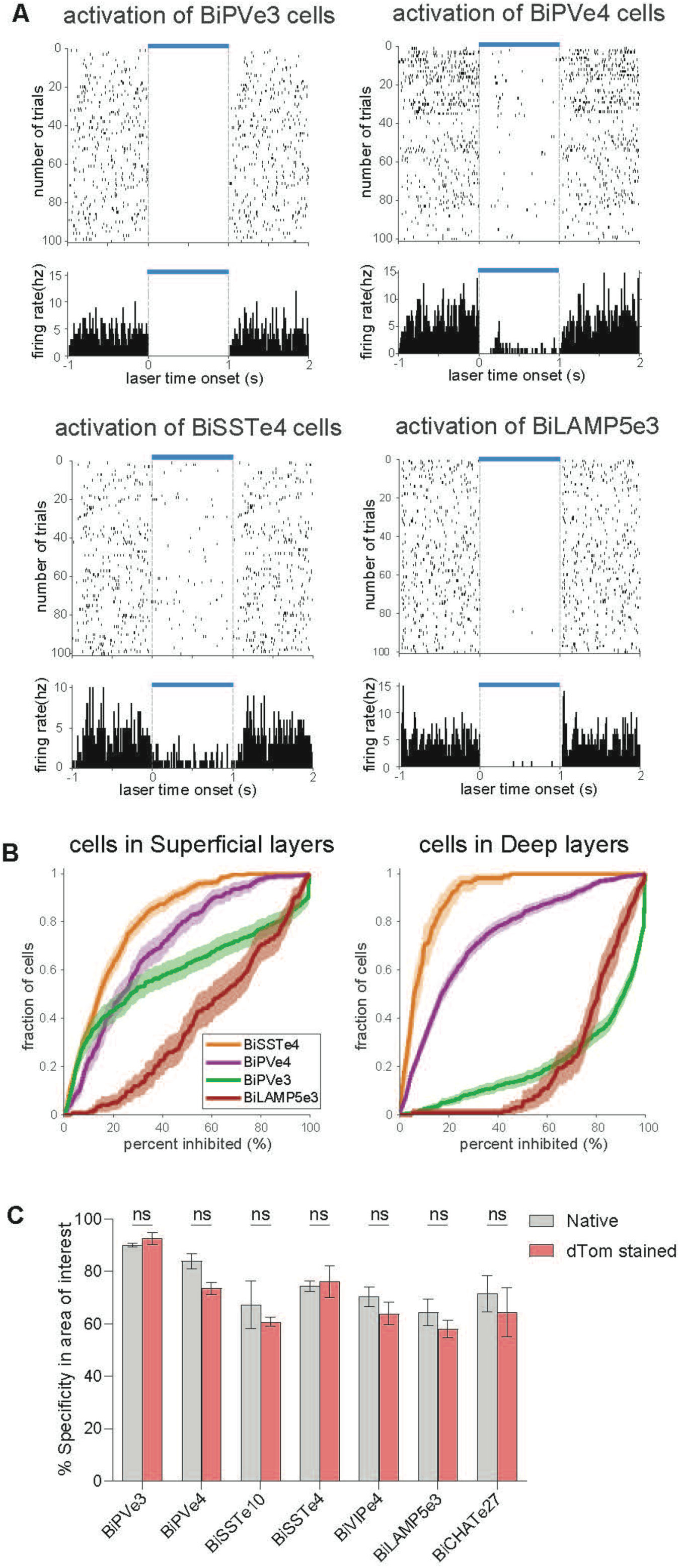
*In vivo* optogenetic inhibition using AAV-enhancer tools and cell count with native and dTom stained conditions, related to Figures 2-5. (A) Optogenetic activation of BiPVe3-, BiPVe4-, BiSSTe4- and BiLAMP5e3-ChR2-mCherry-positive cells provides strong inhibition of cortical neurons. (B) Plot showing the percentage inhibition resulting after optogenetic activation of each enhancer indicated, in superficial (*left panel*) and deep (*right panel*) cortical layers in V1. N=6 mice. (C) Bar graph showing the specificity of each enhancer indicated for the expected cell type-specific marker in the brain area or interest (S1 for all except for BiCHATe27, calculated in the striatum), both with native dTomato fluorescence (*grey*) and with dTomato immunostaining (*red*). Multiple unpaired t test was done. ns=not significant.

## STAR METHODS

### KEY RESOURCES TABLE

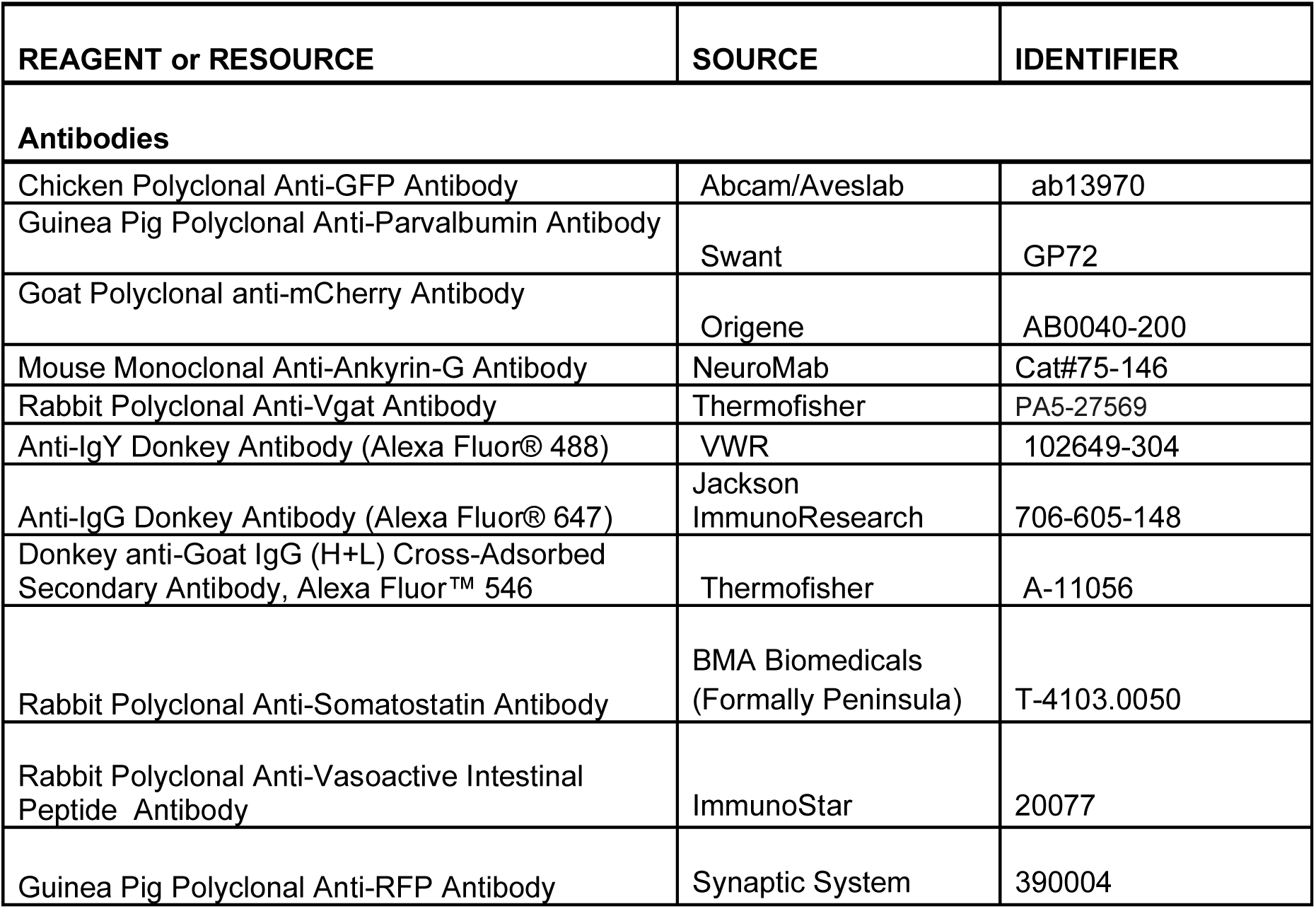
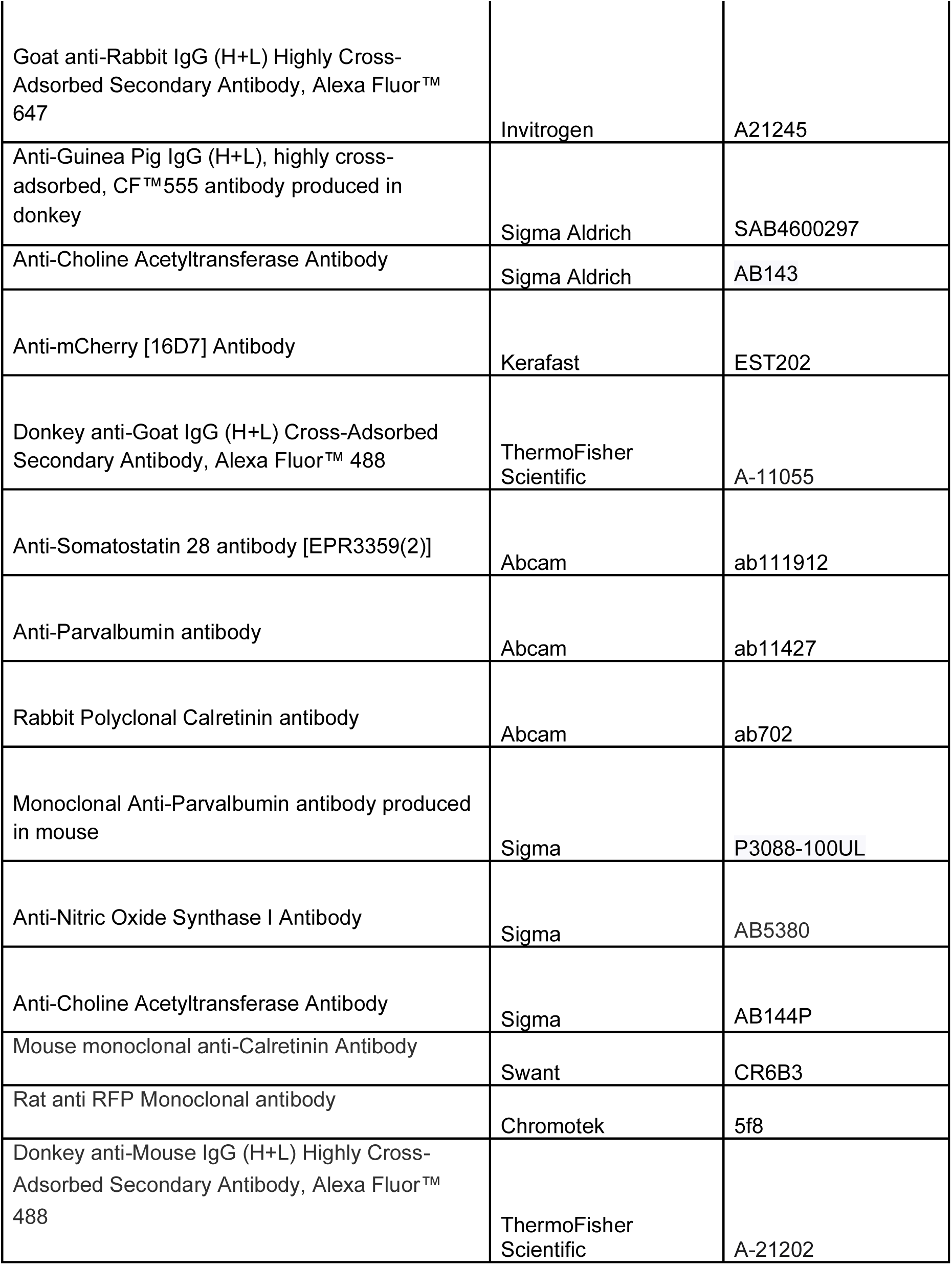
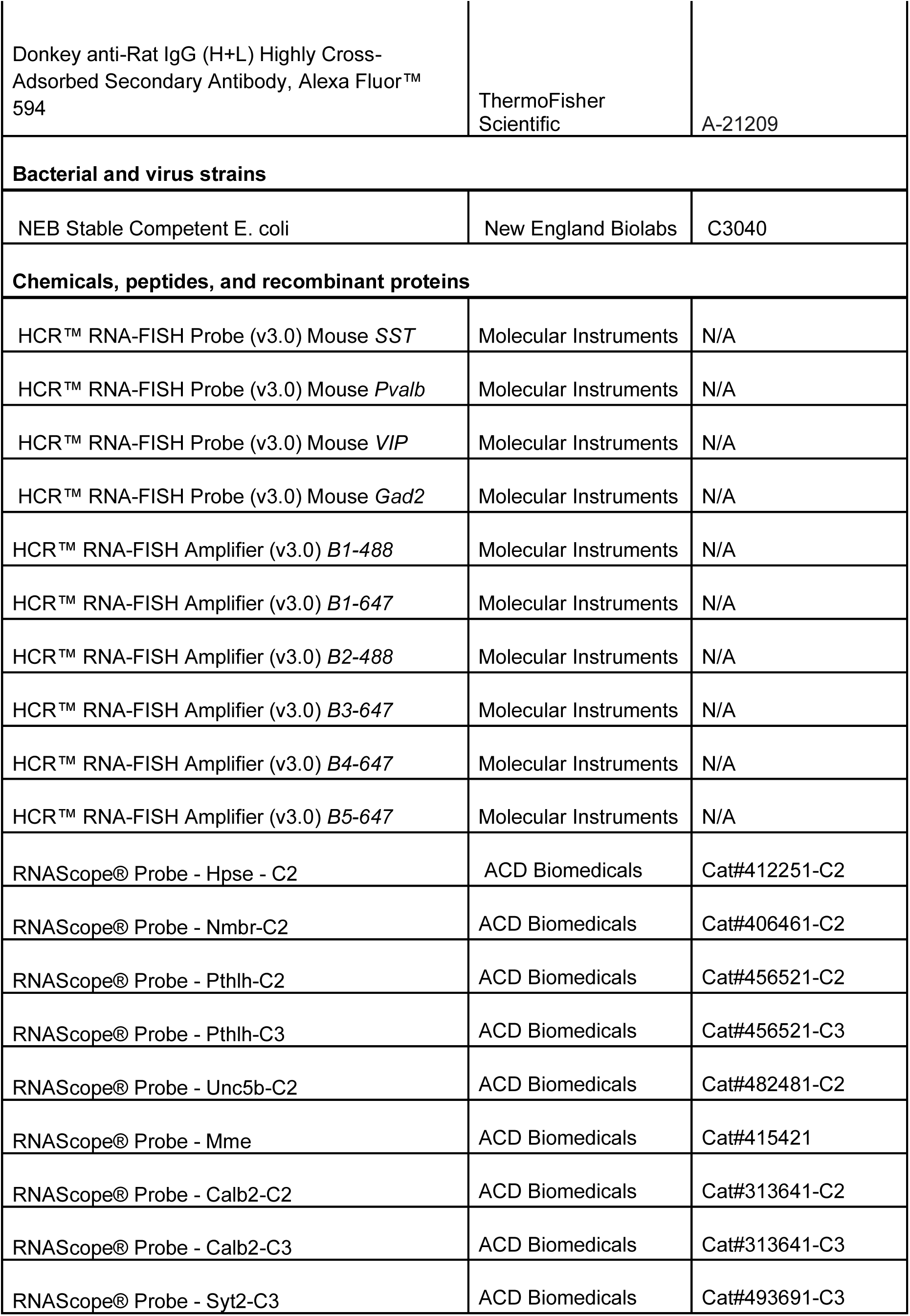
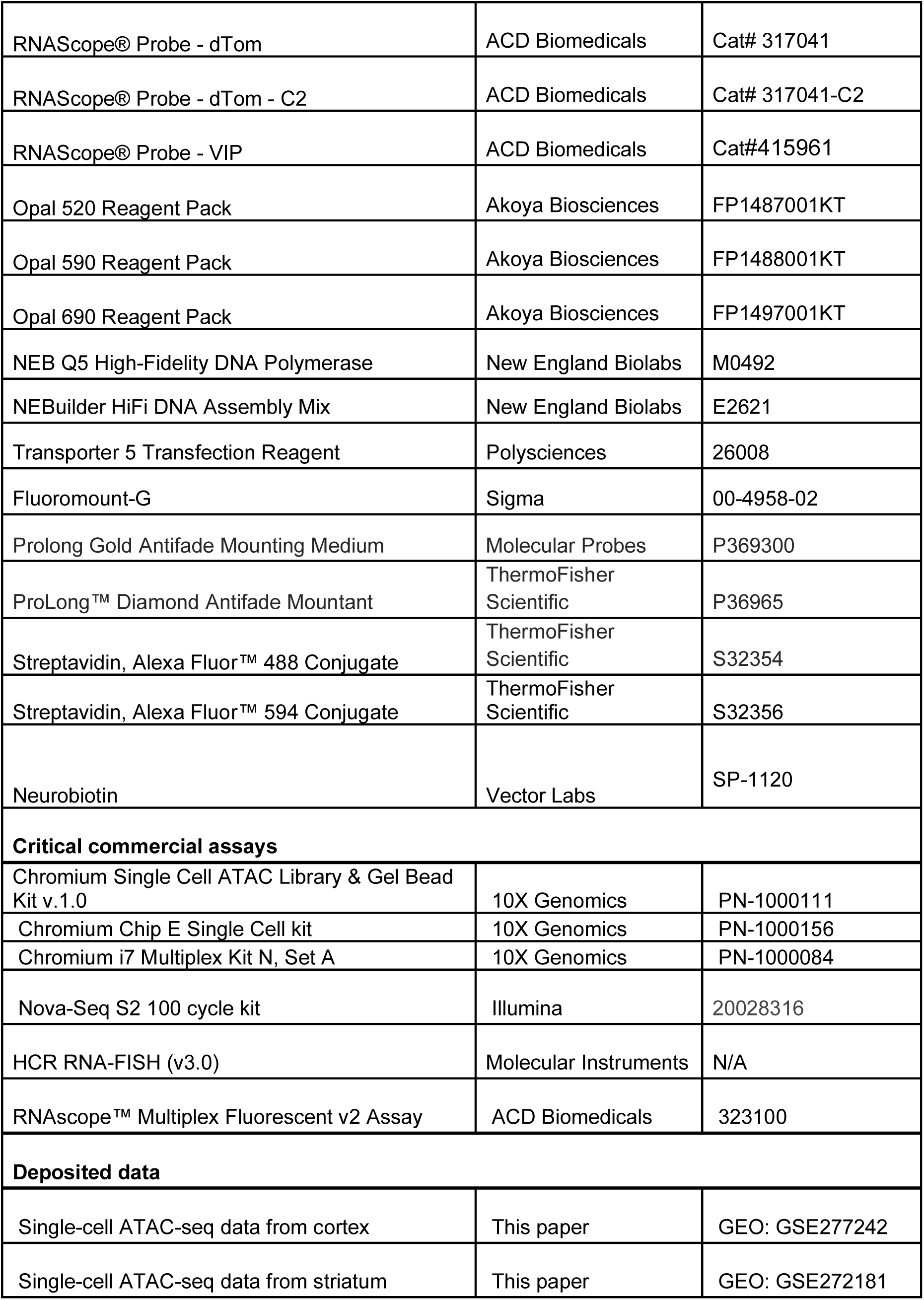
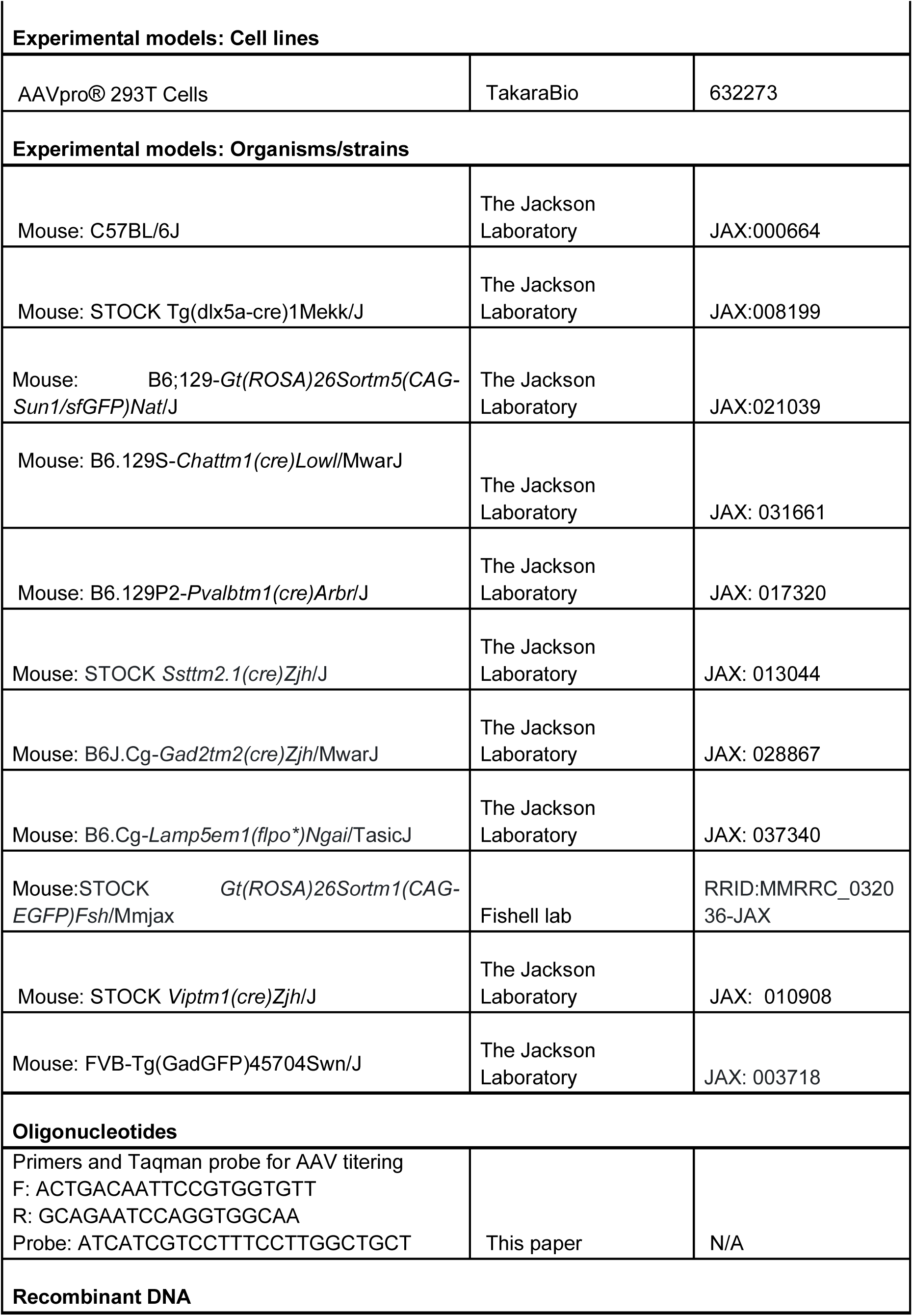
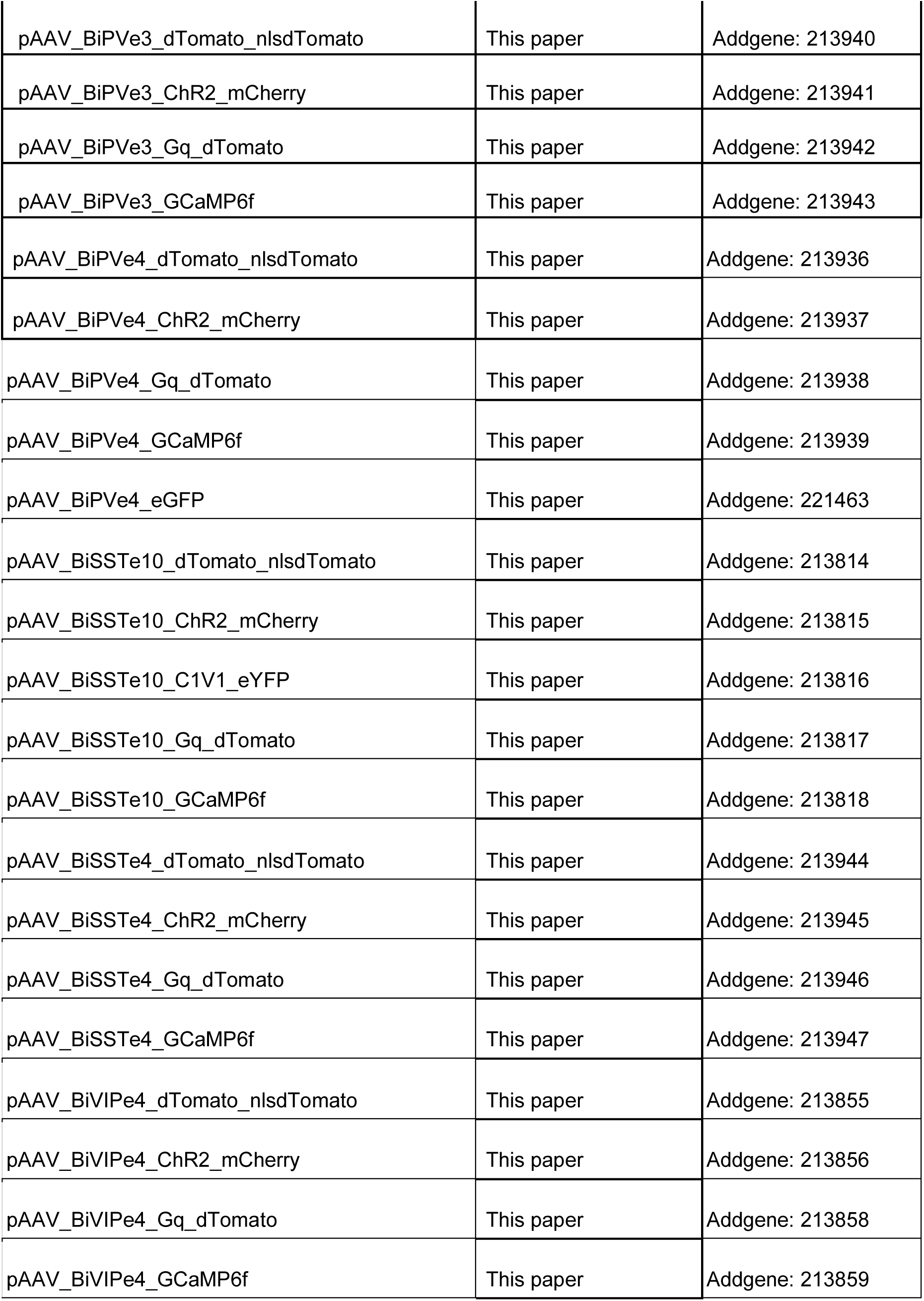
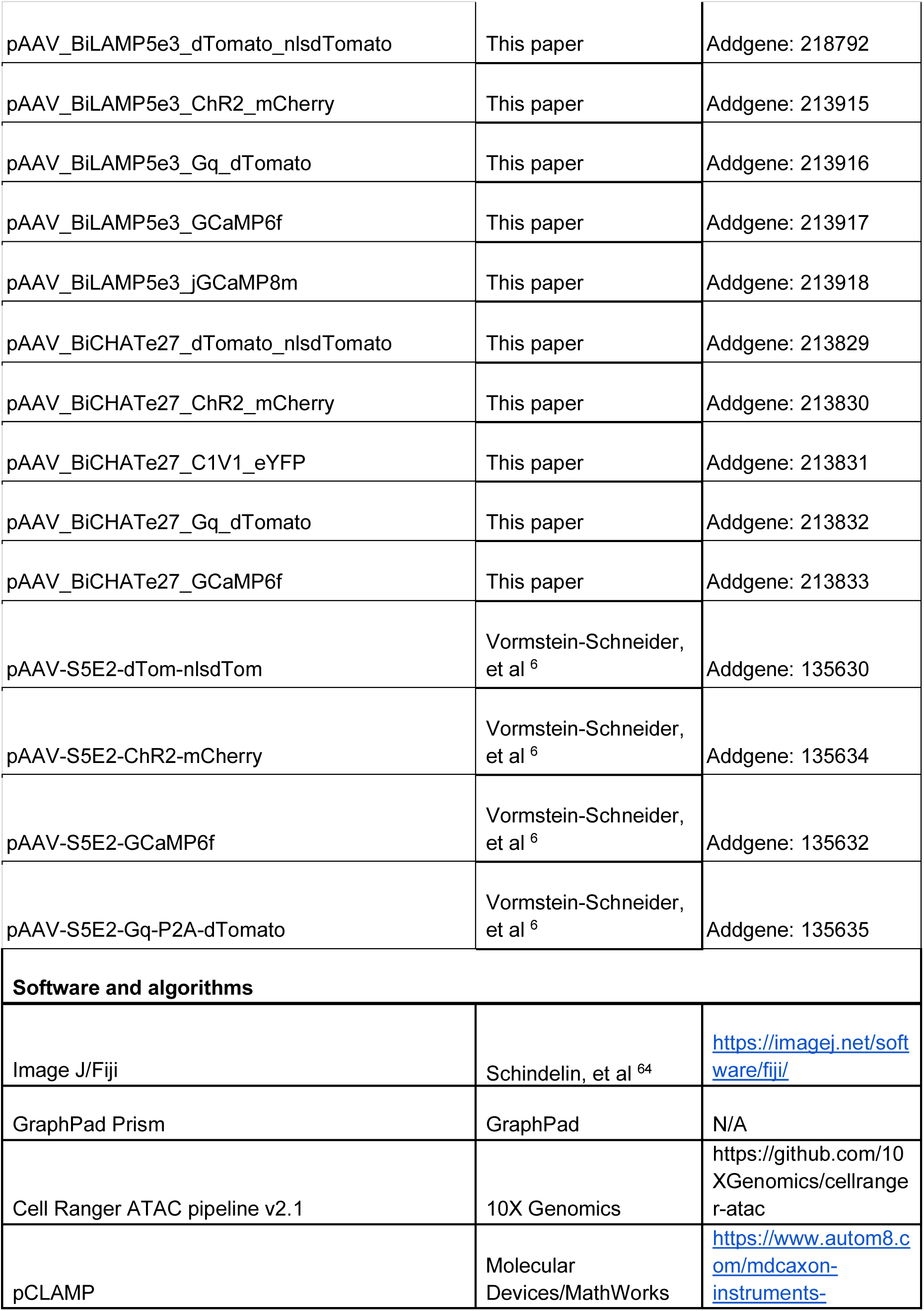
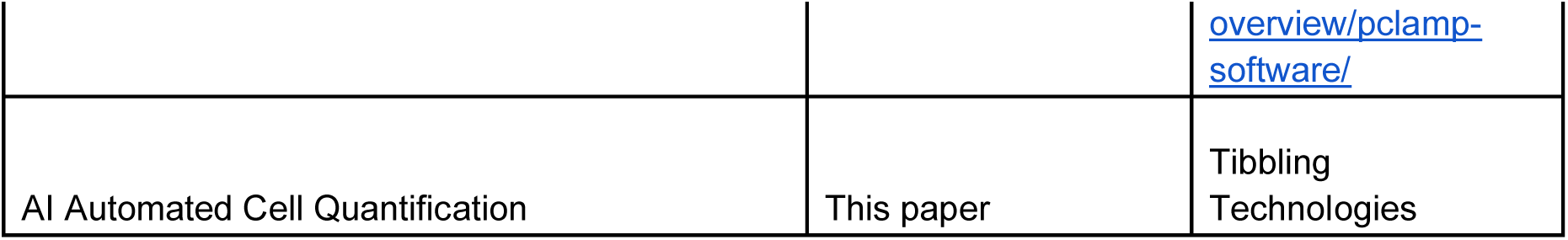

### EXPERIMENTAL MODEL AND SUBJECT DETAILS

#### Mouse breeding and husbandry

All procedures were carried out in accordance with Institutional Animal Care and Use Committee protocols 0156-03-17-2 at the Broad Institute of MIT and Harvard.

See “key resources table” for a detailed list of genetic lines used.

Animals were provided food and water *ad libitum* and were maintained on a regular 12-h day/night cycle at no more than five adult animals per cage. Macroenvironmental temperature and humidity were kept within the ranges of 17.8–26.1C and 30–70%, respectively. These parameters were monitored closely and controlled within rodent colony rooms.

### METHOD DETAILS

#### Single cell ATAC-seq library construction and data processing

The scATAC-seq data for cortex interneuron was collected as described in our previous study.^56^ Data was collected from three brains of postnatal day 28 Dlx5a-cre::INTACT mice (GSE165031 and GSE277242). For striatum scATAC-seq, two brains from postnatal day 30 Dlx5a-cre::INTACT mice were harvested, sectioned coronally on a mouse brain slicer (Zivic Instruments), and the basal ganglia region was dissected in ice-cold artificial cerebrospinal fluid (ACSF). Tissue was then transferred to a Dounce homogenizer containing lysis buffer (10 mM Tris-HCl, 10 mM NaCl, 3 mM MgCl2, 0.01% Tween-20, and 0.01% IGEPAL CA-630, 0.001% digitonin). Tissue was homogenized with 10 strokes of pestle A and 10 strokes of pestle B, and incubated for 5 min on ice before being filtered through a 30-μm filter and centrifuged at 500g for 10 min at 4°C. The pellet was resuspended in 1% bovine serum albumin for sorting GFP+ nuclei on a Sony SH800S cell sorter. Nuclei were sorted into diluted nuclei buffer (10x Genomics). A total of 15,168 GFP+ nuclei were used for scATAC-seq library construction. The scATAC-seq library was prepared using the 10x Genomics platform with the Chromium Single Cell ATAC Library & Gel Bead Kit v.1.0 (PN-1000111), Chromium Chip E Single Cell kit (PN-1000156) and Chromium i7 Multiplex Kit N, Set A (PN-1000084) as per manufacturer’s instructions. Libraries were sequenced using a Nova-Seq S2 100 cycle kit (Illumina).

Reads from scATAC-seq libraries (FASTQ files) were aligned to the mouse mm10 (GENCODE vM23/Ensembl98) reference genome provided by 10x Genomics and the fragment file was produced with the Cell Ranger ATAC pipeline v2.1.0 (10x Genomics) with default parameters (cellranger-atac count). The initial set of peaks were then identified from the fragment files output by Cell Ranger ATAC using MACS2 v2.2.7.1 with the following parameters: ‘--nomodel --shift -100 --extsize 200 -f BED -g 1.87e9’. For the cortex interneuron scATAC-seq data, the peak x cell matrix was generated from the peak set and the fragment file by the R package Signac^60^ v1.7.0 in R v4.1.3, and Signac was used for the data preprocessing. Cells were retained based on the following thresholds: peak_region_fragments <100,000, peak_region_fragments >3,000, >30% of reads in peaks, nucleosome_signal <4, and TSS enrichment score >2. The TF-IDF algorithm was used for the normalization, the LSI algorithm was used for the dimensionality reduction, and the UMAP algorithm was used to generate the 2D-visualization. Cells were then annotated based on the inferred activities of known marker genes. For the striatum scATAC-seq data, processing and analysis were conducted using a novel single-cell ATAC-seq analysis toolkit, which will be described in detail in a forthcoming publication (Dai et al., in preparation).

### Nomination of cell type-specific enhancers

We initially selected enhancer candidates using methods modified from what we have previously described.^6^ We selected and tested 40 enhancer candidates using this method and two of them (BiSSTe10 and BiCHATe27) were effective. We subsequently improved the selection methods computationally and the rest of the enhancers were selected using this improved method. The details of this computational pipeline will be described in a separate publication (Dai et al., in preparation). Briefly, the COSG package v0.9.0 was used with default parameters except for setting mu to 100. With the ranked top enhancer candidates lists output by COSG, we manually checked the peak distributions in the bigwig files corresponding to each cell type and selected the genomic regions surrounding the peak summits. Additionally, the sequence conservation among NHPs (calculated via phastCons with mm10.60way.phastCons.bw, downloaded from http://hgdownload.cse.ucsc.edu/goldenpath/mm10/phastCons60way/)^57,58^ was taken into consideration for the prioritization of enhancer candidates. Using this method, we selected and tested 29 enhancer candidates and 5 of them (BiPVe3, BiPVe4, BiSSTe4, BiVIPe4, and BiLAMP5e3) were the most effective. Of note, the enhancer BiVIPe4 was initially hand-picked in the intron of a VIP-specific gene, *Grpr*, before the new computational pipeline was developed. We subsequently verified that this enhancer can also be identified with the new COSG-based method.

### rAAV plasmids cloning

All enhancer sequences were either synthesized as gene fragments (GenScript) or amplified from mouse (C56BL6/J) genomic DNA using NEB Q5 High-Fidelity DNA Polymerase (NEB M0492). The enhancer fragments were inserted into backbone plasmids with different payloads using NEBuilder HiFi DNA Assembly Mix (NEB E2621). Specifically, the following plasmids were used as backbone for cloning: pAAV-S5E2-dTom-nlsdTom (Addgene plasmid #135630), pAAV-S5E2-ChR2-mCherry (Addgene plasmid #135634), pAAV-S5E2-GCaMP6f (Addgene plasmid #135632) and pAAV-S5E2-Gq-P2A-dTomato (Addgene plasmid #135635). All plasmids were grown in NEB Stable Competent *E. coli* (NEB C3040) at 37°C and were verified by whole plasmid sequencing (Primordium Labs). Plasmids related to this study and their sequences are available from Addgene (see “key resources table” for detailed information). AAV viral preps associated with some of the plasmids are available from Addgene (https://www.addgene.org/Gordon_Fishell/) and UCI Center for Neural Circuit Mapping (https://cncm.som.uci.edu/enhancer-aavs/).

Enhancer sequence information is as follows:

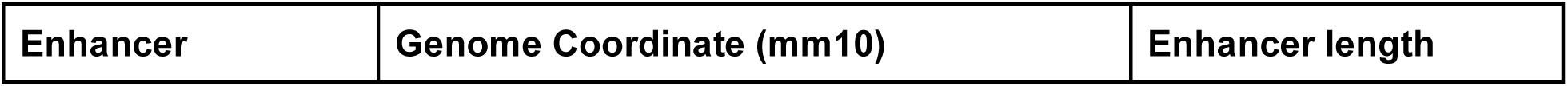
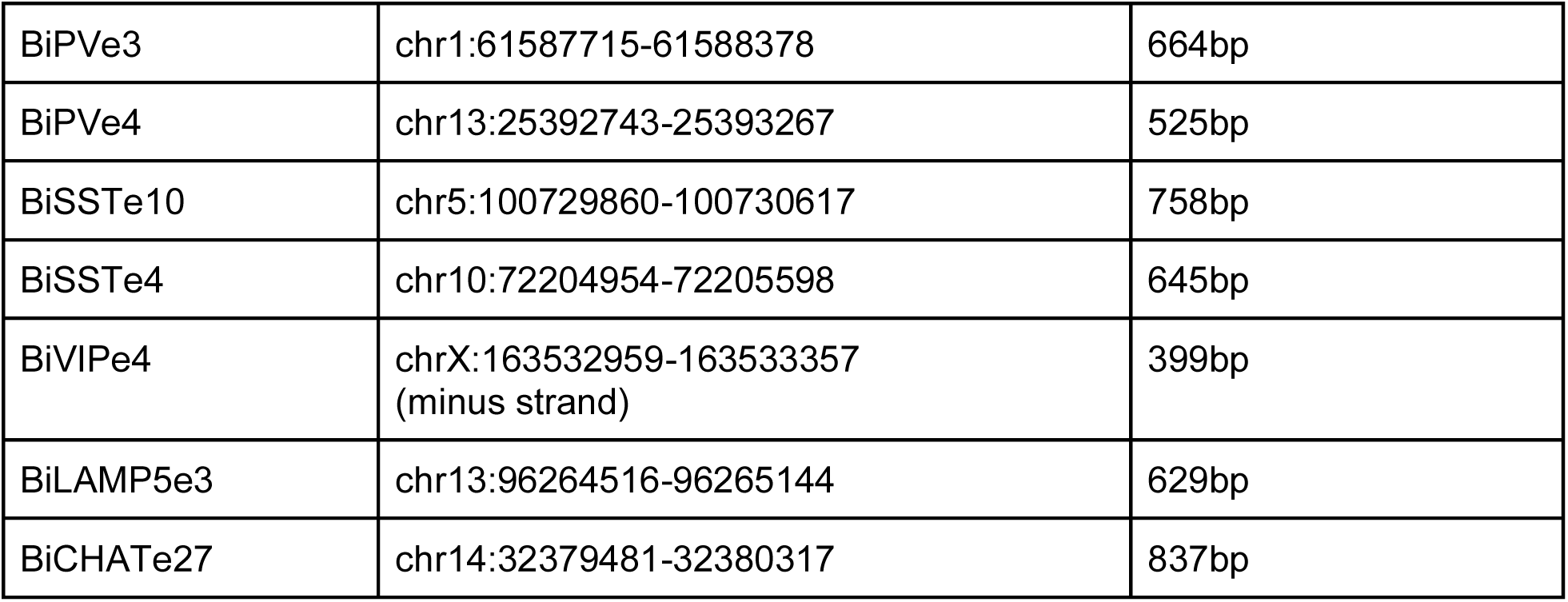

#### rAAV production and titering

AAVs with PHP.eB capsid were produced in AAVpro 293T cells (Takara 630073) following previously published protocol.^59^ Triple transfection was performed with Transporter 5 Transfection Reagent (Polysciences 26008). AAVs were harvested 72 hours post-transfection and purified on iodixanol gradient. AAV titers were determined by ddPCR assay (BioRad AutoDG Droplet Digital PCR System) using primers and probe targeting the WPRE sequence, which is common to all constructs (see “key resources table” for the sequences). A control virus with known titer was included in each titering assay to ensure consistency.

#### Intracranial, systemic (retro-orbital) and intracerebroventricular injections in mice

##### Mouse systemic

Procedure was performed using an all-in-one anesthesia induction chamber and nose cone with 1-3% isoflurane in oxygen flowing at 0.5 L/min. Anesthesia depth was validated with bilateral toe pinch prior to starting the procedure. Retro-orbital injections were performed in 4 weeks-old (P28-P35) or 8 weeks-old animals (P50-P65), both in males and females. 2.0E+11 viral genomes were injected using a BD Biosciences 0.3 mL insulin syringe (ref: 32470) in the retro-orbital sinus, for a final injected volume of 80uL per animal.

##### Mouse intracranial

Meloxicam was injected subcutaneously before the surgery at 5 mg/kg of body weight, and Buprenorphine SR was also injected as a pre-operative analgesic. Anesthesia was induced using an all-in-one anesthesia induction chamber with 1-3% isoflurane in oxygen flowing at 0.5 L/min. Once full depth of anesthesia was reached, the mouse was moved onto a Stoelting Digital Mouse Animal StereoTaxic Frame (ref: 106739) to perform the surgery, with a nose-cone for continuous anesthesia maintenance with 1-3% isoflurane in oxygen flowing at 0.5 L/min. Stereotactically-guided intracranial injections in adult mice (P60-70, both males and females) were performed using a Nanoinject III at 1nL/sec, mainly in the primary somatosensory cortex (coordinates relative to Bregma: -1.00 mm antero-posterior, +/- 2.5 mm medio-lateral, -0.25 and -0.4 dorso-ventral). Virus was diluted using PBS1x to a final titer of 2-4.00E+12 vp/ml and a final volume of 100/200 nL was injected in each location. Post-operative monitoring was performed for 5 days post-injection, and mice were given meloxicam subcutaneously for 2 days post-procedure at 5 mg/kg of body weight. For injection in P5 pups, we used cryoanesthesia and we injected 150 nL of virus at 2 nL/second bilaterally in S1 (AP: +1.5, ML: +/- 1.4 and DV: -0.3 from Lambda).

##### Neonatal mouse intracerebroventricular

Mouse pups (P0-1) were anesthetized on ice for 2-5 minutes avoiding direct contact with ice to avoid freeze damage to the skin of the pup. Virus was diluted using PBS1x to 1E+10 total viral particles in 2-3 µl of final volume. Intracerebroventricular injections (either unilateral or bilateral) were performed using a Nanoinject III at 25 nL/sec in the ventricle (coordinates relative to lambda: + 1.5 mm antero-posterior, +/- 0.8 mm medio-lateral, and -1.5 mm dorso-ventral). Post-surgery, pups recovered on a heating pad for 10 minutes, monitored for color respiration, movement and posture. Once all the pups were injected and regained homeostasis, they are returned to the dam together to encourage re-acceptance by the dam.

#### Immunohistochemistry

All antibodies used in this study are commercially available and have been validated by the manufacturer (see key resources table for detailed list). Injected mice were trans-cardially perfused with 4% paraformaldehyde (PFA) 3 weeks post-injection. The brains were placed in 4% PFA for 2 hours or overnight, washed 3 times with PBS1X, incubated at 4C with 30% sucrose for two days and then sectioned at 40 um using a LEICA VT1000S Microtome. Floating sections were permeabilized with 0.3% Triton X-100, 5% normal donkey serum, and 1xPBS for 30 minutes. The sections were then incubated overnight in 0.1% Triton X-100 and 5% donkey serum with the combination of primary antibodies at 4C, with gentle rocking. The sections were then washed four times with 1xPBS, incubated with fluorophore-conjugated secondary antibodies at 1:1,000 dilution (Invitrogen, USA), counterstained with DAPI (Sigma, USA) and mounted on glass slides using Fluoromount-G (Sigma, USA). Sections not stained were stored in antifreeze buffer (30% glycerol, 30% ethylene glycol, 2% sodium azide in 1xPBS) and kept for long term storage at -20C.

Whole slice images were acquired using a Zeiss Axioscan 7 Slide scanner, with a 20x objective. Tiled images and snapshots were acquired using a Zeiss Axio Imager 2 using a 20x objective. High resolution images of BiPVe3 and BiPVe4-positive synaptic terminals were acquired using a Zeiss LSM 900 Confocal Microscope with a 63x objective.

#### Image analysis

For the analysis of virally-labeled dTom-positive cells across brain areas, sections from five different medio-lateral coordinates based on the Allen Brain Atlas were collected (sections #1,5,10,15 and 20) and dTom-positive cells quantified in all brain areas reported in the reference atlas. The sections with the highest number of dTom-positive cells in a given brain region were used (as it would be the most representative of that region) and data for selected region were used (as it would be the most representative of that region) and data for selected brain areas were reported in the bar-graphs. For example, coordinate 5 was used to report dTom-positive cells in the hippocampus because it is the only coordinate where the hippocampus is best represented and visible.

Sagittal sections, as well as whole slice coronal sections, were all quantified through Tibbling Technologies. A small set of images was quantified manually using Fiji image processing package (Image J), using the Cell Counter plugin.

For automated image analysis (by Tibbling Technologies), we deployed an advanced customized AI-based framework, which includes a high-performing deep neural network (DNN) for neuron classification and segmentation. This DNN was initially trained on a large corpus of neuron samples and subsequently fine-tuned on the shortlisted data collected for this study. The AI framework is optimized based on multiple loss functions including the Structural Similarity Index Measure (SSIM), which is a combination of three different kinds of measurements: brightness (b), contrast (c), and structural (st) comparison. The SSIM between two images *x* and *y* can be expressed mathematically as:

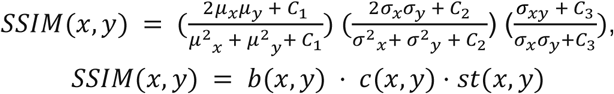

where brightness (b) is computed through the mean (*μ*) of the intensity of a given image, contrast comparison (c) uses the standard deviation (*σ*) and the structural comparison (st) is computed based on the covariance of *x* and *y*. In SSIM, the brightness, contrast, and structure components are relatively independent, which is particularly effective for identifying neurons based on their structures and intensity. Intensity plays a crucial role in segmentation as it allows for the differentiation of neurons from the background and other noisy structures, ensuring accurate and precise identification of cells. The complete loss function is as follows:

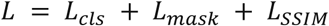

where *L*_*cls*_, *L*_*mask*_ and *L*_*SSIM*_ are the pixel classification, mask prediction and structural similarity losses, respectively. *L*_*cls*_ is calculated as log loss for two class labels (neuron vs. background).

*L*_*mask*_ is computed as average cross-entropy loss for per-pixel binary classification. *L*_*SSIM*_ is the structural similarity between the predicted and ground truth images, since higher SSIM means large similarity, (1-SSIM) is employed for the loss, *L*_*SSIM*_. It ensures that the overall structural integrity of the segmented image is preserved.

During inference, we ensure the reliability of our predictions by incorporating a confidence score derived from the model’s final output layer. For a given input image (*I*), the confidence score is obtained from the softmax output (*p*^) of the classification head for the predicted segmentation mask (*M*^):

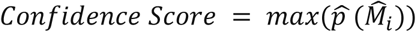

We set a minimum confidence threshold (*T*_*c*_) of 0.6. If the confidence score meets or exceeds *T*_*c*_, the prediction (*P*) is considered reliable.

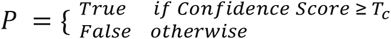

Furthermore, we deployed an automated image registration mapper to the Allen Mouse Brain atlas, which precisely aligned each sample with the corresponding atlas template section. This process is entirely automated, utilizing advanced image registration techniques to ensure accurate and consistent mapping to the brain atlas. The automated pipeline includes corresponding atlas section matching through feature-based alignment, AI-based spatial transformation of the atlas sample, and simultaneous warping of atlas labels to precisely map the full brain atlas. By integrating multi-scale feature extraction and optimization algorithms, our deployed method achieves high accuracy and robustness in aligning diverse brain regions. The neural density in a given brain region is quantified by dividing the number of cells by the area of the brain region.

We validate our end-to-end image analysis pipeline both qualitatively and quantitatively. Qualitative validation involves visually inspecting the registered images for anatomical accuracy. For quantitative validation, we perform a rigorous comparison of the automated neuron segmentation outputs, including cell counts, against manually obtained ground-truth data of enhancer images from the diverse regions of the brain sections. This comparison ensures comprehensive validation across different anatomical areas. Our approach achieves optimal correspondence in both cell counts and DICE score metric with the ground-truth, demonstrating the high precision and reliability of our approach.

Images quantified using FIJI were normalized according to the area of the ROI (region of interest). Area-normalized (either AI-quantified or manually quantified) cell counts for both the reporter-and marker-positive cells were used to calculate the percentage specificity and sensitivity of enhancers using the following formulas:

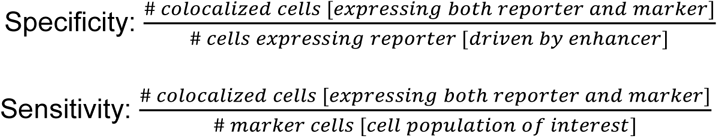

where specificity indicates the selectivity of the enhancer in targeting a given cell type and sensitivity indicates the efficiency of the enhancer in marking all cells of the population of interest in a given area and under the conditions used.

#### *In situ* hybridization and quantification

##### HCR-FISH

Mice brains were harvested after perfusion with 4% PFA in PBS1X and post-fixed in 4% PFA for 2 hours on ice or at 4°C. Brains were then washed with 1XPBS three times and cut in cold PBS1X using a vibratome (Leica VT1200S), in 40 µm thick sections. Brain sections were mounted onto positively charged microscope slides (Superfrost Plus^TM^ Microscope Slides White Tab, # 1255015) and dried completely before placing them at -80°C for long-term storage before performing FISH.

HCR RNA-fluorescent *in situ* hybridization (HCR RNA-FISH) assay was carried out following the manufacturer’s instructions (Molecular Instruments^TM^, see key resources table for detailed list of probes and amplifiers used) with a few modifications. In particular, slides were incubated in 100% EtOH for 5 minutes only one time, the pre-hybridization step was increased to 30 minutes and proteinase K solution was not added to maintain the endogenous dTomato signal. Whole slice images were acquired using a Zeiss Axioscan 7 Slide scanner, with a 20x objective. Tiled images and snapshots were acquired using a Zeiss Axio Imager 2 using a 20x objective. We then performed automated image analysis as previously described.

##### RNAScope

Mouse brains were collected according to the ACDBio Multiplex Fluorescent v2 Kit protocol (ACDBio #323100) for fixed frozen tissue. 20um brain sections were cut using a Leica Cryostat 3050M. Tissue was pre-treated with a series of dehydration, H2O2 treatment, and protease III steps before incubation with the probe for 2 hours at 40 °C. Protease III incubation was performed at room temperature to better preserve protein. For the complete list of probes and reagents used, see key resources table. Three amplification steps were carried out prior to developing the signal with Opal Dyes (Akoya Biosciences). Samples were counterstained with DAPI and mounted using Prolong Gold Antifade Mounting Medium (Molecular Probes).

ROI images were acquired using a Zeiss LSM 800 Confocal Microscope with a 40x objective. Z-stacks (with 0.45 um steps in between) covering 2.3-2.4 um were 2D projected and quantification was manually performed with the Fiji image processing package (Image J), using the Cell Counter plugin on 2D projected images. Importantly, dTomato viral DNA signal can be found in nuclear aggregates and unspecifically picked up by the dTomato probe. For this reason, only cells showing fluorescent mRNA signals also outside the nucleus were counted as dTomato-positive. Specificity and sensitivity were then calculated as previously described.

#### Electrophysiological and morphological characterization of BiPVe3 enhancer

##### Electrophysiology recordings

Local stereotaxic injections were performed in mice 3 weeks of age in primary somatosensory (S1) cortex at the following coordinates: 1.0 mm posterior, 2.5 mm lateral, and 0.5 mm ventral relative to Bregma. 200 nl of 2e12 vg/mL titer virus (AAV.BiPVe3-dTom) was injected at a rate of 1 nl/min using a Nanoject III (Drummond Scientific).

4 weeks following viral injection, mice were anesthetized with isoflurane and the brain was transferred to ice-cold artificial cerebrospinal fluid (ACSF) containing (in mM) 87 NaCl, 75 sucrose; 2.5 KCl, 1.0 CaCl2, 2.0 MgSO4, 26 NaHCO3, 1.25 NaH2PO4, and 10 glucose, equilibrated with 95% O2 and 5% CO2, and sliced at a thickness of 300 μm using a Leica VT-1200S vibratome. Slices were allowed to recover in ACSF for 30 min at 32°C. Prior to recording, slices were transferred to a chamber under an Olympus BX61 microscope and continuously perfused at a rate of 3 ml/min and temperature of 32°C with recording solution containing (in mM): 125 NaCl, 2.5 KCl, 2.0 CaCl, 1.0 MgSO, 26 NaHCO, 1.25 NaH2PO, and 10 glucose. dTom+ cells in Layer 2/3 primary somatosensory cortex (S1) were identified under epifluorescence. For whole cell recordings of intrinsic properties, 3-5 MOhm borosilicate glass pipettes were filled with intracellular solution containing (in mM): 130 K-gluconate, 6.3 KCl, 0.5 EGTA, 1.0 MgCl2, 10 HEPES, 4.0 Mg-ATP, and 0.3 Na-GTP, 0.5% biocytin; pH was adjusted to 7.30 with KOH and osmolarity to 290 mOsm. Signals were sampled at 100 kHz with a MultiClamp 700B amplifier (Molecular Devices), filtered at 10 kHz, digitized using a DigiData 1550A, and acquired using pClamp10 software. Recordings were discarded if the cell had a resting membrane potential greater than −50 mV, or if access resistance was >30 MΩ and/or increased by >20% during the recording. We did not correct for liquid junction potential.

##### Electrophysiology data analysis

Analysis of intrinsic properties was performed using custom MATLAB (MathWorks) code with quality control using a manual confirmation in Clampfit (pCLAMP). Resting membrane potential (Vm) was calculated as the average value of a 600 ms window with no direct current injection. Input resistance (Rm) was calculated using the average response to a −100 pA hyperpolarizing current injection using Ohm’s law (Rm = ΔV/I). Quantification of single spike properties was performed using the first AP during the rheobase current injection. AP threshold was calculated as the value at which the derivative of the voltage (dV/dt) first reached 20 mV/ms. Spike height was calculated as the absolute maximum voltage value of a given AP. Spike amplitude was calculated as the difference between spike height and AP threshold. AP rise time was calculated as the time from the AP threshold to the absolute maximum voltage of the AP. AP half-width was defined as the width of the AP (in ms) at half-maximal amplitude. Afterhyperpolarization (AHP) amplitude was calculated as the difference between the minimum voltage value of the AHP and the AP threshold voltage. Rheobase was defined as the minimum current injection that elicited an AP using a 600 ms sweep at 25 pA intervals. The maximal instantaneous firing was calculated using the smallest inter-spike interval elicited during the current step protocol. Maximal steady-state firing was defined as the maximal mean firing frequency (FF) with a minimum requirement for a spike being the presence of a measurable AP threshold and the voltage overshooting 0 mV.

##### Single cell morphological reconstruction

After whole cell recordings, slices were transferred to cold 4% PFA overnight and then stored in 30% sucrose with 0.1 M phosphate buffer (PB). CUBIC clearing was performed as previously described.^61^ Briefly, slices were subjected to a freeze-thaw cycle then washed with 0.1 M PB and placed in CUBIC #1 solution (25% urea, 25% Quadrol, 15% Triton X-100 in ddH_2-_O) for 3 d. Slices were then washed again in 0.1 M PB, incubated in blocking buffer (5% NGS and 0.3% Triton X-100 in 0.1 M PB) for 1 hr at RT, then incubated in 1:500 dilution of streptavidin conjugated to Alexa-647 (ThermoFisher) in blocking buffer overnight at 4C. Slices were washed again in blocking buffer and then incubated in CUBIC #2 solution (25% urea, 50% sucrose, 10% triethanolamine in ddH_2_O) overnight at RT then mounted on glass slides in CUBIC #2 solution. Filled cells were imaged using a confocal microscope (Zeiss) with a 40X oil-immersion objective.

#### Electrophysiological characterization of BiPVe4 enhancer

##### Electrophysiology recordings and data analysis

Male and female C57BL6/J mice were injected with AAV-BiPVe4-GFP by retro-orbital injection of 2.0E+11 viral particles in 80uL per animal. Mice were anesthetized with 1.8–2% isoflurane. Titrated virus was injected into the retro-orbital sinus of the left eye with a 31G × 5/16 TW needle on a 3/10 mL insulin syringe. Mice were kept on a heating pad for the duration of the procedure until recovery and then returned to their home cage.

GFP-positive cells were targeted for somatic whole-cell recording in layer 2/3 region of the primary somatosensory, auditory and visual cortex by combining gradient-contrast video-microscopy with epifluorescent illumination on custom-built or commercial (Olympus) upright microscopes. Electrophysiological recordings were acquired using Multiclamp 700B amplifiers (Molecular Devices). Signals were filtered at 10 kHz and sampled at 50 kHz with the Digidata 1440B digitizer (Molecular Devices). For whole cell recordings, borosilicate patch pipettes were filled with an intracellular solution containing (in mM) 124 potassium gluconate, 2 KCl, 9 HEPES, 4 MgCl2, 4 NaATP, 3 L-Ascorbic Acid and 0.5 NaGTP. Pipette capacitance was neutralized in all recordings and electrode series resistance compensated using bridge balance in current clamp. Liquid junction potentials were uncorrected. Recordings had a series resistance < 20 MΩ. Action potential trains were initiated by somatic current injection (300 ms) normalized to the cellular capacitance in each recording measured immediately in voltage clamp after breakthrough66. For quantification of individual AP parameters, the 1st AP in a spike train at was analyzed at 12 pA/pF for all cells. Passive properties were determined by averaging the responses of several 100 ms long, −20pA steps during each recording. Data analysis was carried out using a custom written Python software.

#### Electrophysiological, morphological and optogenetic characterization of BiSSTe10 enhancer

##### Electrophysiology, optogenetic recordings and data analysis

After 3 weeks post-RO injection of the AAV-BiSSTe10-dTomato, animals were anesthetized with isoflurane followed by decapitation. The brain was quickly removed and immersed in ice-cold oxygenated sucrose cutting solution containing (in mM) 87 NaCl, 75 Sucrose, 2.5 KCl, 1.25 NaH2PO4, 26 NaHCO3, 10 Glucose, 1 CaCl2, 2 MgCl2 (pH=7.4). 300 μm thick coronal slices were cut using a Leica VT 1200S vibratome through the primary visual cortex. Slices recovered in a holding chamber with ACSF containing (in mM) 124 NaCl, 20 Glucose, 3 KCl, 1.2 NaH2PO4, 26 NaHCO3, 2 CaCl2, 1 MgCl2 (pH=7.4) at 34 °C for 30 minutes and at room temperature for at least 45 minutes prior to recording. For recording, slices were transferred to the recording chamber where dTomato-positive cells were identified under an upright differential interference contrast microscope (BX51WI) using a 40 × objective (water immersion lens, 0.9 numerical aperture). During recording slices were continuously perfused with oxygenated acsf (95%O2 / 5% CO2) at a flow rate of 1ml/min at room temperature.

For whole cell patch clamp recording, 5-7 MΩ pipettes were pulled from thick-walled borosilicate glass capillaries on a P-1000 Flaming Micropipette Puller (Sutter Instrument). The pipettes were filled with internal solution of the following composition: 130 K-Gluconate, 10 KCl, 10 HEPES, 0.2 EGTA, 4 MgATP, 0.3 NaGTP, 5 Phosphocreatine and 0.4% biocytin (pH = 7.3).

Recordings were performed using a Multiclamp 700B amplifier and digitized using Digidata 1440A using a sampling rate of 20KHz. Series resistance was continuously monitored during the recording and cells were discarded if the series resistance was > 40MOhms or if it changed by > 20% during the recording. For measurement of intrinsic properties, cells were held at −70mV. No liquid junction potential correction was made. Intrinsic cellular properties of visually identified dTomato-positive cells were measured in current clamp mode. Resting Membrane Potential (RMP) of the cell was recorded as mean membrane potential from a 1 min long recording in I = 0 mode. Input Resistance (IR) was computed using Ohm’s law (V = I/R) by finding the slope of the IV curve obtained using current injection from −10 to 10pA in steps of 5pA. Highest firing frequency was computed as the maximum firing frequency of a cell for any input current from −100 to 400pA. Rheobase was defined as the minimum input current to evoke firing from a cell. Action potential kinetics were computed from the first spike produced by the cell when a series of increasing input currents were injected. Spike threshold was defined as the membrane potential where dv/dt > 5mV/ms before spike initiation. Action potential half width is defined as the width of an action potential at half of the peak value from spike threshold. Spike Frequency Adaptation (SFA) Is calculated as ISIfirst/ISIlast. Fast hyperpolarization (fAHP) is calculated as the difference between spike threshold and minimum voltage after the spike within 10ms. mAHP is defined as difference between spike threshold and minimum voltage after the spike, from 10 to 20ms. All analysis was done using clampfit and Easy electrophysiology. Statistical analysis was done in GraphPad prism 10.

After recording with pipette solution containing 0.3-0.5% biocytin, the slices were fixed in 4% PFA overnight, then stored till further processing. After thorough washing with PBS1X, the slices were incubated in a blocking solution (10% normal donkey serum, 0.3% Triton X-100 in PBS1X) for 4 hours at room temperature. After thorough wash with PBS1X, slices were incubated with the same blocking solution supplemented with Alexa488-conjugated streptavidin overnight at 4C. The following day, sections were washed 3 times with PBS1x and mounted. Images were then acquired using a confocal microscope (Zeiss LSM 800) with a 40X objective (Plan-Apochromat 20x/0.8 M27).

Whole-cell patch-clamp recordings were obtained from unlabeled putative pyramidal neurons across layers. IPSC recordings were performed at a holding potential of 0 mV. For optogenetic stimulation, 470 nm light was transmitted from a collimated LED (Mightex) attached to the epifluorescence port of the upright microscope. 1 ms pulses of light were directed to the slice in the recording chamber via a mirror coupled to the 60x objective (N.A. = 1.0). Flashes were delivered every 15 s over a total of 15 trials. The LED output was driven by a transistor-transistor logic output from the Clampex software.

#### Electrophysiological characterization of BiSSTe4 enhancer

##### Slice preparation

Following ICV injection of AAV-BiSSTe4-dTomato, after 3 weeks of incubation, acute coronal slices (320 μm) containing primary sensory cortex barrel field (S1-BF) were prepared from adult C57BL6J mice, from postnatal days 20-25. Mice were deeply anesthetized with acute isoflurane exposition and slices prepared using a vibratome (Leica VT1200S) by slicing with the following cutting solution: 220 Sucrose, 11 Glucose, 2.5 KCl,1.25 NaH2PO4, 25 NaHCO3, 7 MgSO4, 0.5 CaCl2. Subsequently, the slices have been placed in ACSF solution at 34°C containing (in mM): 125 NaCl, 2.5 KCl, 2 CaCl2, 1 MgCl2, 1.25 NaH2PO4, 25 NaHCO3, 20 Glucose for 15 –20 min.

After the period of incubation, slices were kept at room temperature for a period of 6 h.

##### Electrophysiology recordings

Whole-cell patch-clamp recordings were performed close to physiological temperature (35°C) using a Multiclamp 700B amplifier (Molecular Devices) and fire-polished thick-walled glass patch electrodes 3.5–5 MOhm tip resistance. Patch pipettes were filled with the following intracellular solution (in mM): 135 K-gluconate, 5 KCl, 10 HEPES, 0.01 EGTA, 4 NaATP, 0.3 NaGTP (295 mOsm, pH adjusted to 7.3 using KOH). To fill the cells for the reconstructions we included in the intracellular solution 1.5% neurobiotin (Vector Laboratories). Recordings were not corrected for liquid junction potential. Series resistance was compensated online by balancing the bridge and compensating pipette capacitance. For all experiments data were discarded if series resistance, measured with a −10 mV pulse in voltage clamp configuration, was >35 ΜΩ. All recordings were low-pass filtered at 10 kHz and digitized at 100 kHz using a Digidata 1440A digitizer-Molecular Devices and acquired with pClamp 10 software.

All the intrinsic properties were measured in current-clamp conditions and calculated from 1-s-long current injections at resting membrane potential with a delta of 5pA. The resting membrane potential was calculated 5 minutes after break-in and Input resistance, calculated in MOhms, from 1 seconds long negative current injections of -10pA by using Ohm’s law. The action potential threshold was considered as the potential at which dV/dt reached 10 mV/ms. The adaptation index was determined by using this formula: 1/ (Freq-first/Freq-last)], with Freq-first and Freq-last representing the frequencies of the first two and the last two spikes at the maximum number of action potential detected. The number of spikes was automatically detected using a threshold at -10mV. The AHP duration was calculated as the time between AHP descending and ascending phase measured 1ms after the action potential descending phase and the most depolarized membrane potential following the AHP for a maximum of 200 ms.

##### Single cell morphological reconstruction

After performing electrophysiological recordings, brain slices were fixed in 4% PFA in 0.1M PB and kept overnight at 4°C and then washed with 1X PBS (4 times, 30 min). Slices were then incubated in streptavidin Alexa Fluor 647 conjugated (1:300; ThermoFisher Scientific) in 1X PBS 0.5% Triton overnight. Slices were subsequently washed with 1X PBS (4 times, 30 min) at RT. After washing the brain slices were processed for CUBIC (Clear, Unobstructed Brain/Body Imaging Cocktails and Computational analysis) clearing.^43^ Slices were immersed in CUBIC L solution reagent 8 hours at 37°C. After incubation, slices were washed with 1X PBS (3 times for 15 min) at RT to ensure complete removal of CUBIC reagent 1. Subsequently the slices have been immersed for 15 min in cubic R solution, washed briefly in PBS and mounted on a APTS covered coverslip (150 microns). Despite a complete clarification, we observed tissue swelling after 15 minutes exposure to CUBIC R solution.Thus, we mounted the slices for the acquisition with a spacer of 500 microns in the cubic mounting solution. Z-stacked images of filled neurons were acquired with a Zeiss LSM 800 Confocal Microscope using a 20X objective.

#### Electrophysiological, optogenetic and morphological characterization of BiVIPe4 enhancer

##### Preparation of brain slices and ex vivo electrophysiology and photo-stimulation

Viral injection of AAV-BiVIPe4-ChR2-mCherry was performed on mice postnatal days 0-3 (P0-3). P0-3 GIN-GFP pups (FVB-Tg(GadGFP)45704Swn/J) were briefly anesthetized on ice. A small volume of virus (30 nl) was unilaterally injected in the right somatosensory cortex (1.5 mm anterior and 1.5 mm lateral from lambda, depth of 0.1-0.4 mm) using Nanoject (Drummond).

Four to six weeks after virus injection, mice were anesthetized with isoflurane (5% isoflurane (vol/vol) in 100% oxygen), perfused intracardially with an ice-cold sucrose solution (in mM, 75 sucrose, 87 NaCl, 2.5 KCl, 26 NaHCO_3_, 1.25 NaH_2_PO_4_, 10 glucose, 0.5 CaCl_2_, and 2 MgSO_4_, saturated with 95% O2 and 5% CO_2_) and decapitated. Coronal slices of 300 μm were made using a vibratome (Leica Biosystems) and were stored in the same solution at 35°C for 30 min and at RT for an additional 30-45 min before recording.

Whole-cell patch clamp recordings were performed in oxygenated artificial cerebrospinal fluid (ACSF, in mM: 125 NaCl, 2.5 KCl, 26 NaHCO_3_, 1.25 NaH_2_PO_4_, 10 glucose, 2 CaCl_2_ and 1 MgCl_2_ equilibrated with 95% O_2_ and 5% CO_2_). Recordings were performed at 30°C–33°C. Electrodes (3-7 MΩ) were pulled from borosilicate glass capillary (1.5 mm OD). Membrane potentials were not corrected for the liquid junction potential. During patching, cell-attached seal resistances were > 1 GΩ. Once whole-cell configuration was achieved, uncompensated series resistance was usually 5 – 30 MΩ and only cells with stable series resistance (< 20% change throughout the recording) were used for analysis. Electrophysiological signals were collected with a Multiclamp 700B amplifier (Molecular Devices), low-pass filtered at 10 kHz, digitized at 20 kHz, and analyzed with pClamp10 (Molecular Devices).

To characterize the intrinsic membrane properties of mCherry expressing cells, a series of hyperpolarizing and depolarizing current steps (800 ms duration, 1 pA increments) were injected at 0.1 Hz under current-clamp configuration. All intrinsic properties were measured by holding cells at -70mV. The pipette intracellular solution was as follows (in mM): 130 potassium gluconate, 6.3 KCl, 0.5 EGTA, 10 HEPES, 5 sodium phosphocreatine, 4 Mg-ATP, 0.3 Na-GTP and 0.3% neurobiotin, pH 7.4 with KOH, 280-290 mOsm.

Inhibitory postsynaptic currents (IPSCs) were recorded from GFP positive cells under voltage-clamp configuration (holding cells at 0mV) using a cesium-based internal solution (in mM: 130 cesium gluconate, 6.3 KCl, 0.5 EGTA, 10 HEPES, 5 sodium phosphocreatine, 4 Mg-ATP, 0.3 Na-GTP adjusted to pH 7.2 with CsOH; 295 mOsm). For optogenetic stimulation, collimated light (590nm, 3 ms duration, 4.7 mW, CoolLED pE-300ultra system) was delivered at 10 s interval into the brain tissue through a 40X water-immersion objective.

##### Electrophysiological analysis

To characterize the intrinsic membrane properties of neurons, the following parameters were measured. The resting membrane potential (V_rest_ in mV) was measured with 0 pA current injection a few minutes after entering whole-cell configuration. All other properties were measured holding the cell at −70 mV. Input resistance (in MΩ) was calculated using Ohm’s law from averaged traces of 100 ms long negative current injections of −20 pA. Action potential (AP) threshold was calculated as the potential when voltage change over time was 10 mV/ms using the first observed spike. AP peak (in mV) was calculated as the time difference in potential from the spike peak to spike threshold. AP/spike half-width (in ms) was calculated as the difference in time between the ascending and descending phases of a putative spike at the voltage midpoint between the peak of spike and spike threshold. The adaptation ratio was calculated as the ratio of the last spike interval to the first spike interval under a positive current injection that elicited approximately 20 Hz firing. Afterhyperpolarization (AHP) amplitude was calculated as the difference between AP threshold and the most negative membrane potential after the AP, measured on the response to the smallest current step evoking an AP (Rheobase). To detect optogenetic stimulation-evoked IPSCs, 20 recorded sweeps were averaged, and the peak amplitude was measured.

##### Single cell morphological reconstruction

After performing electrophysiological recordings, brain slices were fixed in 4% PFA in 0.1M PB and kept overnight at 4°C and then kept in 20% sucrose. The brain slices were processed for CUBIC clearing.^43^ Slices were first washed with 0.1M PB (3 times for 10 min) at RT, followed by immersion in CUBIC reagent 1 for 2 days at 4°C. After 2 days of incubation, slices were washed with 0.1M PB (4 times for 30 min) at RT to ensure complete removal of CUBIC reagent 1. Slices were then incubated in fluorophore-conjugated streptavidin (1:500; ThermoFisher Scientific) in 0.1M PB (0.5% TritonX-100) overnight at 4°C. Slices were subsequently washed with 0.1M PB (4 times, 30 min) at RT and mounted with CUBIC reagent 1. Filled neurons were imaged with a Nikon A1R confocal microscope. Z-stacked images (each stack 1 μm) were acquired with a 40X oil-immersion objective.

##### Immunohistochemistry

Brain sections (50 μm thickness) were collected in 0.1 M PBS. Sections were rinsed in PBS (3 times for 10 min) and incubated in a blocking solution (10% normal serum, 1% BSA and 0.2% Triton X-100 in PBS) for 1 hour at room temperature. Brain sections were then incubated with primary antibodies (rabbit primary antibody for VIP, 1:200 dilution, ImmunoStar; guinea pig primary antibody for RFP, 1:500 dilution, Synaptic System) in diluted blocking solution (1:10 dilution in PBS) for 48 hr at 4°C. Brain sections were washed in PBS (3 times for 15 min) followed by 1 hour of secondary antibody incubation at RT. Alexa Fluor 647 conjugated secondary antibodies (anti-rabbit IgG, 1:200 dilution, Invitrogen) and CF555 conjugated secondary antibodies (anti-guinea pig IgG, 1:200 dilution, Sigma) were used to visualize fluorescence signals. After 1 hour of incubation with secondary antibodies, sections were washed in PBS (3 times for 15 min), mounted on slides and coverslipped with an antifade mounting medium containing DAPI (Thermo Fisher Scientific). Tissues were imaged (Zeiss, Axio Scan). Labeled neurons were automatically detected, manually confirmed, and subsequently quantified (Zeiss, Vision 4D Arivis Pro software). Confocal images were acquired (Nikon, A1R) for representative images.

#### Two-photon calcium imaging of BiVIPe4 targeting cells in mice

##### Surgeries

8-12 weeks old mice (C57BL/6J wild type from Charles River) were anesthetized with isoflurane (5% for induction, and 1-1.5% maintenance during surgery) and mounted on a stereotaxic frame (David Kopf Instruments, CA). Eye cream was applied to protect the eye (GenTeal, Alcon laboratories inc, Tx). After disinfecting the scalp, an incision was made, and the bone surface was cleaned. A circular 3 mm diameter biopsy was applied to remove the skull above the visual cortex in the left hemisphere (edges of biopsy are 1 mm from the midline and 0.5 mm anterior of lambda). AAV carrying BiVIPe4-gCaMP6f expression cassette were injected using glass pipette with 30 µm diameter tip mounted using Nanoject III (Drummond Instruments Ltd.) at a speed of 1 nl/s with a total volume of 100 nl per site (2 sites; 1 mm apart diagonally). After each injection, pipettes were left *in situ* for an additional 10 minutes to prevent backflow. The craniotomy was then sealed with a 3 mm glass coverslip that was glued to another 5 mm glass coverslip (Warner instruments) which in turn was glued to the skull with vetbond (Fisher scientific UK). A custom-built head-post was implanted over the exposed skull with ductile glue and cemented with C&B metabond (Parkell). Mice were monitored for a week after surgery and were injected with analgesia for three days (Carprofen; 0.1 mg / kg of body weight).

##### Two-photon calcium imaging

Data was acquired using a Brucker 2-photon microscope with a frame rate of 30 Hz with a 16X Nikon objective. Calcium imaging data were then preprocessed with Suite2p for motion correction and region of interest (ROI, i.e. neuron). Raw traces extracted from suite2p were further processed in python with custom scripts. To calculate Δf/f0 for each trace, a fluorescence baseline was determined by median filtering the trace with a window of 60s (1800 samples); the Δf/f0 was then computed by subtracting the baseline from the original trance and then dividing by the baseline.

##### Locomotion analysis

Three weeks after surgery, mice were head-fixed and run freely on a belt treadmill (Labmaker) with a built-in encoder for speed and direction. The encoder information was then converted to analog voltage and was sent to PyControl Breakout Board (Labmaker). The speed arbitrary unit was then converted to cm/s and smoothed post hoc.

Spontaneous activity (i.e. in darkness) was recorded for 2 minutes at the beginning of the recording session. Periods with running speed greater than 0.1 cm/s and a duration longer than 1 s were considered locomotion periods. Equal duration prior to the locomotion period was defined as stationary period. The mean change in fluorescence activity (Δf/f_0_) in the respective period was used in all locomotion analysis (Figure 3Q, 4P). The Locomotion modulation index (LMI) was computed to assess the responses of the neurons to locomotion.

##### Visual stimulation

Following the spontaneous activity period, drifting sinusoidal gratings at a spatial frequency of 0.04 cpd were presented to the mice. A total of 8 directions separated by 45 degrees were presented for 2 seconds with 10 seconds inter trial interval. Each condition consisted of 10 trials. Each neuron was recorded under low (25%) and high (100%) contrast conditions.

For analysis, a paired t test comparing the mean amplitude of Δf/f0 in the stimulation period to the mean amplitude 2 s prior to the visual stimulus was performed to determine if the neuron is responsive to each condition. For most of the neurons, we found no difference between the responses in trials with locomotion compared to stationary (likely due to the low number of locomotion trials), hence we pooled the two conditions in our analysis. To determine if the neuron is activated or suppressed, we compared the mean Δf/f0 during the stimulation of the preferred direction (i.e. highest mean response compared to other directions) relative to baseline (i.e. before stimulation). Neurons that increased their activity were considered activated, while neurons that decreased their activity were considered suppressed.

For orientation selectivity, we computed the global orientation selectivity index (gOSI), which considers responses to all orientations:

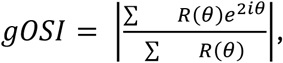

where *i* is the imaginary unit and *θ* is the angle of the moving directions. *R*(*θ*) is the response at direction *θ*.

##### RNAscope

RNA scope was performed on fixed frozen tissue. To prepare the sections, brains previously injected with the enhancer-driven GcaMP6f cassettes were perfused and stored overnight in 4% PFA at 4 C. Fixed tissue was then immersed in a graded series containing 10%, 20%, and 30 % sucrose solution and then froze in OCT at -80 °C. Brains were sectioned using a sliding microtome into 30 µm slices and mounted in Superfrost Plus Microscope Slides. RNAscope *in situ* hybridization assay was performed according to the manufacturer’s protocol using the RNAscope Multiplex Fluorescent Reagent Kit v2. Briefly, slides were baked in a HybEZ Oven at 60°C for 45 min, followed by post-fixation in 4% PFA for 15 min. Slides were dehydrated through 50%, 70%, 95%, 100%, and 100% ethanol for 5 min each at RT. Slides were treated with Hydrogen peroxide for 10 min at RT. Then, slides were incubated in 1x Target Retrieval Reagent for 5 min at >98 °C and dehydrated in 100% alcohol. Protease III was added to each section and incubated for 20 minutes at 40°C in the HybEZ oven. Pretreated sections were incubated with VIP-C1 probe (RNAscope® Probe-Mm-Vip) for 2h at 40°C. After hybridization, 3 amplification steps were performed, incubating the slides for 30 min at 40°C each. Followed by incubation with HRP-C1 solution for 15 min at 40°C. After washing, slides were incubated with Opal 570 fluorophore for 30 min, followed by HRP blocking for 20 min at 40°C.

Following the *in situ* hybridization, sections were immunostained to amplify the expression of BiVIPe4-GcaMP6f virally labeled cells. Sections were incubated for 1 h in 10 % Normal Goat Serum (NGS) in TBS-0.1%BSA at RT. Then, slides were incubated with primary antibody chicken anti-GFP (1:2000) In TBS-0.1%BSA for 2 hours at RT. Slides were washed twice with TBST and incubated with the secondary antibody Alexa Fluor 488 goat anti-chicken (1:500) for 1 hour at RT. Slides were then washed with TBST twice and incubated with DAPI for 30s. Finally, slides were mounted using Prolong Gold Antifade mounting medium. Images were acquired using a Leica Sp8 confocal microscope with a 40x objective.

#### Electrophysiological characterization of BiLAMP5e3 enhancer

##### Slice preparation

Following 3-4 weeks from intracranial injection, acute coronal slices (320 μm) containing primary sensory cortex barrel field (S1-BF) were prepared from adult C57BL6J mice, from postnatal days 45-75. Mice were deeply anesthetized with acute isoflurane exposition and perfused transcardially with ice-cold cutting solution containing the following (in mM): 220 Sucrose, 11 Glucose, 2.5 KCl,1.25 NaH2PO4, 25 NaHCO3, 7 MgSO4, 0.5 CaCl2. After perfusion the brain was quickly removed, and slices prepared using a vibratome (Leica VT1200S). Slices to ACSF solution at 34°C containing (in mM): 125 NaCl, 2.5 KCl, 2 CaCl2, 1 MgCl2, 1.25 NaH2PO4, 25 NaHCO3, 20 Glucose for 15–20 min. After the period of incubation, slices were kept at room temperature for a period of 6 h.

##### Electrophysiology recordings

Whole-cell patch-clamp recordings were performed close to physiological temperature (35°C) using a Multiclamp 700B amplifier (Molecular Devices) and fire-polished thick-walled glass patch electrodes 3.5–5 MOhm tip resistance. Patch pipettes were filled with the following intracellular solution (in mM): 135 K-gluconate, 5 KCl, 10 HEPES, 0.01 EGTA, 4 NaATP, 0.3 NaGTP (295 mOsm, pH adjusted to 7.3 using KOH). Recordings were not corrected for liquid junction potential. Series resistance was compensated online by balancing the bridge and compensating pipette capacitance. For all experiments data were discarded if series resistance, measured with a −10 mV pulse in voltage clamp configuration, was >35 ΜΩ. All recordings were low-pass filtered at 10 kHz and digitized at 100 kHz using a Digidata 1440A digitizer-Molecular Devices and acquired with pClamp 10 software.

All the intrinsic properties were measured in current-clamp conditions and calculated from 1-s-long current injections at resting membrane potential with a delta of 5pA. The resting membrane potential was calculated 5 minutes after break-in and Input resistance, calculated in MOhms, from 1 seconds long negative current injections of -10pA by using Ohm’s law. The amplitude of voltage of Sag current (%) was measured using hyperpolarization induced by −100 pA and calculated as 100 × (Vsag minimum − Vsteady-state)/(Vsag minimum − Vholding) as reported in Schuman et al. 2018. The depolarizing hump and depolarizing ramp ratio were calculated on the current step just before the action potential threshold. This ratio was defined as the measured amplitude (in millivolts) relative to the difference between the resting membrane potential and the step potential. The depolarizing hump was determined as the average amplitude between 40–90 ms while the depolarizing ramp was calculated as the average amplitude between 940–990 ms both measured from the one second depolarizing step. Spike were detected increasing the current steps of 5pA. The action potential threshold was considered as the potential at which dV/dt reached 10 mV/ms. The AP amplitude (in millivolts) was calculated as the difference in potential from the spike peak to the spike threshold. The AP half-width (in milliseconds) was calculated as the time interval between rise and decay phase measured at 50% of amplitude. The adaptation index was determined by dividing the number of spikes in the last 500 ms by the number of spikes in the first 500 ms of the 1 second step during a positive current injection that induced a firing rate of 20–25 Hz. The time to the first spike (in milliseconds) was determined by measuring the time difference between the onset of the positive current injection and the peak of the first action potential. The afterhyperpolarization (AHP) was calculated as the difference in potential between the threshold of the spike and the minimum potential measured after the spike.The AHP half-width as the time between AHP descending and ascending phase measured 1ms after the action potential descending phase and the most depolarized membrane potential following the AHP for a maximum of 200 ms.

#### Optogenetic characterization of BiCHATe27 enhancer in mice

##### Surgeries

Mice (P55-P60) were anesthetized with isoflurane 5% in O_2_ and moved to a stereotactic surgery rig, keeping isoflurane levels at 1.5%-2% and maintained throughout the procedure. At the start of the surgery, Buprenex (0.01 mg/ml; dosage 0.05 ml/10 g) and Meloxicam (0.2 mg/ml; dosage 0.1 ml/10 g) were administered via subcutaneous injections. A small incision was made in the skin to expose the skull. The skull was leveled, and burr holes were made in two using a dental drill (MH170, Foredom Electric Co.) at the corresponding coordinates to target the left cholinergic basal forebrain (site 1: AP = -0.2 mm; ML = 2.5 mm; DV = -4 mm; site 2: AP = -0.8 mm; ML = 2.5 mm; DV = -3.25 mm). Next, AAV-BiCHATe27-ChR2-mCherry was injected in both sites (1.3E+12 – 300 nl per injection site at a speed of 12 nl/min). All injections were performed using glass capillaries pulled with a micropipette puller (P-97, Sutter Instrument) and using a motorized programmable injector (Nanoject III, 3-000-207, Drummond Scientific Co.).

##### In vitro whole-cell recordings

3-4 weeks after surgery, mice (P76-88) were anesthetized with isoflurane 5% in O_2_ and Fatal Plus administered intraperitoneally (18 mg/ml; dosage 0.1 ml/10 g). Following anesthesia, mice were perfused with ice cold slicing artificial cerebrospinal fluid (ACSF) containing (in mM): 160 sucrose, 28 NaHCO_3_, 2.5 KCl, 1.25 NaH_2_PO_4_, 7.25 glucose, 20 HEPES, 3 Na-pyruvate, 3 Na-ascorbate, 7.5 MgCl_2_, 1 CaCl_2_. Following perfusion, mice were decapitated, the brain quickly removed and thalamocortical slices (300 µm) obtained at an angle of 15° from horizontal view^35^ were collected in ice cold slicing ACSF on a vibrating microtome (Leica Microsystems; VT1200S). Slices were placed in a chamber for 30-35 minutes at 35° C in recovery ACSF, containing (in mM): 92 NaCl, 28.5 NaHCO_3_, 2.5 KCl, 1.2 NaH_2_PO_4_, 25 glucose, 20 HEPES, 3 Na-pyruvate, 5 Na-ascorbate, 4 MgCl_2_, 2 CaCl_2_. After recovery, slices were transferred to a chamber with recording ACSF, containing (in mM): 125 NaCl, 2.5 KCl, 1.25 NaH_2_PO_4_, 25 NaHCO_3_, 25 glucose, 1 MgCl_2_, 2 CaCl_2_^24^ and maintained for at least one hour at room temperature. Slices were moved to the recording chamber and maintained under continuous superfusion with recording ACSF at near-physiological temperature (31-33° C) throughout all recordings. All solutions were continuously bubbled with CO_2_-O_2_ (95%-5%). Patch recording pipettes (3–5 MΩ) were obtained from borosilicate glass capillaries pulled with a micropipette puller (P-97, Sutter Instrument) and filled with current-clamp internal solution containing (in mM): 5 KCl, 127.5 K-gluconate, 10 HEPES, 2 MgCl_2_, 0.6 EGTA, 2 Mg-ATP, 0.3 Na-GTP, 5 Na_2_-phosphocreatine, pH 7.3 with KOH.

Whole-cell current clamp recordings were obtained from layer 1 (L1) interneurons in the auditory cortex, identified under IR-DIC. Excitatory postsynaptic potentials (EPSPs) were obtained in L1 interneurons upon light stimulation to evoke ACh release from cholinergic basal forebrain axons in the auditory cortex. Light-evoked responses to a wide-field 470 nm LED pulse (CoolLED, pE-100; 5 ms, 0.1Hz, ∼14 mW/mm^2^) were recorded in current-clamp configuration in the presence of 20 μM DNQX (Sigma) and 50 μM AP5 (Tocris). The cholinergic nature of these inputs was confirmed by application of 10 uM Dihydro-β-erythroidine hydrobromide (DhβE, Tocris) and 10 uM Methyllycaconitine (MLA, Sigma). Data were acquired from cells with initial series resistance below 20 MΩ (60% compensation), collected at a sampling rate of 10 kHz with a Multiclamp 700B amplifier (Molecular Devices), low-pass filtered at 3 kHz and digitized with a digital-to-analog converter board (National Instruments, NI-USB-6343). Custom-designed Matlab software was used for data acquisition and analysis. Experiments were carried out in a motorized upright microscope (Scientifica, SliceScope Pro 1000) coupled to a CCD camera (Hamamatsu Photonics, Orca Flash 4).

##### Immunohistochemistry and image acquisition

Mice were anesthetized with isoflurane 5% in O_2_ and Fatal Plus administered intraperitoneally (18 mg/ml; dosage 0.1 ml/10 g) and subsequently perfused with 4% PFA in 0.1 M PB. Brains were incubated in PBS with azide overnight. The following day, brains were washed 3 times in PBS1X and sliced into coronal sections (50 µm) using a vibrating microtome (Leica Microsystems; VT1000S). Floating slices were maintained in PBS with azide at 4°C. Slices spanning the A1 region and the cholinergic basal forebrain were immunostained (primary antibodies: anti-ChAT (Sigma) and anti-mCherry (Kerafast Inc.), both at 1:100; secondary antibodies: anti-goat-488 and anti-rabbit-647 (Fisher) both at 1:500), treated with 60 nM DAPI and mounted. Images were acquired on a confocal microscope at or 63X (Leica Microsystems, SP8).

#### *In vivo* optogenetic characterization of BiPVe3, BiPVe4, BiSSTe4 and BiLAMP5 enhancers in mice

##### Mice

All animal handling and maintenance was performed according to the regulations of the Institutional Animal Care and Use Committee of Albert Einstein College of Medicine (Protocol # 00001393). Both adult male and female C56BL/6 mice above P50 were used in this study. Animals were kept under a 12h light/dark cycle (lights on 7am) and were maintained under standard conditions.

##### Viral injections

Mouse V1 injections: Mice were anesthetized with isoflurane (5% by volume for induction and between 1 and 2% for maintenance), placed on a stereotaxic frame, and kept warm with a closed loop heating pad. For pain management, animals were given meloxicam at 2.5 mg/kg and local lidocaine on the scalp. After a single midline incision of the skin, we minimized brain damage by performing burr holes, keeping a thin layer of the bone intact where the glass micropipettes could penetrate. Local injection in adult mice were performed by stereotactically guided injections in the primary visual cortex with the following coordinates -3.2 mm posterior, +/- 2.7 mm lateral, 0.25/0.50 mm ventral relative to Bregma with 500 nL of the following AAVs: BiPVe4-ChR2-mCherry, BiPVe3-Chr2-mCherry, BiSSTe4-ChR2-mCherry and BiLAPM5e3-ChR2-mCherry. Postoperative monitoring was performed for five days post injection. Injections were performed using a RWD R-480 Nanoliter Microinjection Pump system at a rate of 2nL/seconds through glass micropipettes that were pulled and then ground to bevel with 40µm diameter with a Narishige diamond wheel. Mice were left to build expression for a minimum of 4 weeks before being recorded. Finally, for electrophysiological recordings, a tungsten ground wire attached to a gold pin was inserted into the cerebellum and a headpost was added. Everything was cemented onto the skull with Optibond or Super-Bond and dental cement. Habituation and head restraint animals were briefly habituated to handling for several days prior to surgery. Animals were allowed to recover for 2 days before continuing habituation to handling. After a minimum of a week post injections, animals were gradually exposed to the head fixation apparatus (https://doi.org/10.25378/janelia.24691311) that consists of a low-friction rodent-driven belt treadmill. Head fixation times were gradually increased over the course of 1 week until animals were comfortable with multi-hour head fixation sessions. We ensured that the treadmill was oriented horizontally without slope with appropriately adjusted distance between the head fixation level and treadmill and that the set up was thoroughly cleaned to minimize odors from other animals.

##### Electrophysiological recordings in mice

AAV-transduced mice were administered dexamethasone 4mg/kg 1h prior craniotomy. A ∼3mm craniotomy was made over the injection site (V1). A 64 channel linear silicon probe (H3 probe, Cambridge Neurotech) physically coupled to a tapered optical fiber was slowly (1µm/sec, over ∼20 minutes) inserted into V1. The craniotomy was kept moisturized during the recording by placing a small volume of silicone oil on the surface of the brain. Between recording days, the craniotomy was protected with Kwik-cast silicone elastomer (World Precision Instruments). Mice were recorded once per day for 3-4 days. Recordings were split into blocks of spontaneous activity and visually evoked activity. Blue light was provided by a fiber coupled LED (MF470F4, Thorlabs). Signals to control LED light intensity were generated using custom written MATLAB code and delivered via NI-DAQ card (NI-PCIe-6323) to the LED control box (LEDD1B, Thorlabs). No photoelectric artifacts were detected in our recordings. Data was acquired using an Intan RHD2000 interface board at 20kHz.

##### Data analysis

We calculate percent inhibited by calculating the difference in mean firing rate one second before laser onset and one second after laser onset divided by the sum of the mean firing rates.

If the percent inhibited is 100, it represents a complete inhibition of the cells; If 0, the cell is not inhibited.

#### Electrophysiological, optogenetic, morphological, and IHC characterization of enhancers in rhesus macaque and mouse striatum

##### Surgery and tissue collection in mice

All experiments were conducted in accordance with animal protocols approved by the National Institutes of Health. Tissue was obtained from three 2-4 month old mice (WT, C57BL/6J) injected with 200ul of AAV-BiCHATe27 in the left/right striatum (AP: 0.98 mm, ML: +/-1.25 mm, DV: 4.63/3.36 mm) 2 weeks prior to tissue harvest. Mice were anesthetized with isoflurane and the brain dissected out in ice-cold saline solution containing (in mM): 90 sucrose, 80 NaCl 3.5 KCl, 24 NaHCO3, 1.25 NaH2PO4, 10 glucose, and saturated with 95% O2 and 5% CO2 (osmolarity 310-320 Osm). Coronal slices containing striatum (300 um) were cut using a VT-1000S vibratome (Leica Microsystems, Bannockburn, IL) and incubated in the above solution at 35C for recovery (30 min) then kept at room temperature until use. Slices were transferred to an upright microscope and perfused with heated (30C) extracellular solution containing (in mM): 130 NaCl, 24 NaHCO3, 10 glucose, 3.5 KCl, 1.25 NaH2PO4, and saturated with 95% O2 and 5% CO2 (osmolarity 300 Osm).

##### Animals (rhesus macaque)

All experiments were performed in accordance with the ILAR Guide for the Care and Use of Laboratory Animals and were conducted under an Animal Study Protocol approved by the ACUC at the National Institute of Mental Health. All procedures adhered to the applicable Federal and local laws, regulations, and standards, including the Animal Welfare Act and Regulations and Public Health Service policy (PHS2002). Tissue was obtained from 8 adult rhesus macaques (3 female), aged 5 – 19.5years, as part of the NIH Comparative Brain Physiology Consortium.

##### Surgery and tissue collection for rhesus macaque

Viral injections targeted the hippocampus, posterior striatum, and motor cortex using stereotaxic coordinates derived from MRI and delivered using a needle guide for enhanced accuracy.^62^ Surgeries were performed under aseptic conditions in a fully equipped operating suite. Within the hippocampus, 10-20 μL of virus was injected at each of 2 locations spaced approximately 2 mm apart in the antero-posterior plane, caudal to the level of the uncus. Within the posterior striatum and motor cortex, 10 μL of virus was injected at each of 4-5 locations spaced approximately 2 mm apart.

AAV.PHP.eB-BiSSTe4-ChR2-mCherry and AAV.PHP.eB-BiVIPe4-ChR2-mCherry viruses were injected into the right/left hippocampus of one female macaque (age 16.9 years). AAV.PHP.eB-BiCHATe27-dTomato virus was injected into the left posterior striatum of two female macaques (ages 7.4, 16.9 years). AAV.PHP.eB-BiSSTe10-dTomato virus was injected into the right hippocampus of three male macaques (ages 9.7, 10, 8.7 years). AAV.PHP.eB-BiSSTe4-dTomato virus was injected into the right hippocampus of one female macaque (age 15.9 years), while AAV.PHP.eB-BiPVe4-dTomato and AAV.PHP.eB-BiPVe3-dTomato were injected into the left/right motor cortex of one male macaque (age 5.1 years).

For brain extraction, 5-8 weeks after virus injection, animals were sedated with ketamine/midazolam (ketamine 5-15 mg/kg, midazolam 0.05-0.3 mg/kg) and maintained on isoflurane. A deep level of anesthesia was verified by an absence of response to toe-pinch and absence of response to corneal reflex. Prior to brain removal and blocking the animals were transcardially perfused with ice-cold sucrose-substituted artificial cerebrospinal fluid (aCSF) containing in mM: 90 Sucrose, 80 NaCl, 3.5 KCl, 1.25 NaH2PO4, 24 NaHCO3, 10 Glucose, 0.5 CaCl, 4.5 MgCl2 saturated with carbogen (95% O2, 5% CO2), with osmolarity 310-320 Osm.

##### Characterization of intrinsic membrane and firing properties in mice and macaques

Borosilicate glass pipettes were pulled to 3-5 MOhm resistance and filled with standard internal K-Gluconate solution containing in mM: 150 K Gluconate, 0.5 EGTA, 3 MgCl2, 10 HEPES, 2 ATP Mg, 0.3 GTP Na2, and 0.3% biocytin (pH corrected to 7.2 with KOH, 285-290 Osm). A Multiclamp 700B amplifier (Molecular Devices) was used for physiology recordings, signals were digitized at 20 kHz (Digidata 1440A, filtered at 3 kHz). Signals were recorded using pClamp 10.7 (Molecular Devices). Intrinsic membrane and firing properties were measured in Clampfit software (Molecular Devices).

Intrinsic membrane and firing properties were measured as follows: resting membrane potential (RMP) was measured in cell attached voltage clamp during depolarizing 100ms voltage ramps from (+100 to -200 mV) every 5 seconds or in whole-cell current clamp when I=0. Spontaneous action potential frequency (sAP freq.) was measured in cell-attached patch clamp by taking the frequency of action potentials recorded in a 10-30s sweep. Membrane time constant (tau) was measured by calculating the mean of 20 repeated 400ms hyperpolarizing current pulses of -20pA and fitting the voltage response with an exponential function. Input resistance (Rin) was determined by taking the slope of a linear regression of the change in voltage in response to 20 increasing current pulses (2s long from -50pA with a 5pA increments). The sag ratio measures the amount of hyperpolarization sag, an indication of hyperpolarization-activated cationic current (Ih). Sag ratio is determined during 800ms negative current pulses by measuring the maximum hyperpolarization voltage deflection (Vpeak) from baseline (Vrest) at the onset of the current pulse, and the steady state voltage deflection (Vss) in the last 200 ms of the current pulse. Sag ratio is calculated as Vrest – Vss / Vrest – Vpeak during current steps where Vpeak = -80mV. Action potential threshold (AP threshold) was measured as the voltage (mV) at which the slope is 10mV/ms during a depolarizing current injection. Action potential half-width (AP Halfwidth) is the length of time (ms) an action potential reaches half of the maximum voltage amplitude. Maximum firing frequency is the maximum action potential frequency a cell can be driven to within 1s depolarizing current pulses. After-hyperpolarization pulse amplitude (AHP amp.) is measured as the peak hyperpolarization voltage deflection from AP threshold at the offset of a depolarizing pulse. ISI accommodation ratio (or inter-spike interval accommodation ratio) is the initial divided by the final inter-spike interval (ms) during the maximum firing frequency action potential train. Finally, half-width accommodation ratio is the half-width measured from the last action potential during the maximum firing frequency train divided by the half-width from the first action potential. All electrophysiological properties and measurements were taken in Clampfit analysis software.

##### Immunohistochemistry in mice and macaques

Following electrophysiological recording and biocytin filling of recorded cells, macaque slices were drop-fixed in 4% paraformaldehyde (PFA) for at least 24 hours (at 4C) prior to transferring to 30% sucrose in 1XPB solution until slices dropped and were re-sectioned at 50 um using a freezing microtome. Re-sectioned floating slices were washed in 1XPB at room temperature (RT) prior to blocking for 2-4 hours at RT in 1XPB + 10% donkey serum + 0.5% TritonX-100. Slices were then incubated at 4C for 16-64 hours in a primary solution containing diluted antibodies and 1XPB + 1% donkey serum + 0.1% TritonX-100 (carrier solution). Slices were then washed at RT in carrier solution before incubating for 2 hours at RT in secondary solution containing diluted secondary antibodies and carrier solution. Finally, slices were washed in 1XPB at RT prior to mounting onto gelatin-coated slides in Mowiol. Mouse slices underwent the same IHC protocol as macaque but replaced 1XPB with 1X PBS. Antibody concentrations were as follows: guinea pig anti-RFP (1:1000, Synaptic Systems), rabbit anti-somatostatin (1:1000, Abcam), rabbit anti-parvalbumin (1:1000, Abcam), rabbit anti-calretinin (1:1000, Abcam), mouse anti-parvalbumin (1:1000, Sigma), Rabbit anti-nNos, (Sigma, 1:500) and goat anti-ChAT (1:500, Sigma,). Biocytin filled cells were stained with Alexa Fluor 488 Streptavidin (1:500, ThermoFisher) for cell reconstructions. Images were captured using Zeiss LSM900 confocal or Olympus VS200 Slideview. Cell counts were performed in ImageJ/Fiji.

#### Characterization of BiVIPe4 enhancer in human tissue

Surgical human tissue was collected from the temporal lobe of two patients (47M and 41F) undergoing temporal lobectomies for epilepsy resection. The specimen was processed into 300um sections and maintained on ex vivo air-interface culture inserts following published protocol.^63^ The rAAV/PHP.eB-BiVIPe4-ChR2-mCherry viral vector was introduced on the day of tissue collection by directly dropping 5ul viral vector (2E+12 vg/ml) onto the slice and fixed after 6 days of *ex vivo* culture. Primary antibody used were chicken anti-GFP 1:500 (AVES Lab), mouse anti-Calretinin 1:500 (Swant), and rat anti-RFP 1:500 (Chromotek), followed by secondary antibodies goat anti-chicken 488 (1:1000), donkey anti-mouse 488 (1:1000), and donkey anti-rat 594 (1:1000). The tissue was mounted with ProLong Diamond Antifade Mountant (Thermo Fisher). Imaging was performed on a Leica SP8 confocal.

## REFERENCES

1. 1. Yao, Z., van Velthoven, C.T.J., Nguyen, T.N., Goldy, J., Sedeno-Cortes, A.E., Baftizadeh, F., Bertagnolli, D., Casper, T., Chiang, M., Crichton, K., et al. (2021). A taxonomy of transcriptomic cell types across the isocortex and hippocampal formation. Cell 184, 3222–3241 e26. 10.1016/j.cell.2021.04.021.

2. Tasic, B., Yao, Z., Graybuck, L.T., Smith, K.A., Nguyen, T.N., Bertagnolli, D., Goldy, J., Garren, E., Economo, M.N., Viswanathan, S., et al. (2018). Shared and distinct transcriptomic cell types across neocortical areas. Nature 563, 72–78. 10.1038/s41586-018-0654-5.

3. Gouwens, N.W., Sorensen, S.A., Baftizadeh, F., Budzillo, A., Lee, B.R., Jarsky, T., Alfiler, L., Baker, K., Barkan, E., Berry, K., et al. (2020). Integrated Morphoelectric and Transcriptomic Classification of Cortical GABAergic Cells. Cell 183, 935–953 e19. 10.1016/j.cell.2020.09.057.

4. Wu, S.J., Sevier, E., Dwivedi, D., Saldi, G.A., Hairston, A., Yu, S., Abbott, L., Choi, D.H., Sherer, M., Qiu, Y., et al. (2023). Cortical somatostatin interneuron subtypes form cell-type-specific circuits. Neuron 111, 2675–2692 e9. 10.1016/j.neuron.2023.05.032.

5. Dimidschstein, J., Chen, Q., Tremblay, R., Rogers, S.L., Saldi, G.A., Guo, L., Xu, Q., Liu, R., Lu, C., Chu, J., et al. (2016). A viral strategy for targeting and manipulating interneurons across vertebrate species. Nat Neurosci 19, 1743–1749. 10.1038/nn.4430.

6. Vormstein-Schneider, D., Lin, J.D., Pelkey, K.A., Chittajallu, R., Guo, B., Arias-Garcia, M.A., Allaway, K., Sakopoulos, S., Schneider, G., Stevenson, O., et al. (2020). Viral manipulation of functionally distinct interneurons in mice, non-human primates and humans. Nat Neurosci 23, 1629–1636. 10.1038/s41593-020-0692-9.

7. Graybuck, L.T., Daigle, T.L., Sedeno-Cortes, A.E., Walker, M., Kalmbach, B., Lenz, G.H., Morin, E., Nguyen, T.N., Garren, E., Bendrick, J.L., et al. (2021). Enhancer viruses for combinatorial cell-subclass-specific labeling. Neuron 109, 1449–1464 e13. 10.1016/j.neuron.2021.03.011.

8. Mich, J.K., Graybuck, L.T., Hess, E.E., Mahoney, J.T., Kojima, Y., Ding, Y., Somasundaram, S., Miller, J.A., Kalmbach, B.E., Radaelli, C., et al. (2021). Functional enhancer elements drive subclass-selective expression from mouse to primate neocortex. Cell Rep 34, 108754. 10.1016/j.celrep.2021.108754.

9. Ben-Simon, Y., Hooper, M., Narayan, S., Daigle, T., Dwivedi, D., Way, S.W., Oster, A., Stafford, D.A., Mich, J.K., Taormina, M.J., et al. (2024). A suite of enhancer AAVs and transgenic mouse lines for genetic access to cortical cell types. bioRxiv. 10.1101/2024.06.10.597244.

10. Hrvatin, S., Tzeng, C.P., Nagy, M.A., Stroud, H., Koutsioumpa, C., Wilcox, O.F., Assad, E.G., Green, J., Harvey, C.D., Griffith, E.C., et al. (2019). A scalable platform for the development of cell-type-specific viral drivers. Elife 8. 10.7554/eLife.48089.

11. Hunker, A.C., Wirthlin, M.E., Gill, G., Johansen, N.J., Hooper, M., Omstead, V., Taskin, N., Weed, N., Vargas, S., Bendrick, J.L., et al. (2024). Enhancer AAV toolbox for accessing and perturbing striatal cell types and circuits. Preprint, 10.1101/2024.09.27.615553 https://doi.org/10.1101/2024.09.27.615553.

12. Buenrostro, J.D., Wu, B., Litzenburger, U.M., Ruff, D., Gonzales, M.L., Snyder, M.P., Chang, H.Y., and Greenleaf, W.J. (2015). Single-cell chromatin accessibility reveals principles of regulatory variation. Nature 523, 486–490. 10.1038/nature14590.

13. Dai, M., Pei, X., and Wang, X.J. (2022). Accurate and fast cell marker gene identification with COSG. Brief Bioinform 23. 10.1093/bib/bbab579.

14. Chan, K.Y., Jang, M.J., Yoo, B.B., Greenbaum, A., Ravi, N., Wu, W.L., Sanchez-Guardado, L., Lois, C., Mazmanian, S.K., Deverman, B.E., et al. (2017). Engineered AAVs for efficient noninvasive gene delivery to the central and peripheral nervous systems. Nat Neurosci 20, 1172–1179. 10.1038/nn.4593.

15. Taniguchi, H., Lu, J., and Huang, Z.J. (2013). The spatial and temporal origin of chandelier cells in mouse neocortex. Science (1979) 339. 10.1126/science.1227622.

16. Paul, A., Crow, M., Raudales, R., He, M., Gillis, J., and Huang, Z.J. (2017). Transcriptional Architecture of Synaptic Communication Delineates GABAergic Neuron Identity. Cell 171, 522–539 e20. 10.1016/j.cell.2017.08.032.

17. Caccavano, A.P., Vlachos, A., McLean, N., Kimmel, S., Kim, J.H., Vargish, G., Mahadevan, V., Hewitt, L., Rossi, A.M., Spineux, I., et al. (2024). Divergent opioid-mediated suppression of inhibition between hippocampus and neocortex across species and development. Preprint, 10.1101/2024.01.20.576455 https://doi.org/10.1101/2024.01.20.576455.

18. Gutzmann, A., Ergül, N., Grossmann, R., Schultz, C., Wahle, P., and Engelhardt, M. (2014). A period of structural plasticity at the axon initial segment in developing visual cortex. Front Neuroanat 8. 10.3389/fnana.2014.00011.

19. Tremblay, R., Lee, S., and Rudy, B. (2016). GABAergic Interneurons in the Neocortex: From Cellular Properties to Circuits. Neuron 91, 260–292. 10.1016/j.neuron.2016.06.033.

20. Helm, J., Akgul, G., and Wollmuth, L.P. (2013). Subgroups of parvalbumin-expressing interneurons in layers 2/3 of the visual cortex. J Neurophysiol 109, 1600–1613. 10.1152/jn.00782.2012.

21. Woodruff, A., Xu, Q., Anderson, S.A., and Yuste, R. (2009). Depolarizing effect of neocortical chandelier neurons. Front Neural Circuits 3, 15. 10.3389/neuro.04.015.2009.

22. Nigro, M.J., Hashikawa-Yamasaki, Y., and Rudy, B. (2018). Diversity and connectivity of layer 5 somatostatin-expressing interneurons in the mouse barrel cortex. Journal of Neuroscience 38. 10.1523/JNEUROSCI.2415-17.2017.

23. Hilscher, M.M., Leão, R.N., Edwards, S.J., Leão, K.E., and Kullander, K. (2017). Chrna2-Martinotti Cells Synchronize Layer 5 Type A Pyramidal Cells via Rebound Excitation. PLoS Biol 15. 10.1371/journal.pbio.2001392.

24. Butt, S.J., Fuccillo, M., Nery, S., Noctor, S., Kriegstein, A., Corbin, J.G., and Fishell, G. (2005). The temporal and spatial origins of cortical interneurons predict their physiological subtype. Neuron 48, 591–604. 10.1016/j.neuron.2005.09.034.

25. Miyoshi, G., Butt, S.J., Takebayashi, H., and Fishell, G. (2007). Physiologically distinct temporal cohorts of cortical interneurons arise from telencephalic Olig2-expressing precursors. J Neurosci 27, 7786–7798. 10.1523/JNEUROSCI.1807-07.2007.

26. Miyoshi, G., Hjerling-Leffler, J., Karayannis, T., Sousa, V.H., Butt, S.J.B., Battiste, J., Johnson, J.E., Machold, R.P., and Fishell, G. (2010). Genetic fate mapping reveals that the caudal ganglionic eminence produces a large and diverse population of superficial cortical interneurons. J Neurosci 30, 1582–1594. 10.1523/JNEUROSCI.4515-09.2010.

27. Lee, S., Kruglikov, I., Huang, Z.J., Fishell, G., and Rudy, B. (2013). A disinhibitory circuit mediates motor integration in the somatosensory cortex. Nat Neurosci 16, 1662–1670. 10.1038/nn.3544.

28. Fishell, G., and Kepecs, A. (2020). Interneuron Types as Attractors and Controllers. Annu Rev Neurosci 43, 1–30. 10.1146/annurev-neuro-070918-050421.

29. Huang, S., Rizzo, D., Wu, S.J., Xu, Q., Ziane, L., Alghamdi, N., Stafford, D.A., Daigle, T.L., Tasic, B., Zeng, H., et al. (2024). Neurogliaform Cells Exhibit Laminar-specific Responses in the Visual Cortex and Modulate Behavioral State-dependent Cortical Activity. bioRxiv. 10.1101/2024.06.05.597539.

30. Pronneke, A., Scheuer, B., Wagener, R.J., Mock, M., Witte, M., and Staiger, J.F. (2015). Characterizing VIP Neurons in the Barrel Cortex of VIPcre/tdTomato Mice Reveals Layer-Specific Differences. Cereb Cortex 25, 4854–4868. 10.1093/cercor/bhv202.

31. Lee, S., Hjerling-Leffler, J., Zagha, E., Fishell, G., and Rudy, B. (2010). The largest group of superficial neocortical GABAergic interneurons expresses ionotropic serotonin receptors. J Neurosci 30, 16796–16808. 10.1523/JNEUROSCI.1869-10.2010.

32. Pfeffer, C.K., Xue, M., He, M., Huang, Z.J., and Scanziani, M. (2013). Inhibition of inhibition in visual cortex: the logic of connections between molecularly distinct interneurons. Nat Neurosci 16, 1068–1076. 10.1038/nn.3446.

33. Pi, H.J., Hangya, B., Kvitsiani, D., Sanders, J.I., Huang, Z.J., and Kepecs, A. (2013). Cortical interneurons that specialize in disinhibitory control. Nature 503, 521–524. 10.1038/nature12676.

34. Fu, Y., Kaneko, M., Tang, Y., Alvarez-Buylla, A., and Stryker, M.P. (2015). A cortical disinhibitory circuit for enhancing adult plasticity. Elife 4, e05558. 10.7554/eLife.05558.

35. 35. de Brito Van Velze, M., Dhanasobhon, D., Martinez, M., Morabito, A., Berthaux, E., Pinho, C.M., Zerlaut, Y., and Rebola, N. (2024). Feedforward and disinhibitory circuits differentially control activity of cortical somatostatin interneurons during behavioral state transitions. Cell Rep 43, 114197. 10.1016/j.celrep.2024.114197.

36. Ferguson, K.A., Salameh, J., Alba, C., Selwyn, H., Barnes, C., Lohani, S., and Cardin, J.A. (2023). VIP interneurons regulate cortical size tuning and visual perception. Cell Rep 42, 113088. 10.1016/j.celrep.2023.113088.

37. Hartung, J., Schroeder, A., Péréz Vázquez, R.A., Poorthuis, R.B., and Letzkus, J.J. (2024). Layer 1 NDNF interneurons are specialized top-down master regulators of cortical circuits. Cell Rep 43. 10.1016/j.celrep.2024.114212.

38. Schuman, B., Machold, R.P., Hashikawa, Y., Fuzik, J., Fishell, G.J., and Rudy, B. (2019). Four unique interneuron populations reside in neocortical layer 1. Journal of Neuroscience 39. 10.1523/JNEUROSCI.1613-18.2018.

39. Munoz, W., and Rudy, B. (2014). Spatiotemporal specificity in cholinergic control of neocortical function. Curr Opin Neurobiol 26, 149–160. 10.1016/j.conb.2014.02.015.

40. Bennett, B.D., Callaway, J.C., and Wilson, C.J. (2000). Intrinsic membrane properties underlying spontaneous tonic firing in neostriatal cholinergic interneurons. J Neurosci 20, 8493–8503. 10.1523/JNEUROSCI.20-22-08493.2000.

41. Kawaguchi, Y. (1993). Physiological, morphological, and histochemical characterization of three classes of interneurons in rat neostriatum. J Neurosci 13, 4908–4923. 10.1523/JNEUROSCI.13-11-04908.1993.

42. Granger, A.J., Wang, W., Robertson, K., El-Rifai, M., Zanello, A.F., Bistrong, K., Saunders, A., Chow, B.W., Nuñez, V., García, M.T., et al. (2020). Cortical ChAT+ neurons co-transmit acetylcholine and GABA in a target-and brain-region-specific manner. Elife 9. 10.7554/eLife.57749.

43. Abs, E., Poorthuis, R.B., Apelblat, D., Muhammad, K., Pardi, M.B., Enke, L., Kushinsky, D., Pu, D.L., Eizinger, M.F., Conzelmann, K.K., et al. (2018). Learning-Related Plasticity in Dendrite-Targeting Layer 1 Interneurons. Neuron 100, 684–699 e6. 10.1016/j.neuron.2018.09.001.

44. Letzkus, J.J., Wolff, S.B., Meyer, E.M., Tovote, P., Courtin, J., Herry, C., and Luthi, A. (2011). A disinhibitory microcircuit for associative fear learning in the auditory cortex. Nature 480, 331–335. 10.1038/nature10674.

45. Takesian, A.E., Bogart, L.J., Lichtman, J.W., and Hensch, T.K. (2018). Inhibitory circuit gating of auditory critical-period plasticity. Nat Neurosci 21, 218–227. 10.1038/s41593-017-0064-2.

46. Povysheva, N. V., Zaitsev, A. V., Kröner, S., Krimer, O.A., Rotaru, D.C., Gonzalez-Burgos, G., Lewis, D.A., and Krimer, L.S. (2007). Electrophysiological differences between neurogliaform cells from monkey and rat prefrontal cortex. J Neurophysiol 97. 10.1152/jn.00794.2006.

47. Chittajallu, R., Auville, K., Mahadevan, V., Lai, M., Hunt, S., Calvigioni, D., Pelkey, K.A., Zaghloul, K.A., and McBain, C.J. (2020). Activity-dependent tuning of intrinsic excitability in mouse and human neurogliaform cells. Elife 9. 10.7554/eLife.57571.

48. Reha, R.K., Dias, B.G., Nelson, C.A., Kaufer, D., Werker, J.F., Kolbh, B., Levine, J.D., and Hensch, T.K. (2020). Critical period regulation acrossmultiple timescales. Preprint, 10.1073/pnas.1820836117 https://doi.org/10.1073/pnas.1820836117.

49. Hensch, T.K. (2005). Critical period plasticity in local cortical circuits. Preprint, 10.1038/nrn1787 https://doi.org/10.1038/nrn1787.

50. Sohal, V.S., and Rubenstein, J.L.R. (2019). Excitation-inhibition balance as a framework for investigating mechanisms in neuropsychiatric disorders. Mol Psychiatry 24. 10.1038/s41380-019-0426-0.

51. Pouchelon, G., Dwivedi, D., Bollmann, Y., Agba, C.K., Xu, Q., Mirow, A.M.C., Kim, S., Qiu, Y., Sevier, E., Ritola, K.D., et al. (2021). The organization and development of cortical interneuron presynaptic circuits are area specific. Cell Rep 37. 10.1016/j.celrep.2021.109993.

52. Favuzzi, E., Deogracias, R., Marques-Smith, A., Maeso, P., Jezequel, J., Exposito-Alonso, D., Balia, M., Kroon, T., Hinojosa, A.J., Maraver, E.F., et al. (2019). Neurodevelopment: Distinct molecular programs regulate synapse specificity in cortical inhibitory circuits. Science (1979) 363. 10.1126/science.aau8977.

53. Brown, D., Altermatt, M., Dobreva, T., Chen, S., Wang, A., Thomson, M., and Gradinaru, V. (2021). Deep Parallel Characterization of AAV Tropism and AAV-Mediated Transcriptional Changes via Single-Cell RNA Sequencing. Front Immunol 12, 730825. 10.3389/fimmu.2021.730825.

54. Jang, M.J., Coughlin, G.M., Jackson, C.R., Chen, X., Chuapoco, M.R., Vendemiatti, J.L., Wang, A.Z., and Gradinaru, V. (2023). Spatial transcriptomics for profiling the tropism of viral vectors in tissues. Nat Biotechnol 41, 1272–1286. 10.1038/s41587-022-01648-w.

55. Wang, D., Tai, P.W.L., and Gao, G. (2019). Adeno-associated virus vector as a platform for gene therapy delivery. Nat Rev Drug Discov 18, 358–378. 10.1038/s41573-019-0012-9.

56. Allaway, K.C., Gabitto, M.I., Wapinski, O., Saldi, G., Wang, C.Y., Bandler, R.C., Wu, S.J., Bonneau, R., and Fishell, G. (2021). Genetic and epigenetic coordination of cortical interneuron development. Nature 597, 693–697. 10.1038/s41586-021-03933-1.

57. Hubisz, M.J., Pollard, K.S., and Siepel, A. (2011). PHAST and RPHAST: phylogenetic analysis with space/time models. Brief Bioinform 12, 41–51. 10.1093/bib/bbq072.

58. Siepel, A., Bejerano, G., Pedersen, J.S., Hinrichs, A.S., Hou, M., Rosenbloom, K., Clawson, H., Spieth, J., Hillier, L.D.W., Richards, S., et al. (2005). Evolutionarily conserved elements in vertebrate, insect, worm, and yeast genomes. Genome Res 15. 10.1101/gr.3715005.

59. Challis, R.C., Ravindra Kumar, S., Chan, K.Y., Challis, C., Beadle, K., Jang, M.J., Kim, H.M., Rajendran, P.S., Tompkins, J.D., Shivkumar, K., et al. (2019). Systemic AAV vectors for widespread and targeted gene delivery in rodents. Nat Protoc 14, 379–414. 10.1038/s41596-018-0097-3.

60. Stuart, T., Srivastava, A., Madad, S., Lareau, C.A., and Satija, R. (2021). Single-cell chromatin state analysis with Signac. Nat Methods 18, 1333–1341. 10.1038/s41592-021-01282-5.

61. Susaki, E.A., Tainaka, K., Perrin, D., Yukinaga, H., Kuno, A., and Ueda, H.R. (2015). Advanced CUBIC protocols for whole-brain and whole-body clearing and imaging. Nat Protoc 10, 1709–1727. 10.1038/nprot.2015.085.

62. 62. Fredericks, J.M., Dash, K.E., Jaskot, E.M., Bennett, T.W., Lerchner, W., Dold, G., Ide, D., Cummins, A.C., Der Minassian, V.H., Turchi, J.N., et al. (2020). Methods for mechanical delivery of viral vectors into rhesus monkey brain. J Neurosci Methods 339, 108730. 10.1016/j.jneumeth.2020.108730.

63. Park, T.I., Smyth, L.C.D., Aalderink, M., Woolf, Z.R., Rustenhoven, J., Lee, K., Jansson, D., Smith, A., Feng, S., Correia, J., et al. (2022). Routine culture and study of adult human brain cells from neurosurgical specimens. Nat Protoc 17, 190–221. 10.1038/s41596-021-00637-8.

64. Schindelin, J., Arganda-Carreras, I., Frise, E., Kaynig, V., Longair, M., Pietzsch, T., Preibisch, S., Rueden, C., Saalfeld, S., Schmid, B., et al. (2012). Fiji: an open-source platform for biological-image analysis. Nat Methods 9, 676–682. 10.1038/nmeth.2019.

